# Distinct Mechanisms of Recognition of Phosphorylated RNAPII C-Terminal Domain by BRCT Repeats of the BRCA1–BARD1 Complex: Insights from Structural and Functional Analyses

**DOI:** 10.1101/2025.01.22.634233

**Authors:** V. Klapstova, K. Sedova, J. Houser, M. Sebesta

**Affiliations:** CEITEC–Central European Institute of Technology, Masaryk University; Brno, Czechia; National Centre for Biomolecular Research, Faculty of Science, Masaryk University; Brno, Czechia

**Keywords:** BRCA1, BRCT repeats, DNA repair, transcription, RNA polymerase II, condensation

## Abstract

Transcription competes with other DNA-dependent processes, such as DNA repair, for access to its substrate, DNA. However, the principles governing the interplay between these processes remain poorly understood. Evidence suggests that the BRCA1-BARD1 complex, a key player in the DNA damage response, may act as a mediator of this crosstalk. In this study, we investigated the molecular mechanism underpinning the interaction between RNA polymerase II (RNAPII) and the BRCA1-BARD1 complex, as well as its functional implications. Our findings reveal that the BRCT repeat of BRCA1 binds the Ser5-phosphorylated CTD of RNAPII, utilising a mechanism previously established for other BRCT ligands. Furthermore, we demonstrate that this interaction is critical for the organisation of RNAPII into condensates with liquid-like properties. Analysis of disease-associated variants within the BRCT repeats further supports the biological relevance of this condensation. Collectively, our results suggest that the BRCA1-BARD1 complex may coordinate transcription and DNA repair by facilitating the organisation of RNAPII into transcription factories.

## Introduction

Accurate transmission of genetic information is paramount for cellular viability. Since DNA functions as a template for replication, recombination, repair, and transcription simultaneously, these processes can intersect, potentially leading to genomic instability. ^1–4^ One of the major sources of conflict arises from the formation of R-loops, tripartite nucleic-acid structures in which the nascent RNA hybridises with its complementary DNA strand, leaving the non-template strand unpaired. R-loops can tether RNA polymerases (RNAPs) to chromatin, thereby increasing the risk of collisions with replication and repair complexes. ^5–8^ To mitigate these transcription–replication/repair conflicts, cells use nucleases (*e.g.*, RNase H) and helicases (*e.g.*, senataxin and members of the DDX family) to dismantle R-loops ^9–16^ and factors such as XRN2 and senataxin to promote the dislodging of RNAP from chromatin. ^17,18^

Although transcription can threaten genome stability by generating R-loops, it also plays a beneficial role in DNA repair. For instance, RNA polymerase II (RNAPII) activity is critical for efficient repair of DNA double-stranded breaks (DSBs) via homologous recombination (HR). ^19–23^ Current models propose that RNAPII produces damage-specific short RNAs (DARTs) at DSB sites to activate the DNA damage response, thereby promoting repair. ^24,25^ The kinase c-ABL phosphorylates the C-terminal domain (CTD) of the catalytic subunit of RNAPII to facilitate this damage-induced transcription, ^26,27^ and once the RNAPII activity is no longer required, the phosphatase PP2A and senataxin helicase are recruited, via INTS6, to remove RNAPII and enable efficient DSB repair. ^28^

The presence of RNAPII at DSBs raises the question of how its activity is coordinated with the early steps of HR. A likely candidate for mediating this crosstalk is the BRCA1–BARD1 complex, known to associate with transcriptionally engaged RNAPII ^29–33^ – possibly via its BRCT repeats, which recognise phosphoproteins. ^33,34^ The BRCA1-BARD1 complex is essential for HR, in which it promotes pathway choice (by antagonizing the 53BP1-mediated non-homologous end-joining pathway) and stimulates RAD51 filament formation required for homology search. ^35–40^ It also ubiquitinates chromatin to activate the DNA damage response ^41,42^ and modulates DNA end processing enzymes, placing the BRCA1-BARD1 complex in a prime position to promote the crosstalk between transcription and HR.

Recent studies suggest that both transcription factors ^43–52^ and DNA repair factors ^24,53–58^ can assemble into membraneless factories, potentially via liquid-liquid phase separation (LLPS). Such clustering may enhance the efficiency and spatial organisation of nuclear processes.

Nevertheless, how specificity and selectivity in these condensates is achieved remains poorly understood, particularly regarding the interplay between transcription and HR machinery.

Here, we provide the first detailed investigation of direct interactions between RNAPII and the BRCA1–BARD1 complex at the molecular level. Using biochemical and structural analyses, we show that the tandem BRCT repeats of BRCA1 and BARD1 differ in their binding for the CTD of RNAPII. We further present the first three-dimensional structure of a tandem BRCT domain bound to a pSer5-CTD peptide, revealing how this binding promotes pCTD condensation in an interaction-dependent manner. Finally, we demonstrate that disease-associated variants within these BRCT repeats impair LLPS *in vitro* and correlate with cellular defects, underscoring the functional relevance of the direct interaction of BRCA1–BARD1 and RNAPII. These findings provide new insights into how transcription is coordinated with early HR events.

## Results

### The BRCA1-BARD1 complex interacts with the RNAPII CTD through its BRCT repeats

Ample evidence supports the association of the BRCA1-BARD1 complex with RNAPII bound to chromatin. ^29,30,59,60^ However, the possibility of a direct, physical interaction between these complexes has not been investigated.

To address this, we purified the BRCA1-BARD1 complex to near-homogeneity from insect cells using a combination of affinity chromatography and size-exclusion chromatography (Fig. 1A, Supplementary Fig. 1A). We first tested whether the BRCA1-BARD1 complex, immobilised on FLAG beads, could pull down RNAPII from HEK293 cell extract. The results indicate that the BRCA1-BARD1 complex pulled-down RNAPII with a CTD primarily phosphorylated on serine 5 (pS5-CTD) (Fig. 1B, Supplementary Fig. 2). Next, we tested the ability of purified GST-(CTD)_26_ peptides – specifically phosphorylated on Y1, S2, and S5 by ABL1, DYRK1a, and the CDK7 complex, respectively^57^ – to pull down BRCA1-BARD1. *In vitro*, the BRCA1-BARD1 complex directly, and specifically, recognises CTD phosphorylated on S2 and S5 (Fig. 1B). Further analysis showed that this interaction is mediated by the BRCT domains, via the conserved residues S1655 and K1702 in BRCA1, and S575 and K619 in BARD1, which have been implicated in phospho-serine recognition ^33,34,61–63^ (Fig. 1D, Supplementary Fig. 3).

**Figure 1:**
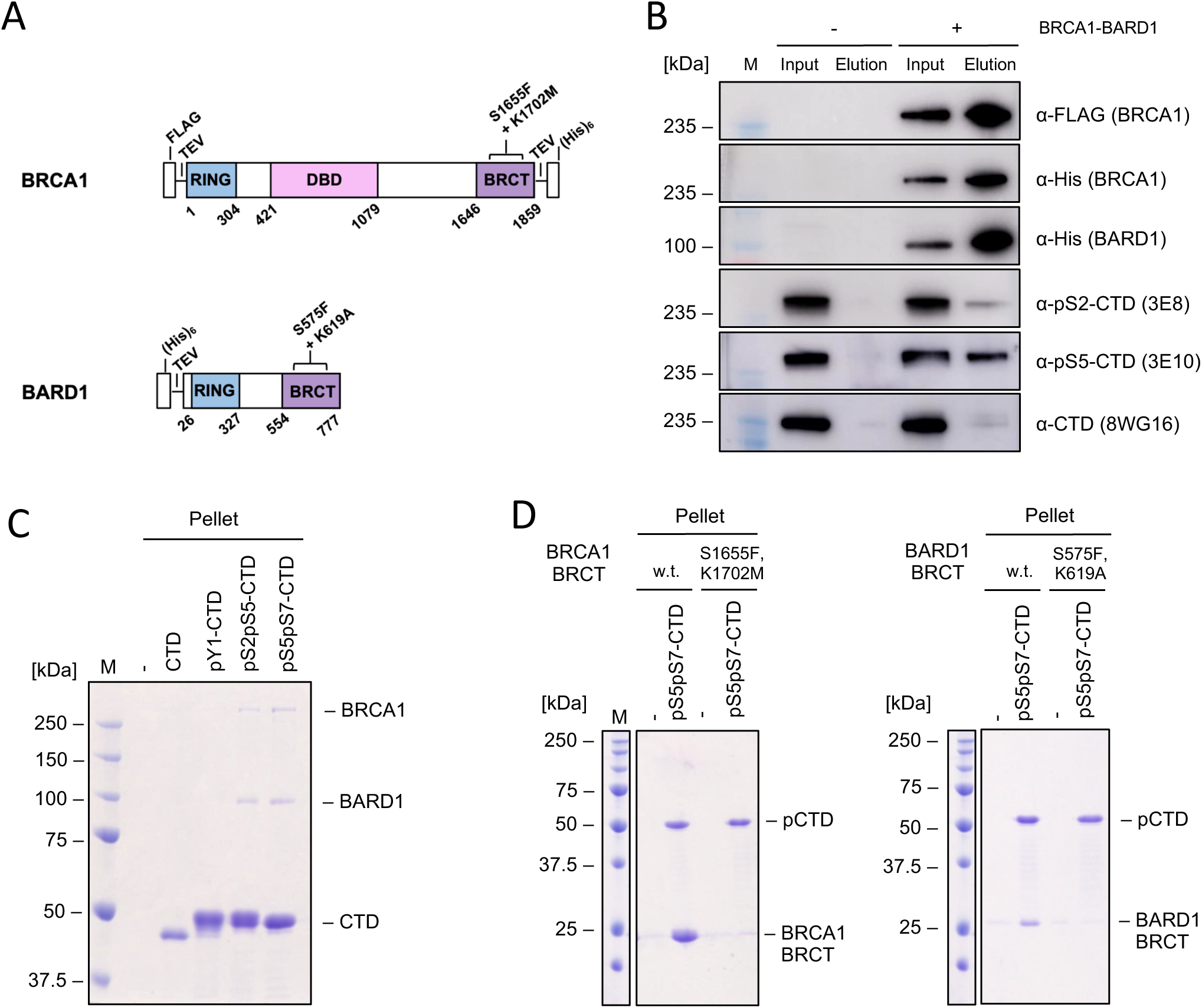
The interaction between the BRCA1-BARD1 complex and RNAPII is direct and mediated by the phosphorylated CTD of RNAPII and the BRCT domains of BRCA1-BARD1. A) Schematic representation of BRCA1 and BARD1 domains. Positions of investigated binding variants, the tags, and cleavage sites are indicated. B) BRCA1-BARD1 interacts with RNAPII via the CTD phosphorylated on Ser2 and Ser5. Western blot analysis of pull-downs from HEK293 lysates. HEK293 cells were lysed, and the lysate was cleared by centrifugation. To the supernatant, FLAG-BRCA1-BARD1 was added, and the samples were incubated with α-FLAG beads. As a control, the HEK293 lysate with no added BRCA1-BARD1 was used. The proteins were eluted using 3xFLAG peptide and the samples were analysed using western blots. C) BRCA1-BARD1 interacts directly with pS2pS5 GST-(CTD)_26_ and pS5pS7 GST-(CTD)_26_ *in vitro*. SDS-PAGE analysis of *in vitro* pull-down assay between GST-(CTD)_26_ and BRCA1-BARD1. Purified BRCA1-BARD1 was incubated with phosphorylated and non-phosphorylated GST-(CTD)_26_ bound to glutathione beads. The samples were centrifuged and the input, unbound (supernatant) and bound (pellet) fractions were analysed using the SDS PAGE. D) Substitutions in the phosphoserine binding site (BRCA1^S1655F, K1702M^, BARD1^S575F, K619A^) abolish the binding. SDS-PAGE analysis of *in* vitro pull-down assay between GST-(CTD)_26_ and BRCA1 BRCT and BARD1 BRCT, respectively. Purified BRCA1 BRCT and BARD1 BRCT, respectively, were incubated with phosphorylated and non-phosphorylated GST-(CTD)_26_ bound to glutathione beads. The samples were centrifuged and the input, unbound (supernatant) and bound (pellet) fractions were analysed using the SDS PAGE.

In summary, these data indicate that the BRCA1-BARD1 complex directly interacts with the phosphorylated CTD (pCTD) of RNAPII. This interaction is mediated by the BRCT repeats on both subunits, and the mechanism of phospho-serine recognition appears similar to that proposed for other known BRCT ligands.

### The BRCT repeats differ in binding kinetics to the phospho-CTD

In our initial *in vitro* binding assay, we observed a clear difference in the amount of pulled-down BRCT repeats (Fig. 1D). To investigate this difference, we measured the binding kinetics between the isolated BRCT repeats and pCTD using bio-layer interferometry (BLI). The data revealed an apparent affinity of 0.02 µM for BRCA1 BRCT and 0.193 µM for BARD1 BRCT (Fig. 2D, Supplementary Fig. 6A). Control experiments showed that the variants of BRCT domains defective in recognition of phosphopeptide (BRCA1 BRCT^S1655F,K1702M^ and BARD1 BRCT^S575F,K619A^) did not bind pCTD, nor did the wild-type BRCT domains bind unmodified CTD, consistent with the *in vitro* pull-down assays described above.

**Figure 2:**
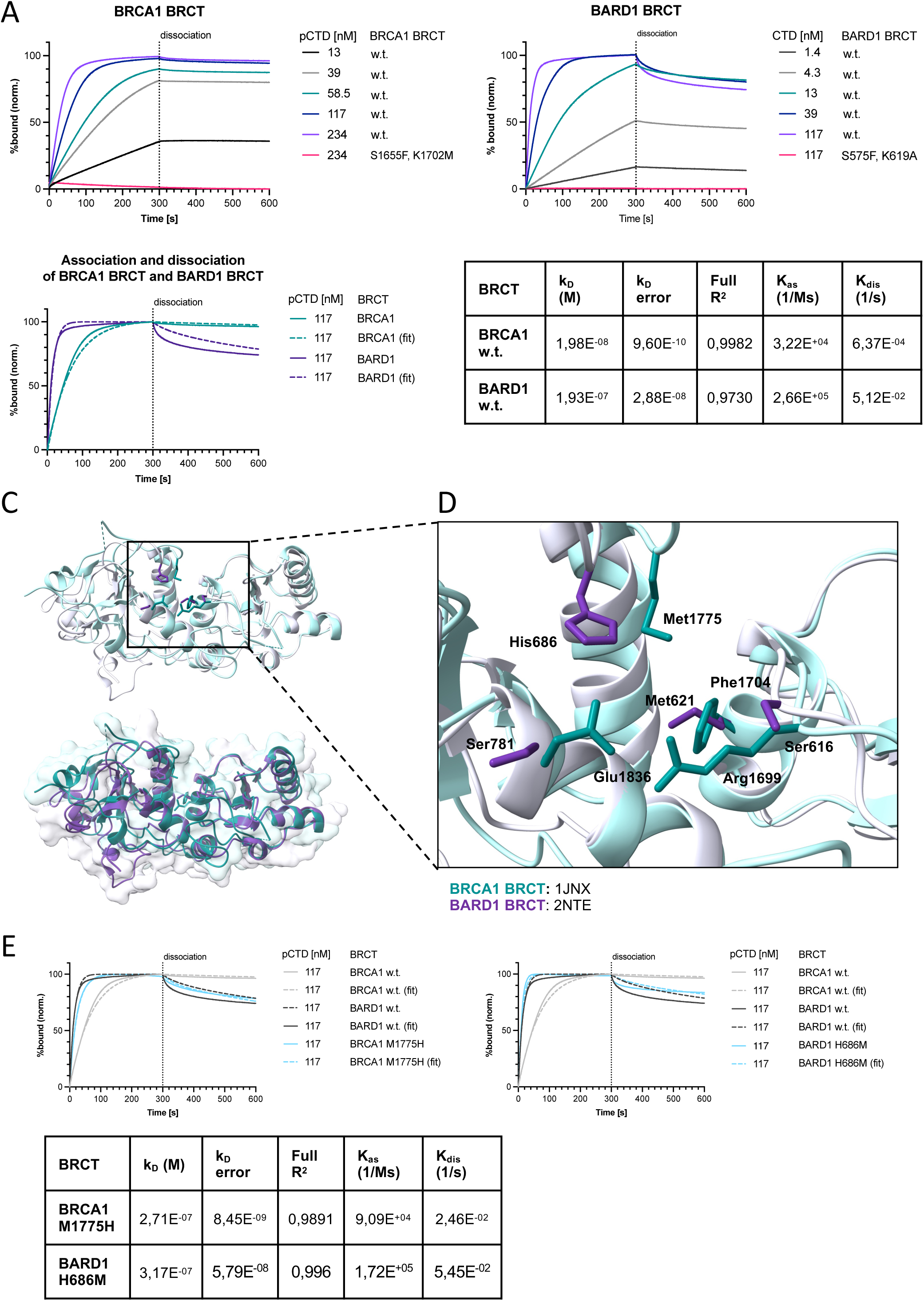
The BRCT domains of the BRCA1–BARD1 complex exhibit differences in their binding kinetics to the phosphorylated CTD of RNAPII. A) Substitutions in the phosphoserine binding site of the BRCT repeats (BRCA1^S1655F, K1702M^, BARD1^S575F, K619A^) abolish the binding to phosphorylated CTD. Sensograms obtained by biolayer-interferometry (BLI). The sensograms represent the mean of 3 measurements for each concentration. The data were analysed in Octet® Analysis Studio Software using 1:2 Bivalent analyte model. The data were plotted using Prism GraphPad 9 software. B) BARD1 BRCT associates with GST-p(CTD)_26_ more dynamically than BRCA1 BRCT. Sensograms obtained by biolayer-interferometry (BLI) and their respective fits (left). Comparison of association (*K*_as_), dissociation (*K*_dis_), and dissociation (*K*_D_) constants for pS5pS7 GST-(CTD)_26_ and BRCA1 and BARD1 BRCT, respectively, obtained by biolayer interferometry (right). The sensograms represent the mean of 3 measurements for each concentration. The association and dissociation constants and the coefficient of determination (R2) indicating the appropriateness of the fit were calculated in Octet® Analysis Studio Software using 1:2 Bivalent analyte model. The data were plotted using Prism GraphPad 9 software. C) Structural alignment of BRCA1 BRCT (PDB: 1JNX, teal) and BARD1 BRCT (PDB: 2NTE, purple) obtained in UCSF Chimera. D) Comparison of the amino-acid composition of the hydrophobic pocket of BRCA1 BRCT (1JNX, teal) and the residues present on the homologous positions in BARD1 BRCT (2NTE, purple). Close up from (C). E) Comparison of sensograms obtained by biolayer-interferometry (BLI) and their respective fits (top) of pS5pS7 GST-(CTD)_26_ binding to BRCA1 BRCT^M1775H^ and BARD1 BRCT^H686H^. Kinetic parameters (association (*K*_as_), dissociation (*K*_dis_), and dissociation (*K*_D_) constants) (bottom). The sensograms represent the mean of 3 measurements for each concentration. The data were analysed in Octet® Analysis Studio Software using 1:2 Bivalent analyte model. The data were plotted using Prism GraphPad 9 software.

The binding profile suggests that the BARD1 BRCT exhibits more dynamic binding, as it associates and dissociates faster from the hyperphosphorylated CTD (Fig. 2B). To investigate this phenomenon further, we aligned the existing apo structures of the BRCT domains (PDB codes: 1JNX, 2NTE) ^64,65^ (Fig. 2C,D, Supplementary Fig. 6B). This structural alignment revealed that the BARD1 hydrophobic pocket is shallower – due to the presence of histidine 686 (H686) instead of methionine 1775 (M1775) in BRCA1 – thereby potentially affecting the accommodation of Y1 residue of the CTD. To test this hypothesis, we engineered BRCA1 BRCT^M1775H^ and BARD1 BRCT^H686M^ to swap these key residues (Fig. 2E). BRCA1 BRCT^M1775H^ exhibited binding kinetics similar to BARD1 BRCT, supporting our model. However, the mirror variant (BARD1 BRCT^H686M^) retained a more dynamic binding profile, suggesting that additional substitutions are necessary to reconstitute a fully BRCA1-like hydrophobic pocket.

In summary, the kinetic binding assays show that the two BRCT domains within the BRCA1-BARD1 complex differ in recognising the Y1 of the pCTD.

### The mechanism of BRCT repeat binding to a pSer5-CTD diheptad

The specific binding of the BRCA1 and BARD1 BRCT repeats towards the hyperphosphorylated CTD prompted structural characterisation of the binding via X-ray crystallography. Mixtures of BRCT repeats and peptides derived from the consensus CTD sequence – modified specifically on either S5 alone or on S2, S5, and S7 – were used. We obtained crystal structures of the ligand-free BRCT repeats for both BRCA1 and BARD1, diffracting at 2.4 Å and 1.8 Å, respectively. These apo-structures were nearly identical to existing ones; therefore, we did not analyse them further.

No additional density corresponding to the CTD-derived peptide was observed in crystals formed by the BARD1 BRCT domain when mixed with either pS5-CTD or pS2pS5pS7-CTD. In contrast, clear density corresponding to the pS5-CTD-derived peptide was observed in the BRCA1 BRCT dataset, despite the BRCA1 BRCT domain exhibiting similar affinities for both peptides (Supplementary Fig. 7A). The BRCA1 BRCT-CTD complex crystallised in the C2221 space group with four monomers in the asymmetric unit, and the structure was refined to 2.8 Å resolution (Supplementary Table 4). Eight of the 14 residues of the pS5-CTD peptide could be fitted into the extra density (Fig. 3A; Supplementary Fig. 7B-E). Structural analysis revealed that the CTD ligand was accommodated within the positively charged binding pocket of the BRCA1 BRCT domain without inducing conformational changes to the overall domain fold. Indeed, structural alignment of the apo and liganded forms indicated no induced-fit movements (Supplementary Fig. 7).

**Figure 3:**
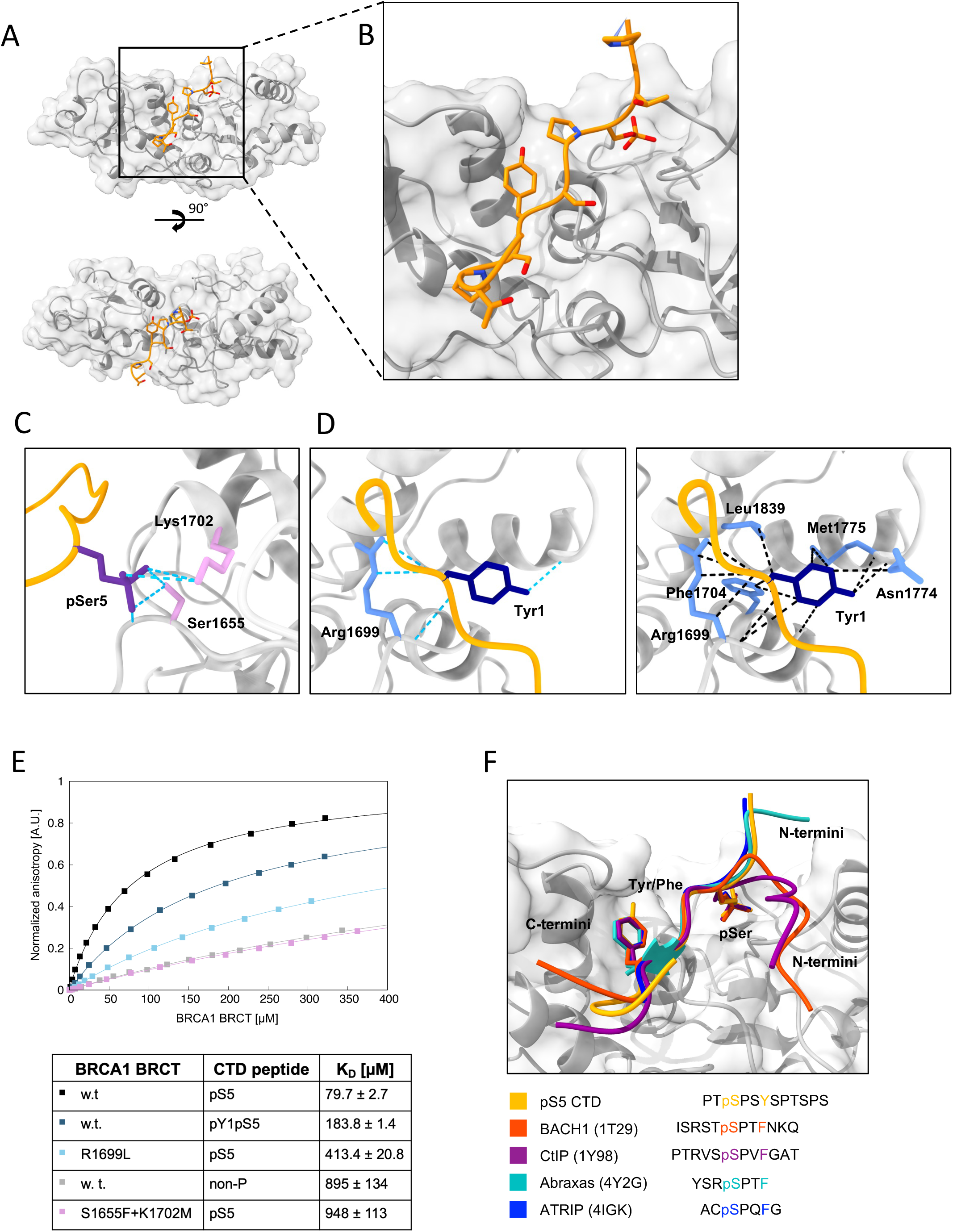
Structural characterisation of the BRCA1 BRCT domain bound to pS5 CTD peptide. **A)** Crystal structure of BRCA1 BRCT (gray) with bound pS5 CTD peptide (yellow). **B)** Detail of the BRCA1 BRCT phospho-peptide binding site (gray) with bound pS5 CTD peptide (yellow). **C)** Detail of the phosphoserine binding site of BRCA1 BRCT (gray) with bound pS5 CTD peptide (yellow). The pS5 of the CTD peptide is depicted in purple, the amino-acid residues interacting with pS5 are depicted in light violet. The hydrogen bonds were displayed as pseudo bonds using the structural analysis tool Hydrogen bonds in UCSF ChimeraX and are depicted in turquoise. The relax distance tolerance was 1.0 Å and the relax angle tolerance was 20.0°. **D)** Detail of the aromatic amino-acid binding pocket of BRCA1 BRCT (gray) with bound pS5 CTD peptide (yellow). The Y1 of the CTD is depicted in navy, the interacting amino-acid residues in light blue. The hydrogen bonds (left) were displayed as pseudobonds using the structural analysis tool Hydrogen bonds in UCSF ChimeraX and are depicted in turquoise. The relax distance tolerance was 1.0 Å and the relax angle tolerance was 20.0°. Van der Waals, hydrophobic and stacking interactions (right) were displayed as pseudobonds using the structural analysis tool in UCSF ChimeraX and are depicted in black. The interacting atoms were indentified based on VDW overlap ≥-0.4 Å. **E)** Validation of the obtained structural model by fluorescence anisotropy measurement. The assays were performed between BRCA1 BRCT domain and the indicated ligands (at 25nM). Anisotropy data were plotted as a function of protein concentration and fitted to a single-site saturation with non-specific binding model using XMGrace. **F)** Comparison of binding of different ligands to BRCA1 BRCT. Alignment of the structures was created using UCSF ChimeraX. BRCA1 BRCT (from the structure with pCTD) is depicted in gray, pCTD peptide in yellow, BACH1 phosphopeptide (PDB: 1T29) in red, CtIP phosphopeptide (PDB: 1Y98) in magenta, Abraxas singly phosphorylated peptide (PDB: 4Y2G) in turquoise, ATRIP phosphopeptide (PDB: 4IGH) in blue.

Closer examination revealed that the pS5 CTD ligand is bound via the canonical two-anchor mechanism, consistent with other known BRCA1 BRCT ligands (Fig. 3A,B). ^66–70^Specifically, pS5 from the first CTD repeat is recognised by S1655 and K1702, and the Y1 from the next CTD repeat occupies a hydrophobic pocket formed by R1699, F1704, N1774, M1775, and L1839. Moreover, the P6 residue from the first CTD repeat contributes to this interaction, aligning with the canonical pSPXY/F binding motif seen in other BRCA1 BRCT ligands (Fig. 3C-E). ^33^

We corroborated these X-ray data by mutational analysis using fluorescence anisotropy (FA) measurements with a pS5-(CTD)_2_ peptide. Replacing S1655 and K1702 reduced BRCA1 BRCT affinity to 895 ± 134 µM, similar to the affinity of the wild-type domain for the non-phosphorylated pS5-(CTD)_2_ (K_D_ = 948 ± 113 µM). Comparable results were obtained for the BARD1 BRCT^M1775K^ variant, confirming that M1775 is essential for accommodating the aromatic residue at +3. Likewise, using a pY1pS5-(CTD)_2_ peptide or the R1699L BRCA1 variant (which disrupts the hydrophobic pocket) weakened binding (K_D_ = 183.8 ± 1.4 µM and 413.4 ± 20.8 µM, respectively) (Fig. 3F).

Because BARD1 BRCT co-crystallisation with pCTD was unsuccessful under these conditions, we used AlphaFold 3 ^71^ (Supplementary Fig. 8A,B) to predict how BARD1 BRCT might bind pS5-(CTD)_2_. The resulting model aligned well with our mutational data, confirming that S575 and K619 recognise pS5 in the CTD and that H686, S616, M621, P687, N690, and I741 interact with Y1 in the CTD (Supplementary Fig. 8C,D).

In summary, these structural analyses clarify why the BRCA1 BRCT domain exhibits a preference for S5-pCTD and suggest a similar canonical binding mode for BARD1 BRCT domain.

### The BRCA1-BARD1 complex undergoes condensation *in vitro* and can simultaneously incorporate hyperphosphorylated CTD and RNA transcript

Our next question concerned the functional significance of BRCA1-BARD1 interacting with hyperphosphorylated CTD. Recent work has shown that phosphorylated RNAPII can cluster into transcription-cycle-specific condensates with liquid-like properties *in vivo*, ^72^ and the hyperphosphorylated form may shuttle as a client.^49,73^ Additionally, the BRCA1-BARD1 complex has also been suggested to be present in condensates *in vivo*. ^74^ We therefore asked whether the purified BRCA1-BARD1 complex can undergo condensation *in vitro*.

To test this hypothesis, we labelled the BRCA1-BARD1 complex *in situ* with Alexa488. During the assay, we used a 1:20 ratio (labelled:unlabelled) complex, because *in situ* labelling had a negative effect on condensation propensity (Supplementary Fig. 9C). When the sample was visualised by fluorescent microscopy in presence of 10% dextran, regularly shaped droplets appeared at concentrations as low as 1.25 µM. With increasing concentration (up to 5 µM), these droplets grew in size but decreased in number, suggesting fusion events consistent with liquid-like behaviour. To determine which forces drive this condensation, we added two known inhibitors: hexane-1,6-diol (which disrupts hydrophobic interactions) ^75^ and ATP (which can weaken electrostatic interactions) ^76^. The presence of hexane-1,6-diol reduced droplet number and caused irregularly shaped objects, indicative of partial aggregation, whilst ATP (5 mM) substantially reduced droplet size and number (Fig. 4B). Therefore, the BRCA1- BARD1 complex undergoes condensation via liquid-liquid phase separation (LLPS) that likely relies on both hydrophobic and electrostatic forces.

**Figure 4:**
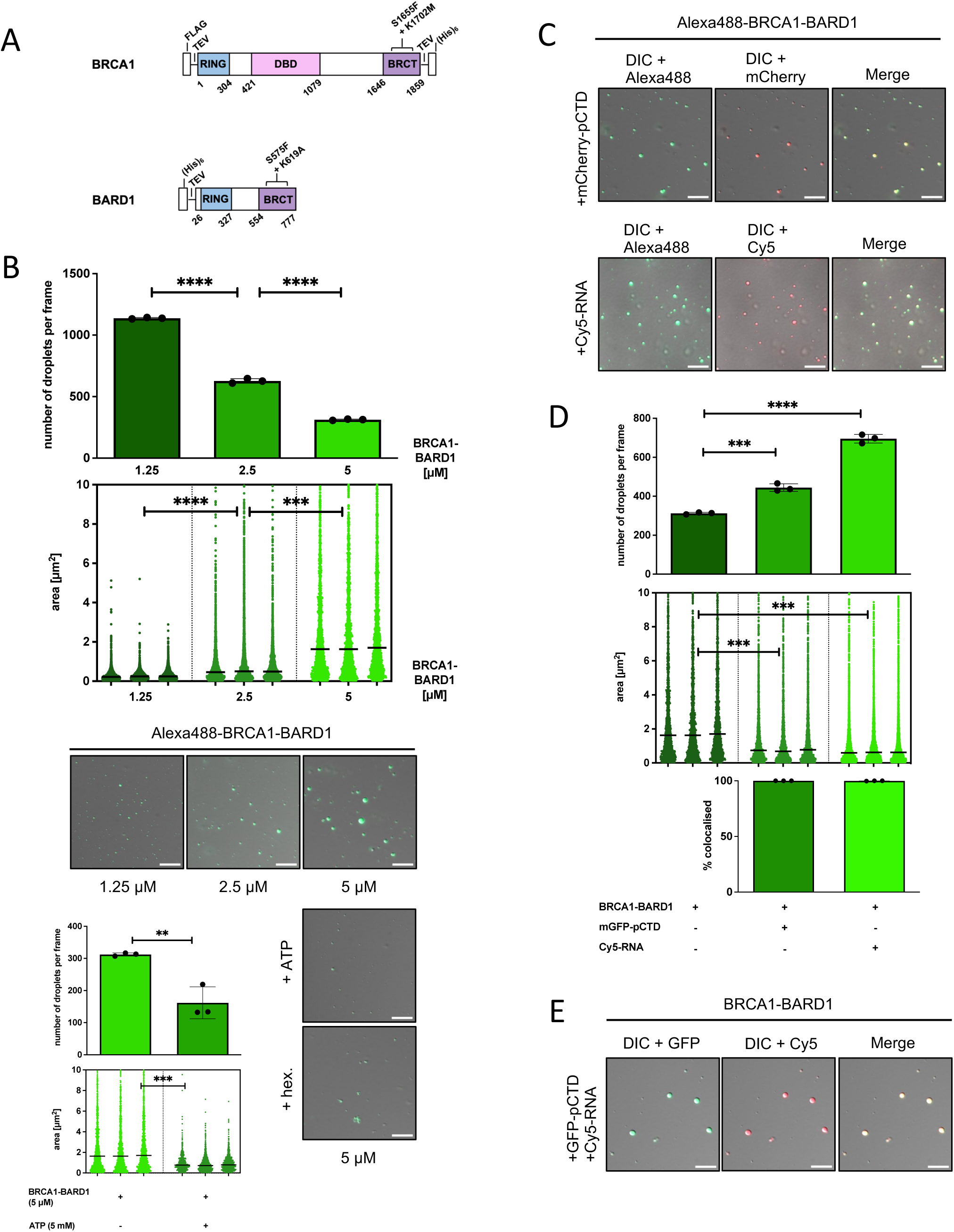
The BRCA1-BARD1 complex forms liquid-like condensates *in vitro*, which accommodate phosphorylated CTD domain of RNAPII and RNA. A) Schematic representation of BRCA1 and BARD1 domains. Positions of investigated binding mutants are indicated. B) Liquid-liquid phase separation (LLPS) assays with purified, Alexa488-labelled BRCA1- BARD1. BRCA1-BARD1 (at 1.25 µM, 2.5 µM, and 5 µM) was mixed with the crowding agent (10% dextran). Bar chart (top) representing quantification (n = 3) of the number of droplets per frame from the LLPS experiments with BRCA1-BARD1. Statistical significance was determined by unpaired t-test. A nested scatterplot (middle) represents quantification (n = 3) of an area of individual droplets from three independent experiments with BRCA1-BARD1, with median area determined per dataset. Statistical significance was determined by nested t- test. Representative images from three experiments (bottom) are depicted as an overlay of differential interference contrast (DIC) and GFP. Where indicated, hexane-1,6-diol (hex; at 10%) was added to inhibit hydrophobic interactions or ATP (at 5mM) to inhibit electrostatic interactions. Scale bars, 10 µm. C) LLPS assays with purified, Alexa488-labelled BRCA1-BARD1, pS5pS7 mCherry-hCTD and Cy5-RNA. BRCA1-BARD1 (at 5 µM) was mixed with phosphorylated CTD (2.5 µM) or Cy5-ITS1 RNA (at 15 nM) in the presence of a crowding agent (10% dextran). Representative images from three experiments are depicted as an overlay of differential interference contrast (DIC), Alexa488, and Cy5. Scale bars, 10 µm. D) Bar chart (top) representing quantification (n = 3) of the number of droplets per frame from the LLPS experiments with the BRCA1-BARD1 complex shown in (C). Statistical significance was determined by unpaired t-test. A nested scatterplot (bottom) represents quantification (n = 3) of an area of individual droplets from three independent experiments with the BRCA1- BARD1 complex shown in (C), with median area determined per dataset. Statistical significance was determined by nested t-test. E) LLPS assays with purified, BRCA1-BARD1, pS5pS7 GFP-(CTD)_26_ and Cy5-RNA. BRCA1-BARD1 (at 5 µM) was mixed with phosphorylated CTD (2.5 µM) and Cy5-ITS1 RNA (at 15 nM) in the presence of a crowding agent (10% dextran). Representative images from three experiments are depicted as an overlay of differential interference contrast (DIC), GFP, and Cy5. Scale bars, 10 µm.

Next, we investigated whether these BRCA1-BARD1 condensates can incorporate hyperphosphorylated CTD (mCherry-pCTD) (as a proxy for RNAPII *in vivo*) and a model ∼1 kb RNA transcript (Cy5-ITS1). Neither factor alone forms condensates. ^57,77^ To this end, we mixed the BRCA1-BARD1 with the said factors either individually or in combination and visualised the objects using fluorescent microscopy. Both pCTD and the RNA transcript were incorporated into condensates formed by the BRCA1-BARD1 complex (Fig. 4C,D). Nearly all assessed condensates contained signal for both the BRCA1-BARD1 complex and mCherry-pCTD. Similar results were observed when pCTD was substituted with the Cy5-labelled RNA transcript. Finally, we tested whether both pCTD and RNA can coexist in the same droplets. Indeed, we observed mGFP-pCTD and Cy5-ITS1 signals within droplets formed by unlabelled BRCA1-BARD1 under the tested conditions, confirming that the complex can simultaneously incorporate pCTD and RNA into the condensates.

In summary, our results demonstrate that BRCA1-BARD1 can form condensates capable of accommodating both pCTD (as a proxy for RNAPII) and RNA (as a proxy for cellular transcripts).

### The BRCT repeats of BRCA1 and BARD1 form condensates able to accommodate pCTD and RNA

Previous work suggested that condensation of the BRCA1-BARD1 complex *in vivo* may involve the BRCT domains in concert with specific transcripts^74^, implicating their active role in the process of condensation. We therefore asked whether the isolated BRCT domains could form droplets containing both RNA and pCTD.

Initially, we confirmed published reports that the BRCT repeats of both BRCA1 and BARD1 bind nucleic acids (DNA and RNA), ^78,79^ albeit to a lower extent than the specialised nucleic-acid-binding domain of BRCA1 (residues 421-1079). Importantly, the BRCT domain of BRCA1 bound RNA via the same site as the pCTD, whereas the BARD1 BRCT bound RNA via a different site, distinct from the canonical pCTD-binding pocket (Supplementary Fig. 11, 12A-C). This finding is further supported by: (i) competition EMSA, in which pCTD could compete for nucleic-acid binding to BRCA1 BRCT but not to BARD1 BRCT; and (ii) AlphaFold 3-generated models, in which an RNA molecule occupies the canonical phosphoserine-binding site in BRCA1 BRCT but a conserved basic patch near R705 in BARD1 BRCT (Supplementary Fig. 11D; 12D,E; 13).

Consistent with this, the BRCT repeats and RNA formed condensates in a concentration-dependent manner that required intact RNA binding (Supplementary Fig. 14A,B; 15).

Next, we tested whether the pCTD might also enter droplets formed by the BRCT domains of the BRCA1-BARD1 complex. We therefore mixed mCherry-pCTD with *in situ* fluorescently labelled BRCT repeats (Alexa488, at a 1:20 ratio labelled:unlabelled). Under stoichiometric conditions and 10% dextran, both BRCT repeats formed droplets with mCherry-pCTD that were significantly larger than condensates of BRCT alone (Fig. 5A,B). The co-localisation of Alexa488 and mCherry signals implies that both proteins entered the condensates, confirmed by a sedimentation assay in which both pCTD and BRCT repeats were found in the pellet (Supplementary Fig. 16B). Subsequently, we used unlabelled BRCT repeats to avoid labelling artefacts (Supplementary Fig. 14C,D).

**Figure 5:**
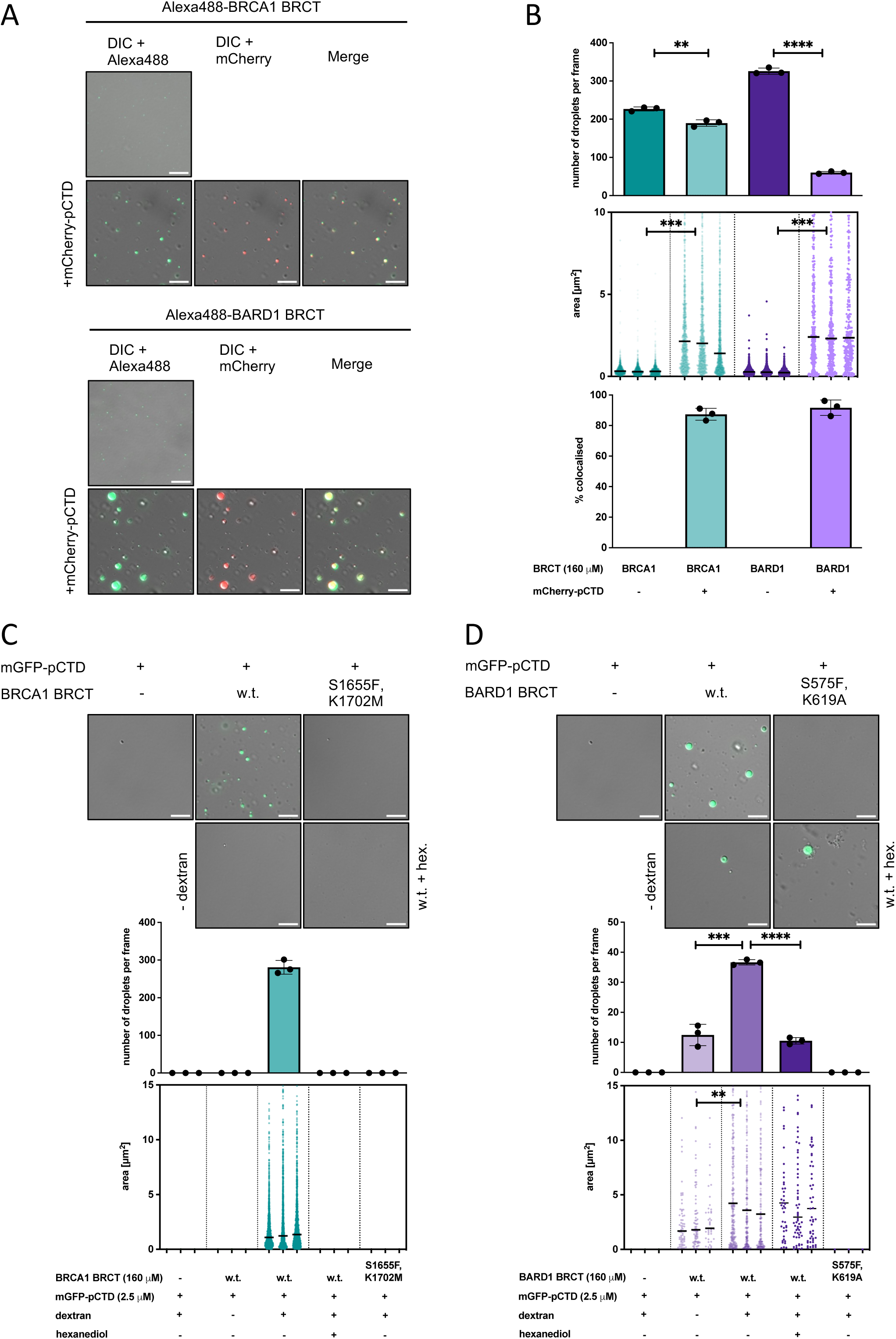
The BRCT repeats of the BRCA1-BARD1 form liquid-like condensates *in vitro*, which accommodate phosphorylated CTD domain of RNAPII and RNA. **A)** LLPS assays with pS5pS7 mCherry-hCTD and Alexa488-labelled BRCA1 BRCT (top) and BARD1 BRCT (bottom). The BRCT (at 160 µM) in absence or presence of phosphorylated CTD (2.5 µM) was mixed with the crowding agent (10% dextran). Representative images from three experiments are depicted as an overlay of differential interference contrast (DIC) with the signal for Alexa488 or mCherry. Scale bars 10µm. **B)** Bar chart (top) representing quantification (n = 3) of the number of droplets per frame from the LLPS experiments with BRCA1 and BARD1 BRCT shown in (A). Statistical significance was determined by unpaired t test. A nested scatterplot (middle) represents quantification (n = 3) of an area of individual droplets from three independent experiments with BRCA1-BARD1 shown in (A), with median area determined per dataset. Statistical significance was determined by nested t test. Bar chart (bottom) depicts the proportion of droplets containing signals for Alexa488-labelled BRCT (green) and pS5pS7 mCherry-hCTD (red). **C)** LLPS assays with purified BRCA1 BRCT and pS5pS7 mGFP-hCTD. BRCA1 BRCT (at 160 µM, w. t. and S1655F,K1702M, respectively) was mixed with phosphorylated CTD (2.5 µM) in the absence or presence of a crowding agent (10% dextran). Representative images (top) from three experiments are depicted as an overlay of differential interference contrast (DIC) and GFP. Hexane-1,6-diol (hex; at 10%) was added to inhibit hydrophobic interactions. Scale bars, 10 µm. Bar chart (middle) representing quantification (n = 3) of the number of droplets per frame from the LLPS experiments. Statistical significance was determined by unpaired t-test. A nested scatterplot (bottom) represents quantification (n = 3) of an area of individual droplets from three independent experiments, with median area determined per dataset. Statistical significance was determined by nested t-test. **D)** LLPS assays with purified BARD1 BRCT and pS5pS7 mGFP-hCTD. BARD1 BRCT (at 160 µM, w. t. and S575F,K619A) was mixed with phosphorylated CTD (2.5 µM) in the absence or presence of a crowding agent (10% dextran). Representative images (top) from three experiments are depicted as an overlay of differential interference contrast (DIC) and GFP. Hexane-1,6-diol (hex; at 10%) was added to inhibit hydrophobic interactions. Scale bars, 10 µm. Bar chart (middle) representing quantification (n = 3) of the number of droplets per frame from the LLPS experiments. Statistical significance was determined by unpaired t-test. A nested scatterplot (bottom) represents quantification (n = 3) of an area of individual droplets from three independent experiments, with median area determined per dataset. Statistical significance was determined by nested t-test.

When LLPS experiments were conducted with BRCA1 BRCT and pCTD-mGFP, droplet formation required both proteins, 10% crowding agent (Fig. 5C,D), and a direct physical interaction. These droplets exhibited liquid-like properties, as hexane-1,6-diol disrupted their formation. In contrast, BARD1 BRCT and mGFP-pCTD formed droplets even in the absence of a crowding agent, although less efficiently, and still required direct binding. Treatment with hexane-1,6-diol led to only partial disruption, suggesting that some droplets had transitioned to a more gel-like state. This may be explained by higher frequency of fusions of the droplets, which is supported by the presence of higher-order structures in the *in vitro* crosslinking experiments with the BARD1 BRCT repeat, compared to the BRCA1 BRCT repeat (Supplementary Fig. 16A).

These experiments support the view that BARD1 BRCT is more prone to condensation, possibly due to a higher propensity for oligomerisation in solution.

Finally, we examined whether both pCTD and RNA can be accommodated together in BRCT-domain droplets. We mixed BRCT domains with pCTD and RNA in the presence of a crowding agent and observed that both domains incorporated pCTD and RNA more efficiently in the presence of Mg^2+^. Consistent with the previously presented data, both pCTD and RNA required specific interactions with BRCT domains for efficient incorporation into condensates. However, the BRCA1 BRCT domain utilised a single interface to interact with both pCTD and RNA, whereas the BARD1 BRCT domain used two distinct interfaces for these interactions (Supplementary Fig. 17, 18).

In summary, our data show the BRCA1-BARD1 BRCT domains form condensates that can simultaneously incorporate pCTD and RNA. However, the domains differ significantly in the mechanism. Whilst BRCA1 BRCT uses an overlapping interface for pCTD and RNA, BARD1 BRCT uses separate, discrete binding sites.

### Characterisation of disease-associated variants of BRCT domains

The ability of the BRCT domains to form condensates provided a direct readout to investigate uncharacterised disease-associated variants of BRCA1 and BARD1 within their BRCT repeats. We focused on several conserved residues lying outside the phospho-serine-binding region (Fig. 6A, Supplementary Fig. 19).

**Figure 6:**
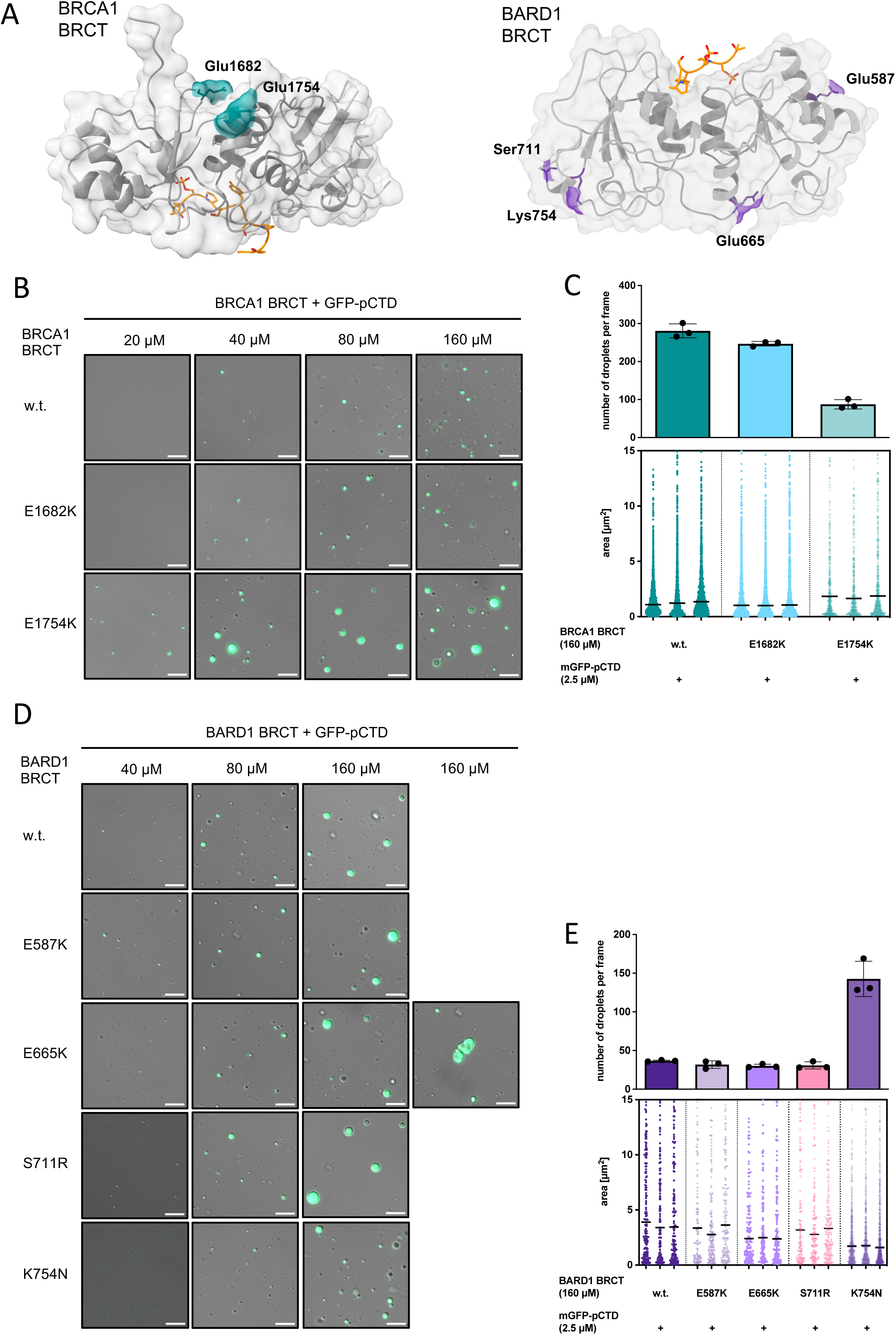
Characterisation of disease-associated variants within the BRCT repeats of the BRCA1-BARD1 complex on their ability to promote condensation *in vitro*. **B)** Positions of the investigated mutations of BRCT domains. The crystal structure of BRCA1 BRCT with the pS5 CTD ligand (left) and the AlphaFold 3-generated model of the BARD1 BRCT with the pS5 CTD ligand (right) were used. Investigated mutations are not located near the phospho-peptide binding sites and therefore should not interfere with the binding. **C)** LLPS assays with purified BRCA1 BRCT and pS5pS7 mGFP-hCTD. BRCA1 BRCT (at20 µM, 40 µM, 80 µM, and 160 µM), w. t., E1682K, and E1754K, was mixed with phosphorylated CTD (2.5 µM) in the presence of a crowding agent (10% dextran). Representative images from three experiments are depicted as an overlay of differential interference contrast (DIC) and GFP. Scale bars, 10 µm. **D)** Bar chart (top) representing quantification (n = 3) of the number of droplets per frame from the LLPS experiments with BRCA1 BRCT (w.t. and mutated variants, at 160 µM) and pS5pS7 mGFP-hCTD, shown in (B), using the green fluorescent signal. Statistical significance was determined by unpaired t test. A nested scatterplot (bottom) representing quantification (n = 3) of an area of individual droplets from three independent experiments with BRCA1 BRCT, pS5pS7 mGFP-hCTD, shown in (B), with median area determined per dataset. Statistical significance was determined by nested t test. **E)** LLPS assays with purified BARD1 BRCT and pS5pS7 mGFP-hCTD. BARD1 BRCT (at 40 µM, 80 µM, 160 µM), w.t., E587K, E665K, S711R, and K754N, respectively, was mixed with phosphorylated CTD (2.5 µM) in the presence of a crowding agent (10% dextran). Representative images from three experiments are depicted as an overlay of differential interference contrast (DIC) and GFP. Scale bars, 10 µm. **F)** Bar chart (top) representing quantification (n = 3) of the number of droplets per frame from the LLPS experiments with BARD1 BRCT (w.t. and mutated variants, at 160 µM), and pS5pS7 mGFP-hCTD, shown in (D), using the green fluorescent signal. The analysis and visualisation were performed as in (C).

We purified and analysed the following variants: BRCA1 BRCT^E1682K^, BRCA1 BRCT^E1754K^, and BARD1 BRCT^E587K^, BARD1 BRCT^E665K^, BARD1 BRCT^S711R^, and BARD1 BRCT^K754N^ (Supplementary Fig. 1A,B). These variants showed significantly lower thermal stability but retained wild-type affinity towards pCTD (Supplementary Fig. 19, 20).

When we tested the ability of these variants to form condensates in the presence of pCTD, BRCA1 BRCT^E1754K^ exhibited a significantly higher propensity for condensation (droplets visible already at 20 µM), whereas BRCA1 BRCT^E1682K^ did not show this effect (Fig. 6B; Supplementary Fig. 22A; 23A). Among the BARD1 variants, BRCT^K754N^ showed defective droplet fusion, producing numerous smaller droplets (Fig. 6C; Supplementary Fig. 22B). Even though variant BARD1 BRCT^E665K^ maintained wild type level of condensates, it exhibited higher propensity to aggregate as evidenced by the appearance of irregularly shaped objects.

Moreover, when we labelled this variant *in situ*, only aggregates were observed, unlike the wild-type variant (Fig6C, Supplementary Fig. 22B;23B).

Overall, we tested several previously uncharacterised disease-associated variants of BRCA1 and BARD1 that alter the condensate-forming properties of the BRCT domains, suggesting that defective condensation might contribute to tumorigenesis.

In summary, we provide a comprehensive mechanistic characterisation of how the BRCA1-BARD1 complex interacts with RNAPII, which may link transcription and DNA repair. Our data also offer new insight into a potential functional role for this interaction in heterotypic condensates containing the BRCA1-BARD1 complex, RNAPII, and nucleic acids.

## Discussion

Understanding the mechanisms underlying crosstalk between essential cellular processes that compete for DNA – replication, recombination, repair, and transcription – is critical for understanding how these processes are orchestrated in a way that avoids collisions and possible genome instability. ^1–4^ Moreover, emerging concepts in mesoscale organisation and compartmentalisation of these processes via condensation in the nucleus suggest that the current state of the art is far from complete. ^53,72^

In this study, we focused on a detailed, mechanistic characterisation of the interaction between the BRCA1-BARD1 complex and RNAPII, with the aim of improving our understanding of the crosstalk between transcription and DNA repair. To this end, we performed a comparative analysis of the BRCT domains found in both BRCA1 and BARD1. Furthermore, we solved the 3D structure of a complex between the BRCA1 BRCT domain and a phosphopeptide containing two repeats of the C-terminal domain of the catalytic subunit of RNAPII phosphorylated on serine 5 (pS5-pCTD). Lastly, we explored the functional consequences of this interaction, uncovering a possible role in the organisation of transcription and/or repair factories via condensation.

The association of BRCA1-BARD1 with ongoing transcription has been known since 1997. ^29^ The initial proposition that the complex might, through its E3 ubiquitin ligase activity (via the RING domains) ^60,80,81^ mediate ubiquitination of RNAPII and its subsequent removal from chromatin upon DNA damage has not been corroborated, as other E3 ubiquitin ligases were shown to mediate this process. ^82–84^ This leaves open the question of the functional significance of the interaction.

Presently, we do not know the precise context in which the BRCA1-BARD1 complex associates with RNAPII *in vivo*. Does the complex bind RNAPII that has been paused or stalled due to encountering a damaged template, or does it associate with RNAPII that has encountered a broken chromosome undergoing repair by recombination? Or does the interaction reflect the role of RNAPII in *de novo* RNA generation specifically at sites of double-stranded DNA breaks (DSBs) – where the BRCA1-BARD1 complex is also present? There is also the possibility that the interaction might be S-phase specific, reflecting the possibility that sites of transcription-replication conflicts might be the place where the complex associates with RNAPII. Moreover, it is not known whether this association is coupled with the E3 ubiquitin activity of the BRCA1-BARD1 complex, ^85,86^ whereby associating with RNAPII would bring the ubiquitin machinery to specific genomic loci. These are only a few unanswered questions.

We sought to help address these questions by providing a thorough, mechanistic characterisation of the interaction. Determining that the preferred substrate is pS5-pCTD helps narrow down the context in which the BRCA1-BARD1 complex may associate with RNAPII. This modification is associated with the early stages of the transcription cycle, ^87–89^ and there is supporting *in vivo* evidence for the complex to be associated with pS5-pCTD. ^30^

Moreover, our comparative analysis of the BRCT domains within the BRCA1-BARD1 complex revealed that they substantially differ in their mode of binding, despite both binding the RNAPII pCTD. The BRCA1 BRCT repeat stably associates with pCTD, whereas the BARD1 BRCT displays a more dynamic binding–likely because of the changes to hydrophobic pocket in BARD1 BRCT. This difference opens the possibility that the complex may use the BRCA1 BRCT repeat for stable association with transcribing RNAPII, whereas BARD1 could dynamically interact with the surroundings of RNAPII, especially given the flexibility and length (∼700 Å) of the CTD. ^90–92^

Additionally, the lack of structural data for the BARD1 BRCT domain bound to its cognate ligand might point to the possibility that phosphoproteins are not the best substrates for the BARD1 BRCT domain. Chemically diverse ligands, such as nucleosomes or (poly)ADP ribose, have been suggested as potential binding partners. ^37,93–95^

The presented crystal structure of the complex between the BRCA1 BRCT domain and a diheptad of CTD phosphorylated on serine 5 (pS5-(pCTD)_2_) is the first structure of a BRCT domain bound to the pCTD, expanding the repertoire of domains that recognise a specific pCTD modification. The canonical two-anchor mode of recognition, wherein the BRCT repeat specifically recognises the phosphoserine at position 5 and the tyrosine at position 1 of the following repeat in the CTD, aligns well with existing crystal structures. ^66–70^ The CTD diheptad is a relatively poor binding substrate (by at least an order of magnitude) compared with other known BRCA1 BRCT ligands. ^67–70,96^ However, given the repetitive nature of the CTD, this lower affinity is likely compensated by avidity effects stemming from the 52-repeat CTD on RNAPII, as evidenced by the biolayer interferometry (BLI) measurements with a 26-repeat CTD. A similar effect has been proposed for other domains recognising the CTD. ^97^

When exploring the functional consequences of the interaction between BRCA1-BARD1 and RNAPII, we found that the complex forms condensates *in vitro* which can simultaneously accommodate pCTD (a likely proxy for RNAPII *in vivo*) and an RNA transcript. Intriguingly, the isolated BRCA1 and BARD1 BRCT domains also formed condensates that included pCTD and RNA. As the BRCA1-BARD1 complex has been implicated in specific condensates *in vivo,* ^74^ our data suggest that the complex may not only be recruited to such condensates but could also actively participate in their formation. Moreover, we and others have recently shown that other DNA repair factors, such as the trimeric SOSS1 complex ^57^ and 53BP1 ^24^ promote condensation *in vivo* at DSB sites, which also contain RNAPII. Thus, it is possible to speculate that the BRCA1-BARD1 complex might promote the organisation of condensates containing RNAPII *in vivo*. Whether these condensates occur precisely at DSB sites – where BRCA1-BARD1 is also present – or at other loci, remains to be determined.

To strengthen our *in vitro* observations, we characterised several disease-associated variants of the BRCT repeats. We identified variants that showed defects in their ability to form condensates, possibly linking defective condensation to pathology.

In conclusion, our study provides mechanistic insight into the interaction between the BRCA1-BARD1 complex and RNAPII and proposes potential functional roles. We anticipate that our results will enable better interpretation of existing *in vivo* data and may guide future studies aimed at uncovering the biological relevance of the BRCA1-BARD1-RNAPII association.

## Materials and Methods

### Plasmid construction

The plasmids were constructed using ligation-independent cloning (LIC), a method, which enables the insertion of DNA fragments into plasmids without the requirement for DNA ligase, and enables the combination of multiple inserts into a single plasmid. ^98^

To express BRCA1 BRCT domains (amino acids 1646-1859) in *E. coli*, fragment of DNA, containing the coding sequence was amplified by PCR and cloned into 2BT (pET His6 TEV LIC cloning vector; addgene #29666) using ligation-independent cloning. Point mutations resulting in the following amino acid substitutions: S1655F, E1682K, R1699W, R1699L, K1702M, E1754K, S1755Y, M1775K, were introduced by site-directed mutagenesis.

To express BARD1 BRCT domains (amino acids 554-777) in *E. coli*, fragment of DNA, containing the coding sequence was amplified by PCR and cloned into 2BT vector. Point mutations resulting in the following amino acid substitutions: S575F, E587K, K619A, E665K, R705A, S711R, K754N, were introduced by site-directed mutagenesis.

To generate plasmids for expression of Avi-tagged BRCA1 BRCT domains (amino acids 1646-1859) in *E. coli*, fragment of DNA, containing the coding sequences of BRCA1 BRCT w.t., S1655F, K1702A, and E1754K, respectively, were amplified by PCR and cloned into H6- msfGFP plasmid (pET Biotin His6 GFP LIC cloning vector; addgene #29725) to add Avi-tag to the sequence. To generate plasmids for expression of Avi-tagged BARD1 BRCT domains (amino acids 554-777), the fragments of DNA, containing the coding sequences of BARD1 BRCT w.t., and S575F+K619A, E665K, R705A, or K754N were amplified by PCR and cloned into H6-msfGFP plasmid (pET Biotin His6 GFP LIC cloning vector; addgene #29725) to add Avi-tag to the sequence. Subsequently, the Avi-tagged fragments were subcloned into 2AT plasmid (pET LIC cloning vector; addgene #29665).

To generate plasmid enabling the expression of biotin-ligase BirA, the coding sequence of BirA was amplified by PCR and cloned into plasmid 2CT-10 (pET His10 MBP Asn10 TEV LIC cloning vector; addgene #55209).

To express the DNA binding region of BRCA1 (amino acids 421-1079), the fragment of DNA, containing the coding sequence was amplified by PCR and cloned into 438C (pFastBac His6 MBP Asn10 TEV cloning vector with BioBrick PolyPromoter LIC Subcloning, addgene #55220).

To generate a vector for co-expression of the full-length human BRCA1-BARD1 complex in insect cells, fragment of DNA, containing FLAG-tagged BRCA1 was first cloned into 2BcT plasmid (pET His6 LIC cloning vector; addgene #37236) to add C-terminal (His)_6_ tag to the construct. Subsequently, FLAG-BRCA1-His was subcloned into plasmid 438A (pFastBac cloning vector with BioBrick PolyPromoter LIC Subcloning, addgene #55218). The DNA fragment containing the ORF for BARD1 was cloned into 438B plasmid (pFastBac His6 TEV cloning vector with BioBrick PolyPromoter LIC Subcloning; addgene #55219). Subsequently, 438A-FLAG-BRCA1-(His)_6_ and 438B-BARD1 plasmids were combined using BioBrick Polypromoter LIC subcloning into a single construct.

The plasmids enabling expression of the kinase module of TFIIH complex (the CDK7 complex) in insect cells, and the plasmid enabling the expression of the catalytic domain of cABL were described in^57^. Plasmids 2BcT-msfGFP-hCTD and 2BcT-mCherry-hCTD were described in^77^. The plasmid enabling the expression of the catalytic domain of DYRK1A was a gift from Nicola Burgess-Brown (Addgene plasmid #38913). Plasmid pGEX4T1-(CTD)_26_- (His)_7_ (provided by Olga Jasnovidova) was used to express and purify GST-(CTD)_26_- (His)_7_. Plasmid pDRIVE-ITS1 used to generate ITS1 RNA by *in vitro* transcription was kindly provided by Stepanka Vanacova.

Plasmids 2AT, 2BT, 2CT-10, 2GT, 2Bc-T, H6-msfGFP, 438A, and 438B were purchased directly from QB3 Macrolab (UC Berkeley). Oligonucleotides used in this study are listed in Supplementary Table 2, list of recombinant plasmids generated in this work is provided in Supplementary Table 3. The sequence integrity of all constructs was verified by sequencing.

### Synthetic RNA/DNA substrates

Oligonucleotides for preparing synthetic fluorescently labelled (Cy3) RNA/DNA substrates were purchased from Sigma (HPLC purified). Substrates were prepared by mixing 3 pmol of labelled oligonucleotides with a 3-fold excess of the unlabelled oligonucleotides in the annealing buffer (25 mM Tris-HCl, pH 7.5; 100 mM NaCl; 3 mM MgCl_2_), followed by initial denaturation at 75°C for 5 min. Substrates were then purified from a native PAGE gel.

### Protein expression in *E. coli*

His-BARD1^554–777^, Avi-BARD1^554-777^ and MBP-BirA were expressed in *E. coli* RIPL strain. BRCA1^1646-1859^ and Avi-BRCA1^1646-1859^ were expressed in *E. coli* Rosetta 1 strain. The expression of the proteins was induced at OD_600_ = 0.6 by addition of IPTG (final concentration 1 mM) and the cultures were incubated overnight at 16°C.

GST-(CTD)_26_, mCherry-hCTD, and GFP-hCTD were expressed in *E. coli* BL21-AI strain grown in TB media. The expression of the proteins was induced at OD_600_ = 0.6 by addition of IPTG (final concentration 0.1 mM) and L-arabinose (final concentration 0.02%) and the cultures were incubated overnight at 30°C.

### Protein expression in insect cells

To generate viruses enabling the production of proteins in insect cells, the coding sequences and the necessary regulatory sequences of the aforementioned constructs were transposed into bacmids using the *E. coli* strain DH10bac. Viral particles were obtained by transfection of the corresponding bacmids into the *Sf*9 cells using the FuGENE Transfection Reagent and further amplification. Proteins were expressed in 300 – 600 ml of Hi5 cells (infected at 1.2×10^6^ cells/ml) with the corresponding P1 virus at multiplicity of infection (MOI) >1. Cells were harvested 48 hours post-infection, washed with 1xPBS, and stored at-80°C.

### Protein purification

#### Purification of BRCA1^1646-1859^ and BARD1^554-777^

Eight grams of *E. coli* pellet was resuspended in 40 ml of ice-cold lysis buffer (50 mM Tris-HCl, pH 7.9; 500 mM NaCl; 1 mM DTT; 10 mM imidazole) containing one tablet of cOmplete^TM^ EDTA-free Protease Inhibitors. Cells were opened up by sonication, the lysate was cleared by centrifugation and incubated for 1 hr with 5 mL of Ni-NTA beads (Qiagen), equilibrated with the lysis buffer. The proteins were eluted with an elution buffer containing 25 mM Tris-HCl, pH 7.9; 300 mM NaCl; 1 mM DTT; and 400 mM imidazole. Elution fractions were dialysed overnight against dialysis buffer (25 mM Tris-HCl, pH 7.9; 300 mM NaCl; 1 mM DTT). The (His)_6_-tag was cleaved off by the TEV protease during the dialysis step. The sample was then purified from the TEV protease using Ni-NTA beads, concentrated, and loaded onto Superdex S-75 column equilibrated in a buffer containing 25 mM Tris-HCl, pH 7.9; 300 mM NaCl; and 1 mM DTT. The samples for phase separation assays were purified using Superdex S-75 column equilibrated in a buffer containing 25 mM HEPES, pH7.5; 220 mM NaCl; 1 mM DTT. The fractions containing purified protein were concentrated, snap-frozen in liquid nitrogen, and stored at-80°C.

#### Purification of BirA, Avi-BRCA1^1646-1859^ and Avi-BARD1^554-777^

Eight grams of *E. coli* pellet was resuspended in 40 ml of ice-cold lysis buffer (50 mM Tris-HCl, pH 7.9; 500 mM NaCl; 1 mM DTT; 10 mM imidazole) containing one tablet of cOmplete, EDTA-free Protease Inhibitors. Cells were opened up by sonication, the lysate was cleared by centrifugation and incubated for 1 hr with 5 mL of Ni-NTA beads (Qiagen), equilibrated with the lysis buffer. The proteins were eluted with an elution buffer containing 25 mM Tris-HCl, pH 7.9; 300 mM NaCl; 1 mM DTT and 400 mM imidazole. Elution fractions were concentrated, and the proteins were further purified using gel filtration column equilibrated in buffer containing 50 mM Tris-HCl, pH 7.9; 300 mM NaCl and 1 mM DTT. BirA was purified using Superdex S-200 column, Avi-BRCA1^1646-1859^ and Avi-BARD1^554-777^ were purified using Superdex S-75 column. The fractions containing purified protein were concentrated, snap-frozen in liquid nitrogen, and stored at-80°C.

#### In vitro biotinylation and purification of biotinylated BRCA1^1646-1859^ and BARD1^554-777^

For preparative purposes, 1 mg of BRCA1^1646-1859^ and Avi-BARD1^554-777^, respectively, were biotinylated by 800 µg of the MBP-BirA in the presence of 3 mM ATP, 3 mM MgCl_2_, and 200 µM biotin for 2 hrs at 23°C. Reactions were loaded directly onto Superdex S-200 equilibrated with 25 mM Tris-HCl, pH 7.9, 250 mM NaCl, 1 mM DTT in order to purify the biotinylated proteins from BirA and ATP. Fractions containing biotinylated BRCA1^1646-1859^ and BARD1^554-777^ were pooled, concentrated, snap-frozen in liquid nitrogen, and stored at-80°C.

#### Purification of BRCA1^421-1079^ from insect cells

Pellets from 300 ml of the Hi5 culture were resuspended in ice-cold lysis buffer (50 mM Tris-HCl, pH 7.9; 500 mM NaCl; 10 % (v/v) glycerol; 1 mM DTT; 0.4 % (v/v) Triton-X; 10 mM imidazole) containing protease inhibitors (0.66 μg/ml pepstatin, 5 μg/ml benzamidine, 4.75 μg/ml leupeptin, 2 μg/ml aprotinin) and 25 U benzonase per ml of lysate. The cleared lysate was incubated for 1 hr with 3 mL of Ni-NTA beads (Qiagen), equilibrated with 50 mM Tris-HCl, pH 7.9; 500 mM NaCl; 10 mM imidazole; and 1 mM DTT. The proteins were eluted with elution buffer containing 25 mM Tris-HCl, pH 7.9; 300 mM NaCl; 1 mM DTT and 400 mM imidazole. Elution fractions were concentrated, and the proteins were further purified using Superdex S-200 column equilibrated in buffer containing 50 mM Tris-HCl, pH 7.9; 300 mM NaCl and 1 mM DTT. Fractions containing purified protein were concentrated, and glycerol was added to a final concentration of 10 % before they were snap-frozen in liquid nitrogen, and stored at-80°C.

#### Purification of the BRCA1-BARD1 complex from insect cells

Pellets from 600 ml of the Hi5 culture were resuspended in ice-cold lysis buffer (50 mM Tris-HCl, pH 7.5; 500 mM KCl; 10 % (v/v) glycerol; 5mM MgCl_2_; 2mM ATP; 1 mM DTT; 0.5 % (v/v) NP-40; 10 mM imidazole) containing protease inhibitors (0.66 μg/ml pepstatin, 5 μg/ml benzamidine, 4.75 μg/ml leupeptin, 2 μg/ml aprotinin) and 25 U benzonase per ml of lysate. The cleared lysate was incubated for 1 hr with 3 mL of Ni-NTA beads (Qiagen), equilibrated with the lysis buffer. The beads were then washed extensively with the wash buffer (25 mM Tris-HCl, pH 7.5; 500 mM KCl; 10 % (v/v) glycerol; 5mM MgCl_2_; 2mM ATP; 1 mM DTT; 10 mM imidazole, protease inhibitors) and subsequently, the proteins were eluted with the elution buffer (25 mM Tris-HCl, pH 7.5; 300 mM KCl; 10 % (v/v) glycerol; 5mM MgCl_2_; 2mM ATP; 1 mM DTT; 400 mM imidazole, protease inhibitors). Elution fractions were concentrated, His-tags were cleaved off using TEV protease and the proteins were further purified using Superose 6 column equilibrated in the gel filtration buffer containing 50 mM Tris-HCl, pH 7.5; 300 mM KCl; and 1 mM DTT. For the biochemical purposes, fractions containing purified protein were concentrated, supplemented with glycerol to a final concentration of 10 %, snap-frozen in liquid nitrogen, and stored at-80°C.

#### Purification of ABL1^cat^ and DYRK1A^cat^

Five grams of *E. coli* BL21 RIPL cells expressing the catalytic domain of ABL1 and DYRK1A, respectively, were resuspended in ice-cold lysis buffer (50 mM Tris-HCl, pH 8; 0.5 M NaCl; 10 mM imidazole; 1 mM DTT), containing protease inhibitors (0.66 μg/ml pepstatin, 5 μg/ml benzamidine, 4.75 μg/ml leupeptin, 2 μg/ml aprotinin) at +4°C. Cells were opened up by sonication. The cleared lysate was passed through 2 mL of Ni-NTA beads (Qiagen), equilibrated with buffer (50 mM Tris-HCl, pH 8; 500 mM NaCl; 10 mM imidazole; and 1 mM DTT). Proteins were eluted with an elution buffer (50 mM Tris-HCl, pH 8; 500 mM NaCl; 1 mM DTT and 400 mM imidazole). The elution fractions containing the proteins of interest were pooled, concentrated, and further fractionated on Superdex S-75 column with SEC buffer (25 mM Tris-Cl pH7.5; 200 mM NaCl, 1 mM DTT). Fractions containing pure proteins of interest were concentrated, glycerol was added to a final concentration of 10 % before they were snap-frozen in liquid nitrogen, and stored at-80°C.

Purified T7 RNAP was kindly provided by Martin Mátl (Ceitec, MU). The purification of the CTD peptides and their subsequent *in vitro* phosphorylation was described earlier. ^57,77^

### Pull-down assays from HEK293T cell lysates

HEK293 human cell lines were cultivated in Dulbecco’s modified Eagle’s medium (DMEM, Sigma) supplemented with 10% (v/v) fetal bovine serum (FBS, Tet-free approved, Sigma), 100 U/ml penicillin and 100 μg/ml streptomycin (Gibco). Cells were grown at 37°C in humidified atmosphere in 5% CO2. Typically, one HEK293 pellet from one 10 cm dish was used per condition. The pellet was resuspended in 1 ml of NETN buffer (50 mM Tris-HCl, pH 8; 150 mM NaCl; 1 mM EDTA; 0.5% NP-40; BSA (final concentration 0.1 mg/ml)) containing benzonase nuclease and cOmpleteTM, EDTA-free Protease Inhibitors. The sample was incubated for 30 min at 4°C and sonicated briefly. The lysate was cleared by centrifugation and the supernatant was incubated with 5 μg of FLAG-tagged BRCA1-BARD1 for 30 min at 4°C. Subsequently, the sample was added to 25 μl of ANTI-FLAG ® M2 Affinity Gel (Sigma), equilibrated in NETN buffer, and incubated for 30 min at 4°C. The beads were washed four times with 4 CV of NETN buffer and the bound proteins were eluted with 50 μl of the NETN buffer containing 3xFLAG peptide (Sigma) (final concentration 0.25 mg/ml). The input and elute were analyzed by western blotting.

### *In vitro* pull-down assay

Purified GST-CTD, GST-Y1P-CTD, GST-S2,5P-CTD, and GST-S5,7P-CTD (5 μg each), respectively, were incubated with BRCA1-BARD1, BRCA1 BRCT, and BARD1 BRCT (5 μg each) in 30 μl of buffer T200 (25 mM Tris–HCl pH7.5; 200 mM NaCl; 10 % glycerol; 1 mM DTT; 0.5 mM EDTA; and 0.01% NP-40) for 30 min at 4°C in the presence of GSH-beads. After washing the beads twice with 100 μl of buffer T200, the bound proteins were eluted with 30 μl of 4xSDS loading dye. The input, supernatant, and eluate, 7 μl each, were analysed on SDS-PAGE gel.

### Electrophoretic Mobility-Shift Assay (EMSA)

Increasing concentrations of tested proteins (25, 50, 100, and 200 nM of BRCA1^421-1079^ and 3.125, 6.25, 12.5, and 25 µM of BRCA1^1646-1859^ and BARD1^554-777^, respectively) were incubated with fluorescently labelled nucleic acid substrates (at a final concentration of 10 nM) in EMSA buffer (25 mM Tris-HCl, pH 7.5; 1 mM DTT; 5 mM MgCl_2_; and 100 mM NaCl) for 20 min on ice. Loading buffer (60 % glycerol in 0.001% Orange-G) was added to the reaction mixtures and the samples were loaded onto a 7.5 % (w/v) polyacrylamide native gel in 0.5 x TBE buffer and run at 75 V for 1h at +4°C. The different nucleic acid species were visualised using FLA-9000 Starion scanner and quantified in the MultiGauge software (Fujifilm). To calculate the relative amount of bound nucleic acid substrate, the background signal from the control sample (without protein) was subtracted using the band intensity - background option. Nucleic acid-binding affinity graphs were generated with Prism-GraphPad 9.

#### Competition EMSA assays in the presence of CTD polypeptide

To assess the effect of phosphorylated CTD on the biding of BRCA1^1646-1859^ or BARD1^554-777^ to nucleic acid substrate, BRCA1^1646-1859^ and BARD1^554-777^ (12.5 µM), respectively, were incubated with the nucleic acid substrate (10nM) for 15 min on ice. Subsequently, increasing concentrations (0.1, 0.3, 0.9 mM) of the pS5pS7 GST-CTD_26_ were added, and the reaction mixtures were further incubated for 15 min on ice. Reactions were next processed as described above. The statistical significance was determined by unpaired t-test analysis.

### *In vitro* crosslinking experiments

To determine the oligomeric state of BRCA1^1646-1859^ and BARD1^554-777^, respectively, the proteins were diluted to 0.5 mg/ml and 2.5 mg/ml, respectively, in buffer H_150_ (25 mM HEPES pH7.5; 150 mM NaCl; 1 mM DTT). To the diluted proteins, glutaraldehyde (0.05% final concentration) was added, and the mixture was incubated for 30 min at 4°C. Subsequently, the reactions were stopped by the addition of 1µl of 1M Tris-HCl. The reactions were analysed by SDS-PAGE.

### Differential scanning fluorimetry

The thermal stability of BRCA1^1646-1859^ and BARD1^554-777^ w.t. and their mutant variants was measured using the Prometheus NT.38 instrument (NanoTemper Technologies). NanoDSF grade capillaries were filled with protein samples at 1 mg/ml in protein dilution buffer (25 mM Tris–HCl, pH 7.9; 250 mM NaCl; and 1 mM DTT). Protein unfolding was detected at a temperature range of 20 to 95°C with a 1°C/min heating rate and 90% excitation power. Onset of protein unfolding (To) and protein melting temperature (Tm) were determined from the ratio of tryptophan emission at 330 and 350 nm.

### Fluorescence anisotropy binding assays

The binding of BRCA1^1646-1859^ and BARD1^554-777^ to CTD peptides was characterised using FluoroLog-3 spectrofluorometer (Horiba Jobin-Yvon Edison) equipped with a thermostatic cell holder with a Neslab RTE7 water bath (Thermo Scientific). 5,6-FAM-labelled CTD peptides, purchased from Caslo ApS, were diluted to 25 nM in FA buffer (25 mM Tris–HCl, pH 7.9; 150 mM NaCl; and 1 mM DTT). BRCA1^1646-1859^ and BARD1^554-777^ samples were titrated against a constant concentration of fluorescently labelled peptides at 25°C. Samples were excited with vertically polarized light at 467 nm, and both vertical and horizontal emissions were recorded at 516 nm. The experiments were performed in technical triplicates. Anisotropy data were plotted as a function of protein concentration and fitted to a single-site saturation with non-specific binding model using XMGrace. The graphs were generated using GnuPlot.

### Biolayer-interferometry (BLI)

To characterise the kinetics of binding of phosphorylated CTD to BRCA1^1646-1859^ and BARD1^554-777^, BLI experiments were performed using Octet RED96e (ForteBio) at 25°C, with shaking at 2200 rpm. The streptavidin biosensors (SA, ForteBio) were pre–hydrated in BLI buffer (25 mM Tris–HCl, pH 7.9; 150 mM NaCl; 1 mM DTT; and 0.05% (v/v) Tween 20) for 10 min. Biotinylated BRCA1^1646-1859^ and BARD1^554-777^, respectively, were immobilised on the biosensor tip at a final concentration of 10 µg/ml. The association of non-phosphorylated or pS5pS7 phosphorylated GST-(CTD)_26_ to the immobilised proteins was measured at five different concentrations of the analyte (1.4, 4.3, 13, 39, and 117 nM) for 300 s. The dissociation of the GST-CTD_26_ was measured for 300 s by subsequent washing of the biosensor with BLI buffer. The experiments were performed in triplicates. The association and dissociation constants and the coefficient of determination (R^2^) indicating the appropriateness of the fit were calculated in Octet® Analysis Studio Software using 1:2 Bivalent analyte model. The data and the fits were plotted using Prism GraphPad 9 software.

### *In vitro* transcription assay

The pDRIVE-ITS1 plasmid was purified from 200 ml of *E. coli* culture using Plasmid Maxi Kit (Qiagen) and linearised by BamHI enzyme. The linearised plasmid was purified from the agarose gel using QIAquick Gel Extraction Kit (Quiagen) and used as a template for the *in vitro* transcription. For the preparation of the Cy5-labelled ITS1 RNA, 8 µg of the template DNA was mixed with ATP, UTP, and GTP (at 2mM), CTP (at 1mM), Cy5-labelled CTP (5- Propargylamino-CTP-Cy5, Jena Bioscience, at 0.05 mM), and T7 RNA polymerase (at 1 µM) in a buffer containing 0.1 M Tris-HCl, pH 8.1; 1% Triton X-100; 16 mM MgCl_2_; 10 mM spermidine; and 50 mM DTT. The reaction was incubated for 4 hrs at 37°C. The product was purified using RNeasy Mini Kit (Qiagen) and stored at-80°C.

### *In vitro* fluorescent labelling of proteins

BRCA1-BARD1, BRCA1^1646-1859^, and BARD1^554-777^, were fluorescently labelled using Alexa Fluor® 488 Conjugation Kit - Lightning-Link® (Abcam) according to the manufacturer’s instructions. Labelled proteins were snap-frozen in liquid nitrogen and stored at-80°C. For the *in vitro* liquid-liquid phase separation assays, the labelled proteins were mixed with the same protein lacking the fluorescent tag in a 1:20 molar ratio.

### *In vitro* liquid-liquid phase separation (LLPS) assays

#### LLPS assays with the full length BRCA1-BARD1 complex

The LLPS assays with non-labelled and Alexa Fluor 488-conjugated BRCA1-BARD1 (final concentrations 1.25, 2.5, and 5 µM), respectively, were performed in the reaction buffer (25 mM HEPES, pH 7.5; 300 mM NaCl; 1 mM DTT) in the presence of a crowding agent (10% (w/v) dextran). Where indicated, Cy5-labelled ITS1 RNA (at 15 nM) and/or pS5pS7 phosphorylated hCTD (at 2.5 µM) were added.

#### LLPS assays with BRCTs and pCTD

The LLPS assays with Alexa Fluor 488-conjugated BRCA1^1646-1859^ and BARD1^554-777^ (w.t. and mutant variants at 160 µM), respectively, were performed in the reaction buffer (25 mM HEPES, pH 7.5; 220 mM NaCl; 1 mM DTT) in the presence of a crowding agent (10% (w/v) dextran). Where indicated, pS5pS7 phosphorylated mCherry-hCTD (at 2.5 µM) was added. The LLPS assays with non-labelled or Alexa Fluor 488-conjugated BRCA1^1646-1859^ and BARD1^554-777^ (w.t. and mutant variants, at 40, 80, and 160 µM), respectively, and pS5pS7 phosphorylated GFP-hCTD (at 2.5 µM) were performed in the reaction buffer (25 mM HEPES, pH 7.5; 220 mM NaCl; 1 mM DTT) in the presence or absence of a crowding agent (10% (w/v) dextran).

#### LLPS assays with BRCTs and ITS1 RNA

The LLPS assays with Alexa Fluor 488-conjugated BRCA1^1646-1859^ and BARD1^554-777^ (w.t. or mutant variants, final concentrations 10, 20, and 40 µM), respectively, and Cy5-labelled ITS1 RNA (final concentration 15 nM) were performed in the reaction buffer (25 mM HEPES, pH 7.5; 220 mM NaCl; 1 mM DTT) in the presence of a crowding agent (10% (w/v) PEG 8000). The LLPS assays with non-labelled BRCA1^1646-1859^ and BARD1^554-777^ (w.t. and mutant variants, at 160 µM), respectively, and Cy5-labelled ITS1 RNA (final concentration 15 nM) were performed in the reaction buffer (25 mM HEPES, pH 7.5; 220 mM NaCl; 1 mM DTT) in the presence of a crowding agent (10% (w/v) dextran).

#### LLPS assays with BRCTs, ITS1 RNA, and pCTD

The LLPS assays with non-labelled and Alexa Fluor 488-labelled BRCA1^1646-1859^ and BARD1^554-777^ (concentration 10 µM), respectively, were performed in the reaction buffer (25 mM HEPES, pH 7.5, 220 mM NaCl, 1 mM DTT) in the presence of a crowding agent (10% (w/v) PEG 8000). Where indicated, Cy5-labelled ITS1 RNA (final concentration 15 nM) and/or pS5pS7 phosphorylated hCTD (final concentration 2.5 µM), and 3mM MgCl_2_ were added.

#### LLPS assays with BRCT cancer-associated mutants

Alexa Fluor 488-conjugated BRCA1^1646-1859^ and BARD1^554-777^ (w.t. or mutant variants, final concentrations 40, 80, and 160 µM) were mixed with a crowding agent (10% (w/v) dextran) in the reaction buffer (25 mM HEPES, pH 7.5, 220 mM NaCl, 1 mM DTT) and incubated for 10 min on ice.

#### Data collection

The mixtures were immediately spotted onto a glass slide, and the condensates were recorded on Zeiss Axio Observer.Z1 with a 63x water immersion objective.

#### Statistical analyses

Analyses and quantifications of the micrographs were performed in Cell-Profiler (version 4.2.6.).^99^ First, five micrographs (2048 pixels (px) per 2048 px, 1 px= 0.103 µm) per condition and per experiment were analysed. Objects (droplets) were identified based on diameter (2-70 px, 0,206-7.5 µm) and intensity using Otsu’s method for thresholding. Picked objects were further filtered based on shape and intensity. For the filtered objects the area, median intensity of the objects for the GFP channel, and the object count per pictures were calculated. The values for droplets areas were converted from the px to µm based on the metadata of the micrographs (0.010609 µm2 = 1px). Graphs for the figures were plotted using Prism GraphPad 9 software. ^100^ Significance is listed as *p ≤ 0.05, **p ≤ 0.01, ***p ≤ 0.001, ****p ≤ 0.0001.

### Crystallisation and data collection

Purified BRCA1 BRCT was concentrated to 9.75 mg/ml and mixed with pS5-CTD peptide in 2:1 ratio (final concentrations 6.5 mg/ml and 3.25 mg/ml of the BRCT and the CTD peptide, respectively) and crystallized using the sitting-drop vapour diffusion method at 4°C. The crystals were obtained using the precipitant containing 0.1M PCB buffer (sodium propionate, sodium cacodylate, and BIS-TRIS propane in the molar ratios 2:1:2), pH = 6, 25% PEG 1500, from the PACT Suite crystallization screen (Qiagen). The crystals were cryoprotected with 40% PEG 400 and frozen in liquid nitrogen. The diffraction data were collected at the PETRA III electron storage ring beamline P13 (DESY, Hamburg, Germany).

## Structure determination

Diffraction images were processed using the programs XDS ^101^ and converted to structure factors using the program Scala from the package CCP4 v.8.0, ^102^ with 5% of the data reserved for the R_free_ calculation. The structures of complexes were solved by molecular replacement with Phaser. ^103^ The BRCA1 BRCT structure (PDB ID: 1JNX) was used as the initial coordinates. The refinement was performed using REFMAC5 ^104^ alternated with manual model building in Coot v.0.9. ^105^ The bound peptides were built up manually using Coot. Molecular drawings were prepared using Pymol (Schrödinger, Inc.). The structures were visualised using UCSF ChimeraX package (Resource for Biocomputing, Visualization, and Informatics at the University of California, San Francisco). ^106^

### Small angle X-ray scattering (SAXS)

SAXS data were collected using a Rigaku BioSAXS-2000 instrument equipped with a HyPix-3000 detector at a sample-detector distance of 0.5 m (I(q) vs s, where q= 4π sin θ/λ; 2θ is the scattering angle and λ = 0.154 nm). One 10 minutes frame were collected at a sample temperature of 20°C. The data were normalized to the intensity of the transmitted beam and radially averaged and the corresponding scattering from the solvent-blank was subtracted to produce the scattering profile.

Radius of gyration and other integral structural parameters were determined using PRIMUS.

### AlphaFold 3 modelling of the BRCT structures

The models of BARD1^554–777^ with pS5 CTD peptide, BRCA1^1646-1859^ with RNA oligo (ACGGAGCCCG) and BARD1^554-777^ with RNA oligo (ACGGAGCCCG) were built based on the prediction of AlphaFold 3. The top ranked prediction was used for the visualization.

## Data availability

All primary data are available in this manuscript, supplementary information, and source data.

## Supplementary data

Supplementary data are available online.

## Funding

This work was supported by the Junior Star Grant from the Grant Agency of the Czech Republic (21-10464M) awarded to MS.

We acknowledge the core facility CELLIM supported by the Czech-BioImaging large RI project (LM2023050 funded by MEYS CR) for their support with obtaining scientific data presented in this paper.

CIISB, Instruct-CZ Centre of Instruct-ERIC EU consortium, funded by MEYS CR infrastructure project LM2023042 and European Regional Development Fund-Project “Innovation of Czech Infrastructure for Integrative Structural Biology” (No. CZ.02.01.01/00/23_015/0008175), is gratefully acknowledged for the financial support of the measurements at the CF Biomolecular Interactions and Crystallography.

## Conflict of interest

The authors declare that they have no conflict of interest.

## Acknowledgement

We thank members of the lab for their support. We thank Katerina Linhartova (CEITEC, Masaryk University) for sharing the 2BcT-GFP-hCTD and 2BcT-mCherry-hCTD plasmids and for assistance with the analysis of the LLPS experiments; Stepanka Vanacova (CEITEC, Masaryk University) for sharing the pDRIVE-ITS1 plasmid; and Olga Jasnovidova (Technical University of Tallinn) for sharing the pGEX4T1-(CTD)_26_-(His)_7_ plasmid. We thank Martin Matl (CEITEC, Masaryk Universty) for sharing T7 RNAP. We thank Jana Kosourova (BIC, CEITEC, Masaryk University) for assistance with crystallisation. We thank Eva Paulenova (BIC, CEITEC, Masaryk University) for assistance with nanoDSF. We thank Jakub Pospisil and Milan Esner (CELLIM, CEITEC, Masaryk University) for assistance with fluorescence microscopy.

## Author contributions

Experimentation: VK, KS; Protein purification: VK; *In silico* modelling: VK, JH; Structure determination: JH Conceptualisation: MS; Funding acquisition: MS; Project administration: MS; Supervision: MS; Writing – original draft: VK, MS; Writing – review and editing: VK, JH, MS

## Notes

### Competing Interest Statement

The authors have declared no competing interest.

### Summary of Updates

In the revised version, defects in export of Supplementary Fig. 2B; 7B; 8A,C,D;12D,E were corrected. Otherwise the no further changes were made to the main text nor figures.

