## Supplementary material for "Distinct Mechanisms of Recognition of Phosphorylated RNAPII C-Terminal Domain by BRCT Repeats of the BRCA1–BARD1 Complex: Insights from Structural and Functional Analyses": Supplentary material

Supplementary Figures 1-23

Supplementary Tables 1-4

#### Supplementary Figure Legends:

##### Supplementary Figure 1: Purified proteins used in this study

**A)** SDS-PAGE gels showing purified BRCA1 BRCT variants used in this study.

**B)** SDS-PAGE gels showing purified BARD1 BRCT variants used in this study.

**C)** SDS-PAGE gels showing purified CTD variants used in this study**.**

**D)** SDS-PAGE gel showing purified BRCA1-BARD1 used in this study.

##### Supplementary Figure 2:

**A)** SDS-PAGE gel analysis of pull-downs from HEK293 lysates.

**B)** Western blot analyses of pull-downs from HEK293 lysates. HEK293 cells were lysed and the lysate was cleared by centrifugation. To the supernatant, FLAG-BRCA1-BARD1 was added, and the samples were incubated with α-FLAG beads. As a control, the HEK293 lysate without added BRCA1-BARD1 was used. The proteins were eluted using 3xFLAG peptide and the samples were analysed using western blots. BRCA1-BARD1 interacts with RNAPII via the CTD phosphorylated on Ser2 and Ser5.

##### Supplementary Figure 3:

**A)** SDS-PAGE analysis of *in vitro* pull-down assay between GST-$(CTD)_{26}$ and BRCA1-BARD1. Purified BRCA1-BARD1 was incubated with phosphorylated and non-phosphorylated GST-$(CTD)_{26}$ bound to glutathione beads. The samples were centrifuged and the input, unbound (supernatant) and bound (pellet) fractions were analysed using the SDS PAGE. BRCA1-BARD1 interacts directly with pS2pS5 GST-$(CTD)_{26}$ and pS5pS7 GST-$(CTD)_{26}$ *in vitro*.

**B)** SDS-PAGE analysis of *in vitro* pull-down assay between GST-$(CTD)_{26}$ and BRCA1 BRCT and BARD1 BRCT, respectively. Purified BRCA1 BRCT and BARD1 BRCT, respectively, were incubated with phosphorylated and non-phosphorylated GST-$(CTD)_{26}$ bound to glutathione beads. The samples were centrifuged and the input, unbound (supernatant) and bound (pellet) fractions were analysed using the SDS PAGE. Substitutions in the phosphoserine binding site (BRCA1$^{S1655F, K1702M}$, BARD1$^{S575F, K619A}$) abolish the binding.

##### Supplementary Figure 4:

**A)** Multiple sequence alignment of amino-acids present in phosphopeptide-binding sites of BRCA1 BRCT domains from various organisms. The alignment was generated using Clustal Omega on EMBL-EBI and visualised using Jalview.

**B)** Multiple sequence alignment of amino-acids present in phosphopeptide-binding sites of BARD1 BRCT domains from various organisms. The alignment was generated using Clustal Omega on EMBL-EBI and visualised using Jalview.

##### Supplementary Figure 5:

**A)** Multiple sequence alignment of sequences of BRCA1 BRCT domains from various organisms. The alignment was generated using Clustal Omega on EMBL-EBI and visualised using Jalview.

**B)** Multiple sequence alignment of sequences of BARD1 BRCT domains from various organisms. The alignment was generated using Clustal Omega on EMBL-EBI and visualised using Jalview.

**Supplementary Figure 6:**

**A)** Sensograms obtained by biolayer-interferometry (BLI). Analysing interactions of BRCA1 $BRCT^{S1655F,K1702M}$, BRCA1 $BRCT^{R1699L}$, and BARD1 $BRCT^{S575F,K619A}$, respectively, with pS5pS7 GST-$(CTD)_{26}$ and GST-$(CTD)_{26}$.The sensograms represent the mean of 3 measurements for each concentration. The data were analysed in Octet® Analysis Studio Software using 1:2 Bivalent analyte model. The data were plotted using Prism GraphPad 9 software.

**B)** Structural alignment of BRCA1 BRCT (1JNX, teal) and BARD1 BRCT (2NTE, purple) obtained in UCSF Chimera.

**C)** Sensograms obtained by biolayer-interferometry (BLI) analysing interactions of BRCA1 $BRCT^{R1699S}$, BRCA1 $BRCT^{M1775H}$, and BARD1 $BRCT^{H686M}$, respectively, with pS5pS7 GST-$(CTD)_{26}$. The sensograms represent the mean of 3 measurements for each concentration. The data were analysed in Octet® Analysis Studio Software using 1:2 Bivalent analyte model. The data were plotted using Prism GraphPad 9 software.

**Supplementary Figure 7:**

**A)** Specificity of the BRCA1 BRCT phospho-peptide binding site determined by fluorescence anisotropy measurement. The assays were performed between BRCA1 BRCT domain and the indicated ligands (at 25nM). Anisotropy data were plotted as a function of protein concentration and fitted to a single-site saturation with non-specific binding model using XMGrace.

**B)** Comparison of the apo-structure of BRCA1 BRCT and the structure of BRCA1 BRCT with bound pS5 CTD peptide (YSPTpSPSYSPTpSPS). Teal: newly determined structure of BRCA1 BRCT in complex with pCTD, Grey: BRCA1 BRCT apo-structure (PDB: 1JNX). Structural alignment was generated in UCSF ChimeraX using the Match maker tool.

**C)** Comparison of the phosphoserine binding site of BRCA1 BRCT with (teal) and without (gray) bound pS5 CTD peptide (yellow).

**D)** Distribution of the surface charge of BRCA1 BRCT with docked pS5 CTD peptide. The electrostatic potential was calculated by the Adaptive Poisson-Boltzmann Solver and visualised in Pymol.

**E)** Detail of the binding site of BRCA1 BRCT with the electron density map representing the pS5 CTD peptide present in the chains A, B and C (left, middle and right, respectively). Structural models and electron density were visualised using Coot (version 0.9.8.93).

**Supplementary Figure 8:**

**A)** Comparison of the crystal structure of the BRCA1 BRCT (dark gray) with the S5 pCTD ligand (dark yellow) and the AlphaFold 3 model of the complex (BRCA1 BRCT – light gray, S5 pCTD peptide – light yellow).

**B)** Detail of the phosphoserine binding site (left) of BRCA1 BRCT (gray) with bound pS5 CTD peptide (yellow) modelled by AlphaFold 3. The pS5 of the CTD peptide is depicted in purple, the amino-acid residues interacting with pS5 are depicted in light violet. The hydrogen bonds were displayed as pseudobonds using the structural analysis tool Hydrogen bonds in UCSF ChimeraX and are depicted in turquoise. The relax distance tolerance was 1.0 Å and the relax angle tolerance was 20.0°. Detail of the aromatic amino-acid binding pocket (middle, right) of BRCA1 BRCT (gray) with bound pS5 CTD peptide (yellow). The Y1 of the CTD is depicted in navy, the interacting amino-acid residues in light blue. The hydrogen bonds (middle) were displayed as pseudobonds using the structural analysis tool Hydrogen bonds in UCSF ChimeraX and are depicted in turquoise. The relax distance tolerance was 1.0 Å and the relax angle tolerance was 20.0°. Van der Waals, hydrophobic and stacking interactions (right) were displayed as pseudobonds using the structural analysis tool in UCSF ChimeraX and are depicted in black. The interacting atoms were identified based on VDW overlap ≥ -0.4 Å.

**C)** AlphaFold 3 model of the BARD1 BRCT (gray) with the S5 pCTD (orange).

**D)** Detail of the phosphoserine binding site (left) of BARD1 BRCT (gray) with bound pS5 CTD peptide (yellow) modelled by AlphaFold3. The pS5 of the CTD peptide is depicted in purple, the amino-acid residues interacting with pS5 are depicted in light violet. The hydrogen bonds were displayed as pseudobonds using the structural analysis tool Hydrogen bonds in UCSF ChimeraX and are depicted in turquoise. The relax distance tolerance was 1.0 Å and the relax angle tolerance was 20.0°. Detail of the aromatic amino-acid binding pocket (middle, right) of BARD1 BRCT (gray) with bound pS5 CTD peptide (yellow). The Y1 of the CTD is depicted in navy, the interacting amino-acid residues in light blue. The hydrogen bonds (middle) were displayed as pseudobonds using the structural analysis tool Hydrogen bonds in UCSF ChimeraX and are depicted in turquoise. The relax distance tolerance was 1.0 Å and the relax angle tolerance was 20.0°. Van der Waals, hydrophobic and stacking interactions (right) were displayed as pseudobonds using the structural analysis tool in UCSF ChimeraX and are depicted in black. The interacting atoms were identified based on VDW overlap ≥ -0.4 Å.

**Supplementary Figure 9:**

**A)** LLPS assays with purified BRCA1-BARD1 complex. BRCA1-BARD1 (at 5 μM) was mixed with phosphorylated mGFP-pCTD (2.5 μM) and/or Cy5-ITS1 RNA (at 15 nM) in the presence of a crowding agent (10% dextran). Representative images from three experiments (bottom) are depicted as an overlay of differential interference contrast (DIC), GFP, and Cy5. Scale bars, 10 μm.

**B)** Bar chart (top) representing quantification (n = 3) of the number of droplets per frame from the LLPS experiments with the BRCA1-BARD1 complex shown in (A). Statistical significance was determined by unpaired t-test. A nested scatterplot (bottom) represents quantification (n = 3) of an area of individual droplets from three independent experiments with the BRCA1-

BARD1 complex shown in (A), with median area determined per dataset. Statistical significance was determined by nested t-test.

**C)** Effect of the Alexa-488 labelling on the LLPS propensities of the BRCA1-BARD1 complex. Bar chart (top) representing quantification (n = 3) of the number of droplets per frame from the LLPS experiments with BRCA1-BARD1, pS5pS7 mCherry-hCTD and Cy5-RNA. Statistical significance was determined by unpaired t-test. A nested scatterplot (middle) represents quantification (n = 3) of an area of individual droplets from three independent experiments with BRCA1-BARD1, with median area determined per dataset. Statistical significance was determined by nested t-test. Representative images from three experiments (bottom) are depicted as an overlay of differential interference contrast (DIC) and GFP. Scale bars, 10 μm.

**Supplementary Figure 10:**

**A)** LLPS assays with purified BRCA1 BRCT and pS5pS7 mGFP-hCTD. BRCA1 BRCT (at 40 μM, 80 μM or 160 μM), w. t.; S1655F,K1702M; R1699L; M1775K was mixed with phosphorylated mGFP-pCTD (2.5 μM) in the presence of a crowding agent (10% dextran). Representative images from three experiments are depicted as an overlay of differential interference contrast (DIC) and GFP. Scale bars, 10 μm.

**B)** Bar chart (top) representing quantification (n = 3) of the number of droplets per frame from the LLPS experiments with BRCA1 BRCT w.t. shown in (A). Statistical significance was determined by unpaired t-test. A nested scatterplot (bottom) represents quantification (n = 3) of an area of individual droplets from three independent experiments with BRCA1 BRCT w.t. shown in (A)., with median area determined per dataset. Statistical significance was determined by nested t-test.

**C)** LLPS assays with purified BARD1 BRCT and pS5pS7 mGFP-hCTD. BARD1 BRCT (at 40 μM, 80 μM or 160 μM), w. t., and S575F,K619A, respectively was mixed with mGFP-pCTD (2.5 μM) in the presence of a crowding agent (10% dextran). Representative images from three experiments are depicted as an overlay of differential interference contrast (DIC) and GFP. Scale bars, 10 μm.

**D)** Bar chart (top) representing quantification (n = 3) of the number of droplets per frame from the LLPS experiments with BARD1 BRCT w.t. shown in (C). Statistical significance was determined by unpaired t-test. A nested scatterplot (bottom) represents quantification (n = 3) of an area of individual droplets from three independent experiments with BARD1 BRCT w.t. shown in (C)., with median area determined per dataset. Statistical significance was determined by nested t-test.

**Supplementary Figure 11:**

**A)** Scan of representative gel from EMSA experiments performed with Cy3-labelled RNA and BRCA1 BRCT.

**B)** Scan of representative gels from EMSA experiments performed with Cy3-labelled RNA and BARD1 BRCT.

**C)** Scan of representative gels from EMSA experiments performed with Cy3-labelled RNA or Cy3-labelled dsDNA and BRCA1 BRCT or BRCA1 DNA-binding domain (DBD).

**D)** Scan of representative gels from competition EMSA experiments performed with Cy3-labelled RNA, Cy3-labelled ssDNA, and Cy3-labelled dsDNA, respectively, with BRCA1 BRCT and BARD1 BRCT, respectively, in the presence of pS5pS7 GST-$(CTD)_{26}$.

**Supplementary Figure 12:**

**A)** Graphs representing quantification of EMSA experiments (n = 3) performed with Cy3-labelled RNA and BRCA1 BRCT (left) and BARD1 BRCT (right), respectively. Statistical significance was determined using unpaired Student's t-test.

**B)** Bar charts representing the binding of RNA and double-stranded DNA to BRCA1 BRCT and BARD1 BRCT, respectively, in the presence of pS5pS7 GST-$(CTD)_{26}$. Statistical significance was determined using unpaired Student's t-test.

**C)** Graphs representing quantification of EMSA experiments (n = 3) comparing the binding affinity of BRCA1 BRCT and BRCA1 DNA-binding domain (DBD) to Cy3-labelled RNA and double-stranded DNA.

**D)** AlphaFold 3-generated models of RNA binding to the BRCA1 BRCT (left) and BARD1 BRCT (right). BRCA1 BRCT interacts with RNA through the phosphopeptide-binding site, whilst BARD1 BRCT through an alternative binding-site.

**E)** Detail of the RNA binding sites (bottom) of the BRCA1 BRCT (left) and BARD1 BRCT (right). The hydrogen bonds were displayed as pseudobonds using the structural analysis tool Hydrogen bonds in UCSF ChimeraX and are depicted in turquoise. The relax distance tolerance was 1.0 Å and the relax angle tolerance was 20.0°. Van der Waals, hydrophobic and stacking interactions (right) were displayed as pseudobonds using the structural analysis tool in UCSF ChimeraX and are depicted in black. The interacting atoms were identified based on VDW overlap ≥ -0.4 Å.

**Supplementary Figure 13:**

**A)** Comparison of the surface charges of BRCA1 BRCT (left) and BARD1 BRCT (right). The electrostatic potential was calculated by the Adaptive Poisson-Boltzmann Solver and visualised in Pymol.

**B)** Multiple sequence alignment of amino-acids present in putative RNA-binding sites of BARD1 BRCT domains from various organisms. The alignment was generated using Clustal Omega on EMBL-EBI and visualised using Jalview.

**Supplementary Figure 14:**

**A)** LLPS assays with purified, Alexa488-labelled BRCT domains and Cy5-ITS1 RNA. BRCA1 BRCT (left, at 10, 20, and 40 µM) and BARD1 BRCT (right, at 10, 20, and 40 µM), respectively, were mixed with ITS1 RNA (15nM) in the presence of a crowding agent (10% PEG 8000). Representative images from three experiments are depicted as an overlay of differential interference contrast (DIC), Alexa488, and Cy5 signal. Scale bars, 10 µm.

**B)** LLPS assays with purified, Alexa488-labelled BRCT domains and Cy5-ITS1 RNA. BRCA1 BRCT (left, w.t., and S1655F,K1702M, respectively, at 160 µM) and BARD1 BRCT (right, w.t.; S575F,K619A; and R705A, respectively, at 160 µM), respectively, were mixed with Cy5-labelled ITS1 RNA (15nM) in the presence of a crowding agent (10% dextran). Representative images from three experiments are depicted as an overlay of differential interference contrast (DIC), and Cy5 signal. Scale bars, 10 µm.

**C)** Effect of the Alexa-488 labelling on the LLPS propensity of BRCT domains. BRCA1 BRCT (left, non-labelled or Alexa488-labelled, at 160 µM) and BARD1 BRCT (right, non-labelled or Alexa488-labelled, at 160 µM), respectively, were mixed with Cy5-labelled ITS1 RNA (15nM) in the presence of a crowding agent (10% dextran). Representative images from three experiments are depicted as an overlay of differential interference contrast (DIC), and Cy5 signal. Scale bars, 10 µm.

**D)** Effect of the Alexa-488 labelling on the LLPS propensity of BRCT domains. Bar chart (top) representing quantification (n = 3) of the number of droplets per frame from the LLPS experiments with BRCT domains and ITS1 RNA shown in (C). Statistical significance was determined by unpaired t-test. A nested scatterplot (bottom) representing quantification (n = 3) of an area of individual droplets from three independent experiments with BRCT domains and ITS1 RNA shown in (C), with median area determined per dataset. Statistical significance was determined by nested t-test.

**Supplementary Figure 15:**

**A)** LLPS assays with purified, Alexa488-labelled BRCT domains and Cy5-ITS1 RNA. BRCA1 BRCT (left, at 40 µM) and BARD1 BRCT (right, at 40 µM), respectively, were mixed with ITS1 RNA (15nM) in the presence of a crowding agent (10% PEG 8000). Representative images from three experiments are depicted as an overlay of differential interference contrast (DIC), Alexa488 and Cy5 signal. Scale bars, 10 µm.

**B)** Bar chart (top) representing quantification (n = 3) of the number of droplets per frame from the LLPS experiments with BRCA1 BRCT (left) and BARD1 BRCT (right), shown in (A). Statistical significance was determined by unpaired t-test. A nested scatterplot (bottom) representing quantification (n = 3) of an area of individual droplets from three independent experiments with BRCA1 BRCT (left) and BARD1 BRCT (right), shown in (A), with median area determined per dataset. Statistical significance was determined by nested t-test.

**C)** Bar chart (top) representing quantification (n = 3) of the number of droplets per frame from the LLPS experiments with BRCT domains and ITS1 RNA. BRCA1 BRCT (left, w.t. and S1655,K1702M, at 160 µM) and BARD1 BRCT (right, w.t.; S575F,K619A; and R705A, respectively, at 160 µM), respectively, were mixed with Cy5-labelled ITS1 RNA (15nM) in the presence of a crowding agent (10% dextran). Statistical significance was determined by unpaired t-test. A nested scatterplot (bottom) representing quantification (n = 3) of an area of individual droplets from three independent experiments with BRCT domains and ITS1 RNA, with median area determined per dataset. Statistical significance was determined by nested t-test.

**Supplementary Figure 16:**

**A)** SDS-PAGE gels analysis of cross-linked BRCA1 BRCT and BARD1 BRCT samples, respectively, showing the multimerization of BRCT domains.

**B)** SDS-PAGE gel analysis of a sedimentation assays. BARD1 BRCT 2M denotes variant BARD1 BRCT$^{S575F,K619A}$, BRCA1 BRCT 2M denotes variant BRCA1 BRCT$^{S1655F,K1702M}$.

**C)** SAXS analysis-BRCA1 BRCT wt P(r) comparison (left) and BARDA1 BRCT wt P(r) comparison (right) showing increase of Maximum distance R with increasing concentration of the sample (2mg/ml – green, 3mg/ml – yellow, 4mg/ml – red).

**Supplementary Figure 17:**

**A)** LLPS assays with purified BRCA1 BRCT, pS5pS7 mGFP-(hCTD), and Cy5-ITS1 RNA. BRCA1 BRCT (at 10 μM) was mixed with phosphorylated pCTD (2.5 μM), and/or ITS1 RNA (15nM) in the presence of a crowding agent (10% PEG 8000). Where indicated, $MgCl_2$ was present (at 3mM). Representative images from three experiments are depicted as an overlay of differential interference contrast (DIC), GFP and Cy5 signal. Scale bars, 10 μm.

**B)** Bar chart (top) representing quantification (n = 3) of the number of droplets per frame from the LLPS experiments with BRCA1 BRCT, pS5pS7 mGFP-hCTD, and Cy5-ITS1 RNA, shown in (A), using the green (left) or red (right) fluorescent signal. Statistical significance was determined by unpaired t-test. A nested scatterplot (bottom) representing quantification (n = 3) of an area of individual droplets from three independent experiments with BRCA1 BRCT, pS5pS7 mGFP-hCTD, and Cy5-ITS1 RNA, shown in (A), with median area determined per dataset. Statistical significance was determined by nested t-test.

**C)** LLPS assays with purified BARD1 BRCT, pS5pS7 mGFP-hCTD, and Cy5-ITS1 RNA. BARD1 BRCT (at 10 μM) was mixed with phosphorylated CTD (2.5 μM), and/or ITS1 RNA (15nM) in the presence of a crowding agent (10% PEG 8000). Where indicated, $MgCl_2$ was present (at 3mM). Representative images from three experiments are depicted as an overlay of differential interference contrast (DIC), GFP, and Cy5 signal. Scale bars, 10 μm.

**D)** Bar chart (top) representing quantification (n = 3) of the number of droplets per frame from the LLPS experiments with BARD1 BRCT, pS5pS7 mGFP-hCTD, and Cy5-ITS1 RNA, shown in (C), using the green (left) or red (right) fluorescent signal. Statistical significance was determined by unpaired t-test. A nested scatterplot (bottom) representing quantification (n = 3) of an area of individual droplets from three independent experiments with BARD1 BRCT, pS5pS7 mGFP-hCTD, and Cy5-ITS1 RNA, shown in (C), with median area determined per dataset. Statistical significance was determined by nested t-test.

**Supplementary Figure 18:**

**A)** LLPS assays with purified BRCA1 BRCT, pS5pS7 mGFP-hCTD, and Cy5-ITS1 RNA. BRCA1 BRCT (w.t. and S1655F,K1702M, at 10 μM) was mixed with phosphorylated CTD (2.5 μM), and ITS1 RNA (15nM) in the presence of a crowding agent (10% PEG 8000). Representative images from three experiments are depicted as an overlay of differential interference contrast (DIC), GFP, and Cy5 signal. Scale bars, 10 μm.

**B)** Bar chart (top) representing quantification (n = 3) of the number of droplets per frame from the LLPS experiments with BRCA1 BRCT, pS5pS7 mGFP-hCTD, and Cy5-ITS1 RNA, shown in (A), using the green or red fluorescent signal. Statistical significance was determined by unpaired t-test. A nested scatterplot (bottom) representing quantification (n = 3) of an area of individual droplets from three independent experiments with BRCA1 BRCT, pS5pS7 mGFP-hCTD, and Cy5-ITS1 RNA, shown in (A)., with median area determined per dataset. Statistical significance was determined by nested t-test.

**C)** LLPS assays with purified BARD1 BRCT, pS5pS7 mGFP-hCTD, and Cy5-ITS1 RNA. BRCA1 BRCT (w.t.; S575FK619A; and R705A, respectively, at 10 μM) was mixed with phosphorylated CTD (2.5 μM), and ITS1 RNA (15nM) in the presence of a crowding agent (10% PEG 8000). Representative images from three experiments are depicted as an overlay of differential interference contrast (DIC), GFP, and Cy5 signal. Scale bars, 10 μm.

**D)** Bar chart (top) representing quantification (n = 3) of the number of droplets per frame from the LLPS experiments with BARD1 BRCT, pS5pS7 mGFP-hCTD, and Cy5-ITS1 RNA, shown in (C), using the green or red fluorescent signal. Statistical significance was determined by unpaired t-test. A nested scatterplot (bottom) representing quantification (n = 3) of an area of individual droplets from three independent experiments with BARD1 BRCT, pS5pS7 mGFP-hCTD, and Cy5-ITS1 RNA, shown in (C), with median area determined per dataset. Statistical significance was determined by nested t-test.

**E)** Sensograms obtained by biolayer-interferometry (BLI) (left). Comparison of association ($K_{as}$), dissociation ($K_{dis}$), and dissociation ($K_D$) constants for pS5pS7 GST-$(CTD)_{26}$ and BARD1 BRCT w.t. and BARD1 BRCT$^{R705A}$, respectively, obtained by biolayer interferometry (right). The sensograms represent the mean of 3 measurements for each concentration. The data were analysed in Octet® Analysis Studio Software using 1:2 Bivalent analyte model. The data were plotted using Prism GraphPad 9 software.

**F)** Comparison of sensograms obtained by biolayer-interferometry (BLI) with BARD1 BRCT w.t., and BARD1 BRCT$^{R705A}$, and their respective fits (left). Comparison of association ($K_{as}$), dissociation ($K_{dis}$), and dissociation ($K_D$) constants for pS5pS7 GST-$(CTD)_{26}$ and BARD1 BRCT w.t., and BARD1 BRCT$^{R705A}$, respectively, obtained by biolayer interferometry (right).

**Supplementary Figure 19:**

**A)** Conservation of the selected positions in BRCA1 BRCT that were mutated in cancer. Multiple sequence alignment of sequences of BRCA1 BRCT domains from various organisms was generated using Clustal Omega on EMBL-EBI and visualised using Jalview.

**B)** Conservation of the selected positions in BARD1 BRCT that were mutated in cancer. Multiple sequence alignment of sequences of BARD1 BRCT domains from various organisms was generated using Clustal Omega on EMBL-EBI and visualised using Jalview.

**Supplementary Figure 20:**

Quantification of the thermal stability of BRCA1 BRCT (w.t., E1682K, R1699L, R1699W, and E1754K) and BARD1 BRCT (w.t., E587K, E665K, S711R, and K754N) measured by nanoDSF. The bar charts depict the temperatures corresponding to the onset of protein unfolding ($T_o$), the melting temperature ($T_m$), and the onset of aggregation ($T_{agg}$) from n = 9 measurements. Statistical significance was determined by unpaired t-test.

**Supplementary Figure 21:**

**A)** Sensograms obtained by biolayer-interferometry (BLI) with BRCA1 BRCT$^{E1754K}$ . The sensograms represent the mean of 3 measurements for each concentration. The data were analysed in Octet® Analysis Studio Software using 1:2 Bivalent analyte model. The data were plotted using Prism GraphPad 9 software.

**B)** Comparison of sensograms obtained by biolayer-interferometry (BLI) with BRCA1 BRCT w.t. and BRCA1 BRCT$^{E1754K}$ and their respective fits (left). Comparison of association ($K_{as}$), dissociation ($K_{dis}$), and dissociation ($K_D$) constants for pS5pS7 GST-(CTD)$_{26}$ and BRCA1 BRCT w.t. and BRCA1 BRCT$^{E1754K}$, respectively, obtained by biolayer interferometry (right). Cancer-associated substitution BRCA1 BRCT$^{E1754K}$ does not affect the binding of the pS5pS7 GST-(CTD)$_{26}$.

**C)** Sensograms obtained by biolayer-interferometry (BLI) with BARD1 BRCT$^{E665K}$ (left) and BARD1 BRCT$^{K754N}$ (right). The sensograms represent the mean of 3 measurements for each concentration. The data were analysed in Octet® Analysis Studio Software using 1:2 Bivalent analyte model. The data were plotted using Prism GraphPad 9 software.

**D)** Comparison of sensograms obtained by biolayer-interferometry (BLI) with BARD1 BRCT w.t., BARD1 BRCT$^{E665K}$, and BARD1 BRCT$^{K754N}$ and their respective fits (left). Comparison of association ($K_{as}$), dissociation ($K_{dis}$), and dissociation ($K_D$) constants for pS5pS7 GST-(CTD)$_{26}$ and BARD1 BRCT w.t., BARD1 BRCT$^{E665K}$, and BARD1 BRCT$^{K754N}$, respectively, obtained by biolayer interferometry (right). Cancer-associated substitutions BARD1 BRCT$^{E665K}$, and BARD1 BRCT$^{K754N}$ do not affect the binding of the pS5pS7 GST-(CTD)$_{26}$.

**Supplementary Figure 22:**

**A)** Bar charts (top) representing quantification (n = 3) of the number of droplets per frame from the LLPS experiments with BRCA1 BRCT variants E1682K, and E1754K shown in (Fig. 5B). Statistical significance was determined by unpaired t-test. Nested scatterplots (bottom) representing quantification (n = 3) of an area of individual droplets from three independent experiments with BRCA1 BRCT variants E1682K and E1754K, respectively, shown in (Fig. 5B), with median area determined per dataset. Statistical significance was determined by nested t-test.

**B)** Bar charts (top) representing quantification (n = 3) of the number of droplets per frame from the LLPS experiments with BARD1 BRCT variants E587K, E665K, S711R, and K754N shown in (Fig. 5C). Statistical significance was determined by unpaired t-test. Nested scatterplots (bottom) representing quantification (n = 3) of an area of individual droplets from three independent experiments with BARD1 BRCT variants E587K, E665K, S711R, and K754N, respectively, shown in (Fig. 5C), with median area determined per dataset. Statistical significance was determined by nested t-test.

**Supplementary Figure 23:**

**A)** LLPS assays with purified, Alexa488-labelled BRCT domains. BRCA1 BRCT (left, at 40 µM, 80 µM, 160 µM), w. t., and E1754K, respectively, was mixed with a crowding agent (10% dextran). BARD1 BRCT (right, at 40 µM, 80 µM, 160 µM), w. t., and K754N, respectively, was mixed with a crowding agent (10% dextran). Representative images from three experiments are depicted as an overlay of differential interference contrast (DIC) and Alexa488 signal. Scale bars, 10 µm.

**B)** Aggregation of purified, Alexa-488 labelled BARD1 BRCT$^{E665K}$. BARD1 BRCT$^{E665K}$ (160 µM), w. t., and K754N, respectively, was mixed with a crowding agent (10% dextran). Representative image from three experiments is depicted as an overlay of differential interference contrast (DIC) and Alexa488 signal. Scale bar, 10 µm.

**C)** SDS-PAGE gels with cross-linked samples of BRCA1 BRCT and BARD1 BRCT disease-associated mutants, comparing the multimerization of w.t. proteins and their respective disease-associated variants.

**D)** Bar charts (top) representing quantification (n = 3) of the number of droplets per frame from the LLPS experiments with BRCA1 BRCT (w.t. and E1754K, left) and BARD1 BRCT (w.t. and K754N, right) shown in (A). Statistical significance was determined by unpaired t-test. Nested scatterplots (bottom) representing quantification (n = 3) of an area of individual droplets from three independent experiments with BRCA1 BRCT (w.t. and E1754K, left) and BARD1 BRCT (w.t. and K754N, right) shown in (A), with median area determined per dataset. Statistical significance was determined by nested t-test.

**Supplementary Table 1: List of plasmids used in this study.**

| Name | Addgene # | Plasmid | Tags / Fusion proteins | Bacterial resistance |
|---|---|---|---|---|
| 2AT | 29665 | pET LIC cloning vector | - | Ampicilin |
| 2BT | 29666 | pET His6 TEV LIC cloning vector | His6-TEV (N-terminal on backbone) | Ampicilin |
| 2CT-10 | 55209 | pET His10 MBP Asn10 TEV LIC cloning vector | His10-MBP-N10-TEV (N-terminal on backbone) | Ampicilin |
| 2Bc-T | 37236 | pET His6 LIC cloning vector | TEV-His6 (C-terminal on backbone) | Ampicilin |
| 2E | 29775 | pET empty polycistronic destination vector | - | Kanamycin |
| 438-A | 55218 | pFastBac cloning vector with BioBrick PolyPromoter LIC Subcloning | - | Ampicilin |
| 438-B | 55219 | pFastBac His6 TEV cloning vector with BioBrick PolyPromoter LIC Subcloning | His6-TEV (N-terminal on backbone) | Ampicilin |
| H6-msfGFP | 29725 | pET Biotin His6 GFP LIC cloning vector | Biotin- His6-TEV (N terminal on backbone), GFP (C-terminal on backbone) | Kanamycin |

**Supplementary Table 2: List of oligonucleotides used in this study.**

| Name | Sequence | Use |
|---|---|---|
| T7F | TAATACGACTCACTATAGGG | Sequencing |
| T7R | GCTAGTTATTGCTCAGCGG | Sequencing |
| M13F | CCCAGTCACGACGTTGTAAAACG | Bacmid screening |
| M13R | AGCGGATAACAATTTCACACAGG | Bacmid screening |
| FB01 | CCTATAACTATTCCGGATTATTCATACCGTC | Sequencing |
| FB02 | CAGGTTCAGGGGGAGGTGTG | Sequencing |
| pRVK007 | AAGAATTGTTACAAATCACC | Sequencing of BRCA1 |
| pRVK008 | TGGATAACACTAAATAGC | Sequencing of BRCA1 |
| pRVK009 | ATAAGCAATATGGAACTCG | Sequencing of BRCA1 |
| pRVK010 | GAACCAAATAAATGTGTG | Sequencing of BRCA1 |
| pRVK011 | AGAACATTCCAAGTACAG | Sequencing of BRCA1 |
| pRVK012 | CAATATACCTTCTCAGTC | Sequencing of BRCA1 |
| pRVK013 | AGAAAAGTAGTGAATACC | Sequencing of BRCA1 |
| pRVK014 | TACTTCCAATCCAATGCAATGCCGGATAATCGGCAG | Forward primer for cloning of BARD1 into 438B, v1 tag |
| pRVK015 | TTATCCACTTCCAATGTTATTATTAGCTGTCAAGAGGAAGCA | Reverse primer for cloning of BARD1 into 438B, v1 tag |
| pRVK016 | GAAAGTCAGATATGTTGTG | Sequencing of BARD1 |

| | | |
|---|---|---|
| pRVK017 | GTGGAGATTTTGTTAAGC | Sequencing of BARD1 |
| pRVK018 | TACTTCCAATCCAATCGATGGACTACAAGGACGATGACGACAAGGGAAGTGGTATGGATTTATCTGCTCTTCGC | Forward primer for cloning of BRCA1 into 438A, vBac F tag, FLAG tag |
| pRVK019 | TTATCCACTTCCAATGTTATTATCAGTAGTGGCTGTGGGG | Reverse primer for cloning of BRCA1 into 438A, v1 tag |
| pRVK032 | TCTGTAGAAGTCTTTTGG | Sequencing of BRCA1, reverse |
| pRVK041 | TACTTCCAATCCAATGCAGTCAACAAAAGAATGTCC | Forward primer for cloning of BRCA1 $^{1646\text{-}1859}$ into 2BT, v1 tag |
| pRVK042 | TTATCCACTTCCAATGTTATTATCAGGGGATCTGGGGTAT | Reverse primer for cloning of BRCA1 $^{1646\text{-}1859}$ into 2BT, v1 tag |
| pRVK048 | TACTTCCAATCCAATGCAGCTAGCCACTGCTCAGTA | Forward primer for cloning of BARD1 $^{554\text{-}777}$ into 2BT, v1 tag |
| pRVK064 | CAAAAGAATGTCCATGGTGGTGTTTGGCCTGACCCCAGAAGAATTTATG | Forward primer for site-directed mutagenesis to introduce a S1655F substitution into BRCA1 |
| pRVK065 | CATAAATTCTTCTGGGGTCAGGCCAAACACCACCATGGACATTCTTTTG | Reverse primer for site-directed mutagenesis to introduce a S1655F substitution into BRCA1 |
| pRVK066 | GTTTGTGTGTGAACGGACACTGATGTATTTTCTAGGAATTGCGGGAGG | Forward primer for site-directed mutagenesis to introduce a K1702M substitution into BRCA1 |
| pRVK067 | CCTCCCGCAATTCCTAGAAAATACATCAGTGTCCGTTCACACACAAAC | Reverse primer for site-directed mutagenesis to introduce a K1702M substitution into BRCA1 |
| pRVK068 | GATGGACCTCTTGTACTTATAGGCTTTGGGCTGTCTTCAGAACAACAG | Forward primer for site-directed mutagenesis to introduce a S575F substitution into BARD1 |
| pRVK069 | CTGTTGTTCTGAAGACAGCCCAAAGCCTATAAGTACAAGAGGTCCATC | Reverse primer for site-directed mutagenesis to introduce a S575F substitution into BARD1 |
| pRVK070 | GGTGATGCAGTTCAAAGTACCTTGGCGTGTATGCTTGGGATTCTCAATGG | Forward primer for site-directed mutagenesis to introduce a K619M substitution into BARD1 |
| pRVK071 | CCATTGAGAATCCCAAGCATACACGCCAAGGTACTTTGAACTGCATCACC | Reverse primer for site-directed mutagenesis to introduce a K619M substitution into BARD1 |
| pRVK072 | TTTAAGAAGGAGATATAGTTCATGGACTACAAAGACGATGACGACAAGGAAAACCTGTACTTCCAATCCAATATGGATTTATCTGCTCTTCG | Forward primer for cloning of BRCA1 into 2BcT, FLAG tag, v3 tag |
| pRVK073 | GGATTGGAAGTAGAGGTTCTCTCAGTAGTGGCTGTGGGG | Reverse primer for cloning of BRCA1 into 2BcT, v3 tag |
| pRVK074 | TACTTCCAATCCAATCGATGGACTACAAAGACGATGAC | Forward primer for cloning of N-terminally FLAG-tagged sequences into 438 plasmids, vBAC F tag |
| pRVK075 | TTATCCACTTCCAATGTTATTAATGGTGATGGTGATGGTG | Reverse primer for cloning of C-terminally His-tagged sequences into 438 plasmids, v1 tag |
| pRVK087 | TACTTCCAATCCAATGCATATTCTGGTTCTTCAGAGAAA | Forward primer for cloning cloning of BRCA1$^{421\text{ - }1079}$ into 438C, v1 tag |
| pRVK092 | TTATCCACTTCCAATGTTATTACAATTTTGGCCCTCTGTTTCT | Reverse primer for cloning of BRCA1 $^{421\text{ - }1079}$ into 438C, v1 tag |
| pRVK093 | ACTGAGCCACAGATAATACAAGAGCATCCCCTCACAAATAAATTAAAGCGT | Forward primer for site-directed mutagenesis to introduce a R496H substitution into BRCA1 |
| pRVK094 | ACGCTTTAATTTATTTGTGAGGGGATGCTCTTGTATTATCTGTGGCTCAGT | Reverse primer for site-directed mutagenesis to introduce a R496H substitution into BRCA1 |

| | | |
|---|---|---|
| pRVK095 | TACTGAGCCACAGATAATACAAGAGTGTCCCCTCACAAATAAATTAAAGC | Forward primer for site-directed mutagenesis to introduce a R496C substitution into BRCA1 |
| pRVK096 | GCTTTAATTTATTTGTGAGGGGACACTCTTGTATTATCTGTGGCTCAGTA | Reverse primer for site-directed mutagenesis to introduce a R496C substitution into BRCA1 |
| pRVK097 | CAGCAGTATAAGCAATATGGAACTCAAATTAAATATCCACAATTCAAAAG | Forward primer for site-directed mutagenesis to introduce a E597K substitution into BRCA1 |
| pRVK098 | CTTTTGAATTGTGGATATTTAATTTGAGTTCCATATTGCTTATACTGCTG | Reverse primer for site-directed mutagenesis to introduce a E597K substitution into BRCA1 |
| pRVK099 | GCTGAGGAGGAAGTCTTCTACCAGGAATATTCATGCGCTTGAACTAGTAG | Forward primer for site-directed mutagenesis to introduce a H619N substitution into BRCA1 |
| pRVK100 | CTACTAGTTCAAGCGCATGAATATTCCTGGTAGAAGACTTCCTCCTCAGC | Reverse primer for site-directed mutagenesis to introduce a H619N substitution into BRCA1 |
| pRVK101 | CAAGGGACTAATTCATGGTTGTTCCGAAGATAATAGAAATGACACAGAAG | Forward primer for site-directed mutagenesis to introduce a K820E substitution into BRCA1 |
| pRVK102 | CTTCTGTGTCATTTCTATTATCTTCGGAACAACCATGAATTAGTCCCTTG | Reverse primer for site-directed mutagenesis to introduce a K820E substitution into BRCA1 |
| pRVK103 | ATTGGGACATGAAGTTAACCACAGTTGGGAAACAAGCATAGAAATGGAAG | Forward primer for site-directed mutagenesis to introduce a R841W substitution into BRCA1 |
| pRVK104 | CTTCCATTTCTATGCTTGTTTCCCAACTGTGGTTAACTTCATGTCCCAAT | Reverse primer for site-directed mutagenesis to introduce a R841W substitution into BRCA1 |
| pRVK110 | CAGATGCTGAGTTTGTGTGTGAATGGACACTGAAATATTTTCTAGGAATTGC | Forward primer for site-directed mutagenesis to introduce a R1699W substitution into BRCA1 |
| pRVK111 | GCAATTCCTAGAAAATATTTCAGTGTCCATTCACACACAAACTCAGCATCTG | Reverse primer for site-directed mutagenesis to introduce a R1699W substitution into BRCA1 |
| pRVK112 | CAGATGCTGAGTTTGTGTGTGAACTGACACTGAAATATTTTCTAGGAATTGC | Forward primer for site-directed mutagenesis to introduce a R1699L substitution into BRCA1 |
| pRVK113 | GCAATTCCTAGAAAATATTTCAGTGTCAGTTCACACACAAACTCAGCATCTG | Reverse primer for site-directed mutagenesis to introduce a R1699L substitution into BRCA1 |
| pRVK114 | GCAGGTGGGGGCCAGATCCTCAGTGCAAAGCCCAAGCCAGACAGTGACGTG | Forward primer for site-directed mutagenesis to introduce a R705A substitution into BARD1 |
| pRVK115 | CACGTCACTGTCTGGCTTGGGCTTTGCACTGAGGATCTGGCCCCCACCTGC | Reverse primer for site-directed mutagenesis to introduce a R705A substitution into BARD1 |
| pRVK128 | CAGAACAACAGAAAATGCTCAGTAAGCTTGCAGTAATTCTTAAGGC | Forward primer for site-directed mutagenesis to introduce a E587K substitution into BARD1 |
| pRVK129 | GCCTTAAGAATTACTGCAAGCTTACTGAGCATTTTCTGTTGTTCTG | Reverse primer for site-directed mutagenesis to introduce a E587K substitution into BARD1 |
| pRVK130 | CCACGCAGAAGCAGGCTCAACAGAAAACAGCTGTTGCCAAAGCTGTTTG | Forward primer for site-directed mutagenesis to introduce a E665K substitution into BARD1 |
| pRVK131 | CAAACAGCTTTGGCAACAGCTGTTTTCTGTTGAGCCTGCTTCTGCGTGG | Reverse primer for site-directed mutagenesis to introduce a E665K substitution into BARD1 |
| pRVK132 | CAGTAGAAAGCCCAAGCCAGACCGTGACGTGACTCAGACCATCAATAC | Forward primer for site-directed mutagenesis to introduce a S711R substitution into BARD1 |

| | | |
|---|---|---|
| pRVK133 | GTATTGATGGTCTGAGTCACGTCACGGTCTGGCTTGGGCT TTCTACTG | Reverse primer for site-directed mutagenesis to introduce a S711R substitution into BARD1 |
| pRVK134 | CAGAGAGGGTTCGGCAGGGCAACGTCTGGAAGGCTCCTT CGAGCTG | Forward primer for site-directed mutagenesis to introduce a K754N substitution into BARD1 |
| pRVK135 | CAGCTCGAAGGAGCCTTCCAGACGTTGCCCTGCCGAACCC TCTCTG | Reverse primer for site-directed mutagenesis to introduce a K754N substitution into BARD1 |
| pRVK139 | TTTAAGAAGGAGATATAGATCATGGGAGGCCTGAACGAT | Forward primer for cloning of Avi-tagged sequences into 2AT, v2 tag |
| pRVK140 | TTATGGAGTTGGGATCTTATTAGCTGTCAAGAGGAAGCAA | Reverse primer for cloning of BARD1 into 2AT, v2 tag |
| pRVK141 | TACTTCCAATCCAATGCAAAGGATAACACCGTGCCA | Forward primer for cloning of BirA into 2CT10, v1 tag |
| pRVK142 | TTATCCACTTCCAATGTTATTATTTTCTGCACTACGCAG | Reverse primer for cloning of BirA into 2CT10, v1 tag |
| pRVK159 | GTTGCTATGGGCCCTTCACCAACAAGCCCACAGATCAACT GGAATGGATG | Forward primer for site-directed mutagenesis to introduce a M1775K substitution into BRCA1 |
| pRVK160 | CATCCATTCCAGTTGATCTGTGGGCTTGTTGGTGAAGGGC CCATAGCAAC | Reverse primer for site-directed mutagenesis to introduce a M1775K substitution into BRCA1 |
| pRVK161 | GTTGCTATGGGCCCTTCACCAACCACCCCACAGATCAACT GGAATGGATG | Forward primer for site-directed mutagenesis to introduce a M1775H substitution into BRCA1 |
| pRVK162 | CATCCATTCCAGTTGATCTGTGGGGTGGTTGGTGAAGGGC CCATAGCAAC | Reverse primer for site-directed mutagenesis to introduce a M1775H substitution into BRCA1 |
| pRVK163 | CAGATGCTGAGTTTGTGTGTGAATCGACACTGAAATATTT TCTAGGAATTGC | Forward primer for site-directed mutagenesis to introduce a R1699S substitution into BRCA1 |
| pRVK164 | GCAATTCCTAGAAAATATTTCAGTGTCGATTCACACACAA ACTCAGCATCTG | Reverse primer for site-directed mutagenesis to introduce a R1699S substitution into BRCA1 |
| pRMS217 | GACGCTGCCGAATTCTACCAGTGCCTTGCTA GGACATCTTTGCCCACCTGCAGGTTCACCC | 61-mer DNA primer fluorescently labelled on 5’ end with Cy3 used to create R-loop/D-loop/bubble substrates for EMSAs |
| pRMS219 | GGGTGAACCTGCAGGTGGGCGGCTGCTCAT CGTAGGTTAGTTGGTAGAATTCGGCAGCGTC | 61-mer DNA primer used to create R-loop/D-loop/bubble substrates for EMSAs and helicase assays |
| pRMS223 | AAAGAUGUCCUAGCAAGGCAC | 21-mer RNA primer used to create an R-loop substrate for EMSAs and helicase assays |
| pRMS391 | GGGTGAACCTGCAGGTGGGCAAAGATGTCC TAGCAAGGCACTGGTAGAATTCGGCAGCGT C | 61-mer DNA primer used to create a double stranded DNA substrate for EMSA and helicase assays |
| pRMS433 | CACTTTAACTAATCTAATTACTAAAGAGACTACTCATGTT GTTATG | Forward primer for site-directed mutagenesis to introduce a E1682K substitution into BRCA1 |
| pRMS434 | CATAACAACATGAGTAGTCTCTTTAGTAATTAGATTAGTT AAAGTG | Reverse primer for site-directed mutagenesis to introduce a E1682K substitution into BRCA1 |
| pRMS435 | CCAAGGTCCAAAGCGAGCAAGAAAATCCCAGGACAGAAA GATCTTC | Forward primer for site-directed mutagenesis to introduce a E1754K substitution into BRCA1 |
| pRMS436 | GAAGATCTTTCTGTCCTGGGATTTTCTTGCTCGCTTTGGAC CTTGG | Reverse primer for site-directed mutagenesis to introduce a E1754K substitution into BRCA1 |
| pRMS437 | CCAAGGTCCAAAGCGAGCAAGAGAATACCAGGACAGAAA GATCTTCAG | Forward primer for site-directed mutagenesis to introduce a S1755Y substitution into BRCA1 |

| | | |
|---|---|---|
| pRMS438 | CTGAAGATCTTTCTGTCCTGGTATTCTCTTGCTCGCTTTGG ACCTTGG | Reverse primer for site-directed mutagenesis to introduce a S1755Y substitution into BRCA1 |
| pRMS679 | TTGTCTTCTCATAAATTTAATCCCCGTACGCTTATACTCCT TTAACTACCCAGTCTTCCCG | 61-mer DNA primer fluorescently labelled on 5’ end with Cy3 used as a single-stranded DNA for EMSAs |

**Supplementary Table 3: List of recombinant plasmids used in this study.**

| Name of the plasmid | Backbone | ORF | Boundaries (AA) | Purpose |
|---|---|---|---|---|
| pVK1.1 | 438B | BARD1 | 1-777 (full-length) | Transfer vector / cloning intermediate |
| pVK1.2 | 2BT | BRCA1 | 1646-1859 | CTD binding assays, LLPS experiments |
| pVK1.3 | 2BT | BARD1 | 554-777 | CTD binding assays, LLPS experiments |
| pVK1.4 | 2BT | BRCA1 S1655F | 1646-1859 | Transfer vector / cloning intermediate |
| pVK1.5 | 2BT | BRCA1 K1702M | 1646-1859 | Transfer vector / cloning intermediate |
| pVK1.6 | 2BT | BARD1 S575F | 554-777 | Transfer vector / cloning intermediate |
| pVK1.7 | 2BT | BARD1 K619A | 554-777 | Transfer vector / cloning intermediate |
| pVK1.8 | 2BT | BRCA1 S1655F, K1702M | 1646-1859 | CTD binding assays, LLPS experiments |
| pVK1.9 | 2BT | BARD1 S575F, K619A | 554-777 | CTD binding assays, LLPS experiments |
| pVK1.10 | 2BT | BRCA1 E1682K | 1646-1859 | CTD binding assays, LLPS experiments |
| pVK1.11 | 2BT | BRCA1 E1754K | 1646-1859 | CTD binding assays, LLPS experiments |
| pVK1.12 | 2BT | BRCA1 S1755Y | 1646-1859 | CTD binding assays, LLPS experiments |
| pVK1.13 | 2BcT | FLAG-BRCA1 | 1-1859 (full-length) | Transfer vector / cloning intermediate |
| pVK1.14 | 438A | FLAG-BRCA1-His | 1-1859 (full-length) | Transfer vector / cloning intermediate |
| pVK1.15 | 438A | FLAG-BRCA1-His, His-BARD1 | BRCA1 1-1859 (full-length), BARD1 1-777 (full-length) | Structural studies, biochemical characterization |
| pVK1.16 | 438C | BRCA1 | 421-1079 | DNA binding assays |
| pVK1.17 | 438C | BRCA1 R496H | 421-1079 | DNA binding assays |
| pVK1.18 | 438C | BRCA1 R496C | 421-1079 | DNA binding assays |
| pVK1.19 | 438C | BRCA1 E597K | 421-1079 | DNA binding assays |
| pVK1.20 | 438C | BRCA1 H619N | 421-1079 | DNA binding assays |
| pVK1.21 | 438C | BRCA1 R841W | 421-1079 | DNA binding assays |
| pVK1.22 | 2BT | BRCA1 R1699W | 1646-1859 | CTD binding assays |
| pVK1.23 | 2BT | BRCA1 R1699L | 1646-1859 | CTD binding assays |
| pVK1.24 | 2BT | BARD1 R705A | 554-777 | CTD binding assays |
| pVK1.25 | 2BT | BARD1 E587K | 554-777 | CTD binding assays, LLPS experiments |
| pVK1.26 | 2BT | BARD1 E665K | 554-777 | CTD binding assays, LLPS experiments |

| | | | | |
|---|---|---|---|---|
| pVK1.27 | 2BT | BARD1 S711R | 554-777 | CTD binding assays, LLPS experiments |
| pVK1.28 | 2BT | BARD1 K754N | 554-777 | CTD binding assays, LLPS experiments |
| pVK1.29 | H6-msfGFP | BARD1 | 554-777 | Transfer vector / cloning intermediate |
| pVK1.30 | H6-msfGFP | BARD1 S575F, K619A | 554-777 | Transfer vector / cloning intermediate |
| pVK1.31 | H6-msfGFP | BARD1 R705A | 554-777 | Transfer vector / cloning intermediate |
| pVK1.32 | 2AT | BARD1 | 554-777 | CTD binding assays |
| pVK1.33 | 2AT | BARD1 S575F, K619A | 554-777 | CTD binding assays |
| pVK1.34 | 2AT | BARD1 R705A | 554-777 | CTD binding assays |
| pVK1.35 | 2CT10 | BirA | 1-321 (full-length) | Transfer vector / cloning intermediate |
| pVK1.36 | 2AT | His-MBP-BirA | 1-321 (full-length) | Biotinylation |
| pVK1.37 | H6-msfGFP | BRCA1 | 1646-1859 | Transfer vector / cloning intermediate |
| pVK1.38 | H6-msfGFP | BRCA1 S1655F, K1702M | 1646-1859 | Transfer vector / cloning intermediate |
| pVK1.39 | H6-msfGFP | BRCA1 R1699L | 1646-1859 | Transfer vector / cloning intermediate |
| pVK1.40 | H6-msfGFP | BRCA1 E1754K | 1646-1859 | Transfer vector / cloning intermediate |
| pVK1.41 | 2AT | Avi-His-BRCA1 | 1646-1859 | CTD binding assays |
| pVK1.42 | 2AT | Avi-His-BRCA1 S1655F, K1702M | 1646-1859 | CTD binding assays |
| pVK1.43 | 2AT | Avi-His-BRCA1 R1699L | 1646-1859 | CTD binding assays |
| pVK1.44 | 2AT | Avi-His-BRCA1 E1754K | 1646-1859 | CTD binding assays |
| pVK1.45 | 2AT | Avi-His-BARD1 E665K | 554-777 | CTD binding assays |
| pVK1.46 | 2AT | Avi-His-BARD1 K754N | 554-777 | CTD binding assays |
| pVK1.47 | 2BT | BRCA1 M1775K | 1646-1859 | CTD binding assays |
| pVK1.48 | 2BT | BRCA1 M1775H | 1646-1859 | CTD binding assays |
| pVK1.49 | 2AT | Avi-His-BRCA1 M1775H | 1646-1859 | CTD binding assays |
| pVK1.50 | 2BT | BRCA1 R1699S | 1646-1859 | CTD binding assays |
| pVK1.51 | 2AT | Avi-His-BRCA1 R1699S | 1646-1859 | CTD binding assays |
| pVK1.52 | 438B | CDK7, MAT1, CCNH | full-length proteins | phosphorylation of CTD |
| pVK1.53 | 2BcT | mCherry-RPB1 CTD | 1593-1970 | LLPS experiments |

**Supplementary Table 4: Data collection and refinement statistics. Data in parentheses for highest resolution shell.**

| | |
|---|---|
| ***Data collection*** | |
| Complex | BRCA1/2xCTD |
| Beamline, diffraction source | P13, PETRA III |
| Wavelength (Å) | 0.9763 |
| Space group | $C222_1$ |
| a, b, c (Å) | 112.83, 134.76, 180.38 |
| α, β, γ (°) | 90.00, 90.00, 90.00 |
| No. of monomers in asymmetric unit | 4 |
| Resolution range (Å) | 49.37–3.00 (2.85–2.70) |
| Total no. of reflections | 382 280 (56 707) |
| No. of unique reflections | 27 850 (4 024) |
| Completeness (%) | 99.8 (99.9) |
| Redundancy | 13.7 (14.1) |
| ⟨ I/σ(I)⟩ | 7.2 (1.0) |
| $R_{merge}$ (%) | 0.289 (3.230) |
| CC(1/2) | 0.996 (0.505) |
| Wilson B ($Å^2$) | 76.3 |
| ***Refinement statistics*** | |
| No. of protein amino acids | 845 |
| No. of protein atoms | 6609 |
| No. of peptide-ligand atoms | 210 |
| Resolution limits | 49.42–3.00 (3.08–3.00) |
| No. of reflections in working set | 26 437 (1938) |
| No. of reflections in test set | 1386 (89) |
| Final $R_{cryst}$ (%) | 0.255 (0.498) |
| Final $R_{free}$ (%) | 0.312 (0.544) |
| Mean B factor ($Å^2$) | 103.0 |
| R.m.s. deviations | |
| Bonds (Å) | 0.004 |
| Angles (°) | 1.257 |
| Planar groups (Å) | 0.005 |
| Chiral volumes ($Å^3$) | 0.064 |
| Ramachandran plot | |
| Most favoured (%) | 93.2 |
| Allowed (%) | 6.8 |

**Supplementary figure 1** **Klapstova *et al.*,**

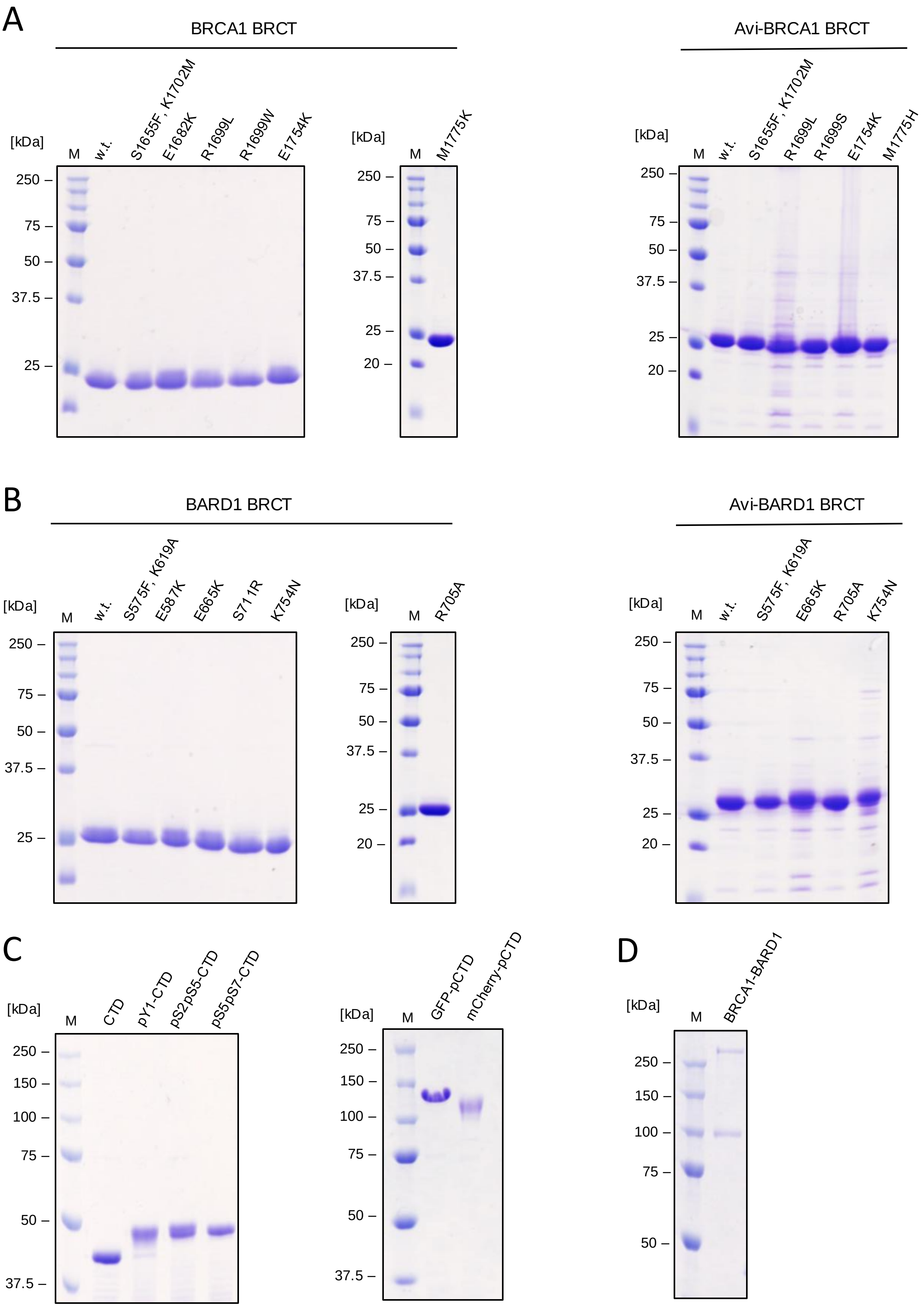

**Supplementary figure 2** Klapstova *et al.*,

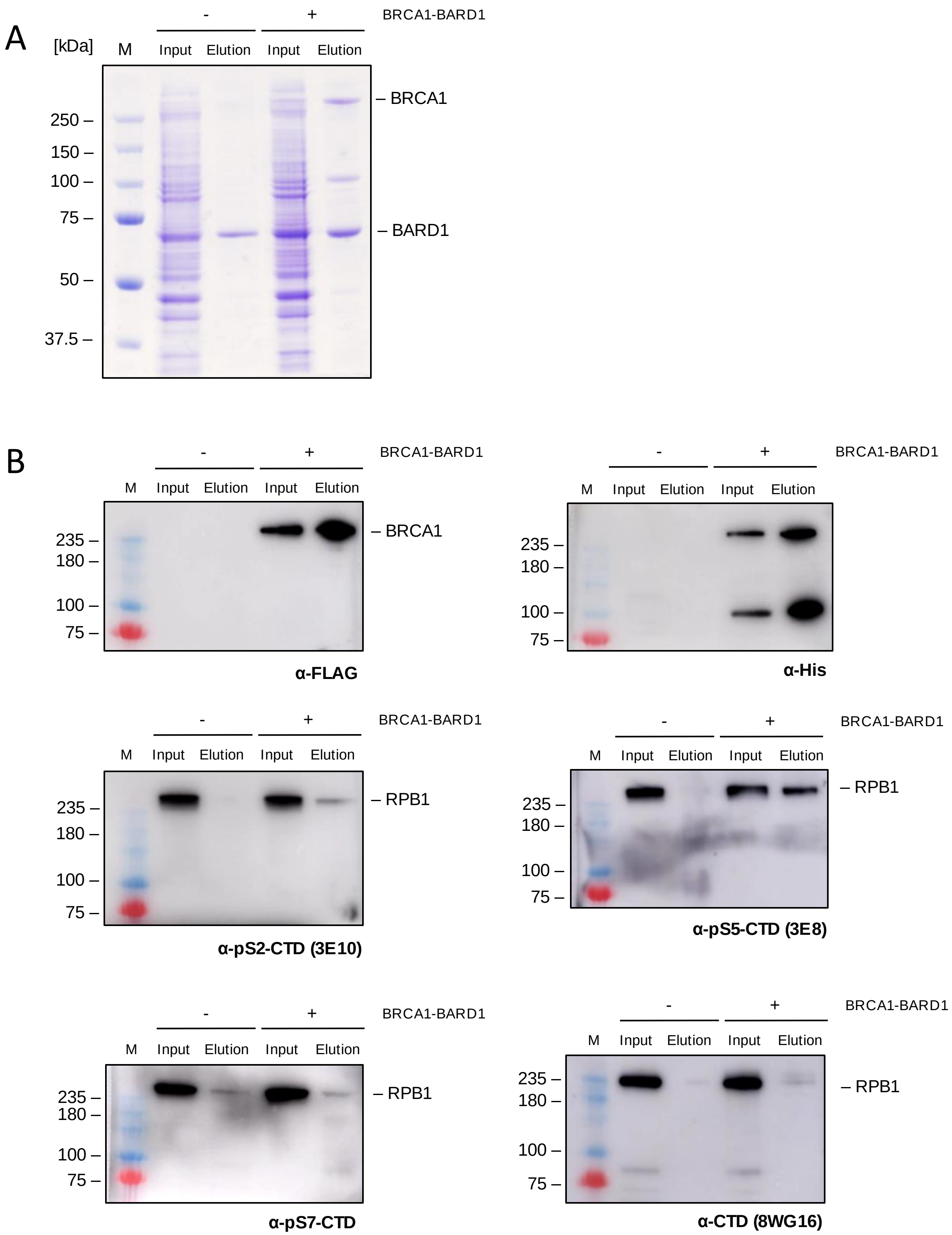

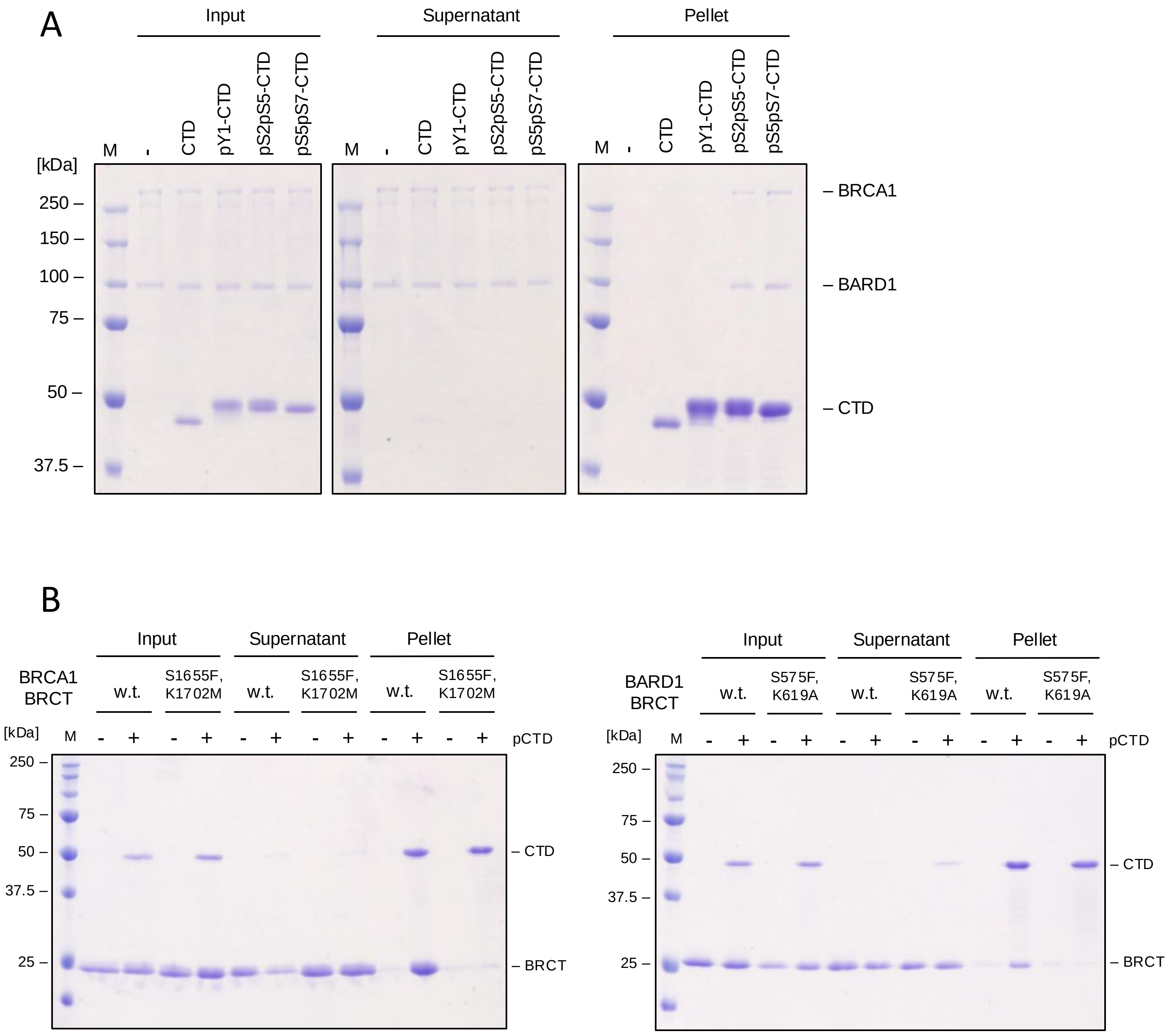
A
Input
Supernatant
Pellet
M
-
CTD
pY1-CTD
pS2pS5-CTD
pS5pS7-CTD
[kDa]
250 –
150 –
100 –
75 –
50 –
37.5 –
– BRCA1
– BARD1
– CTD
B
BRCA1 BRCT
w.t.
S1655F, K1702M
BARD1 BRCT
S575F, K619A
pCTD
– BRCT
25 –

**Supplementary figure 4** Klapstova *et al.*,

A BRCA1 BRCT

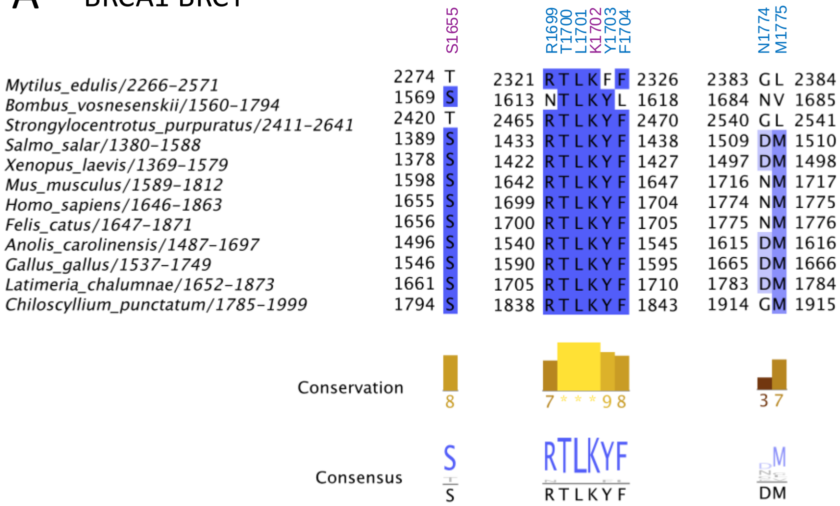


B BARD1 BRCT

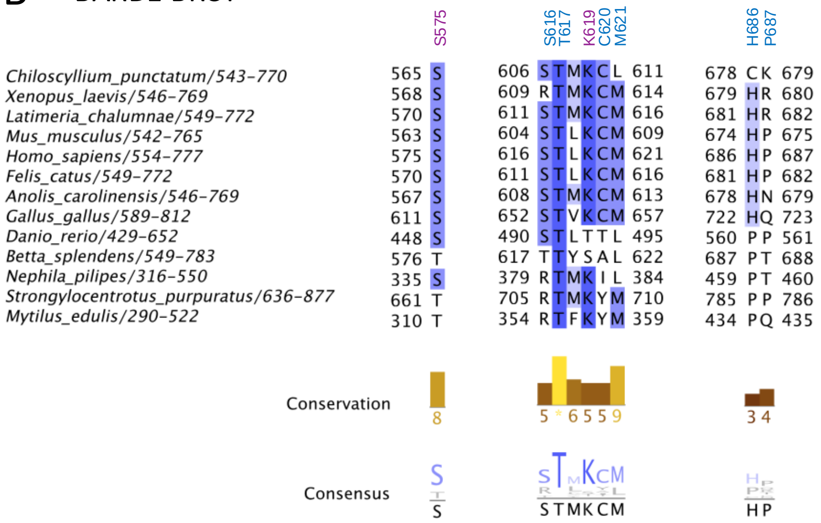

**Supplementary figure 5** **Klapstova *et al.*,**

A BRCA1 BRCT

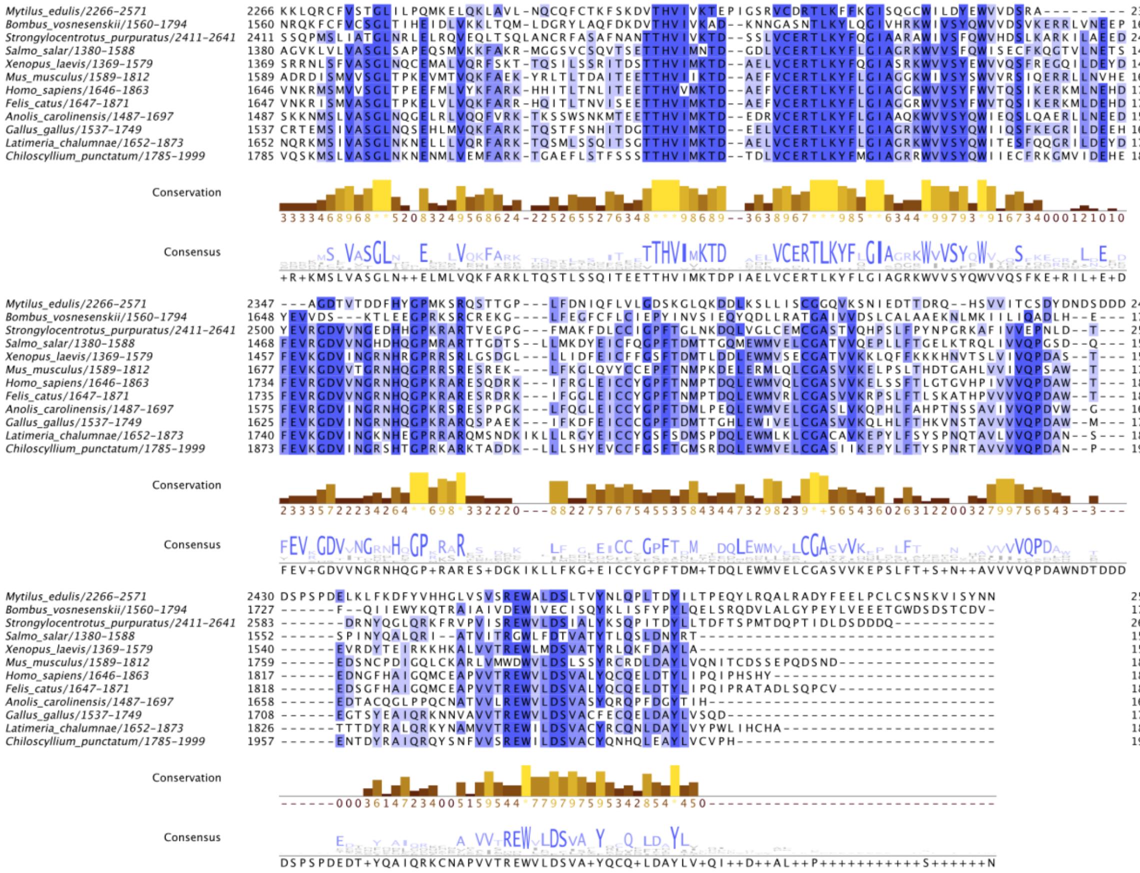


B BARD1 BRCT

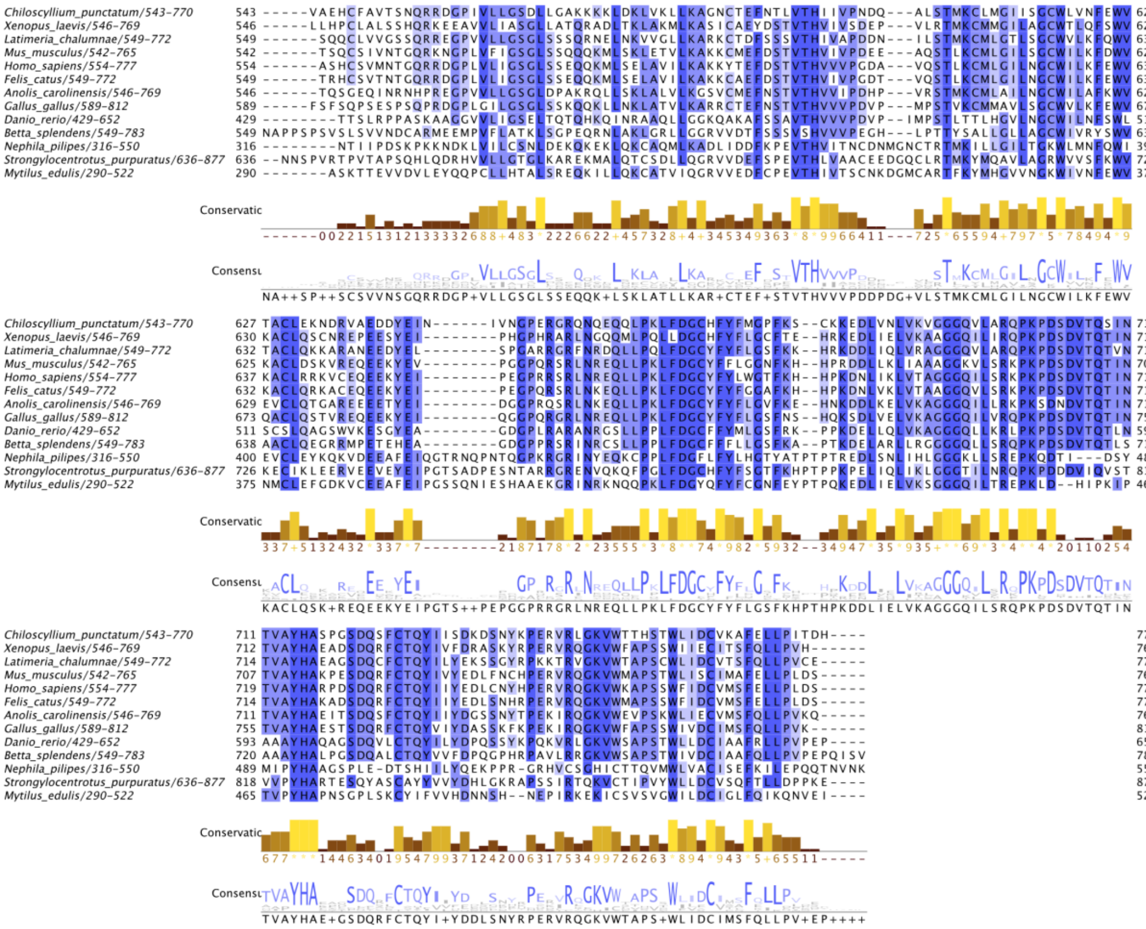

### Supplementary figure 6

Klapstova ***et al.***,

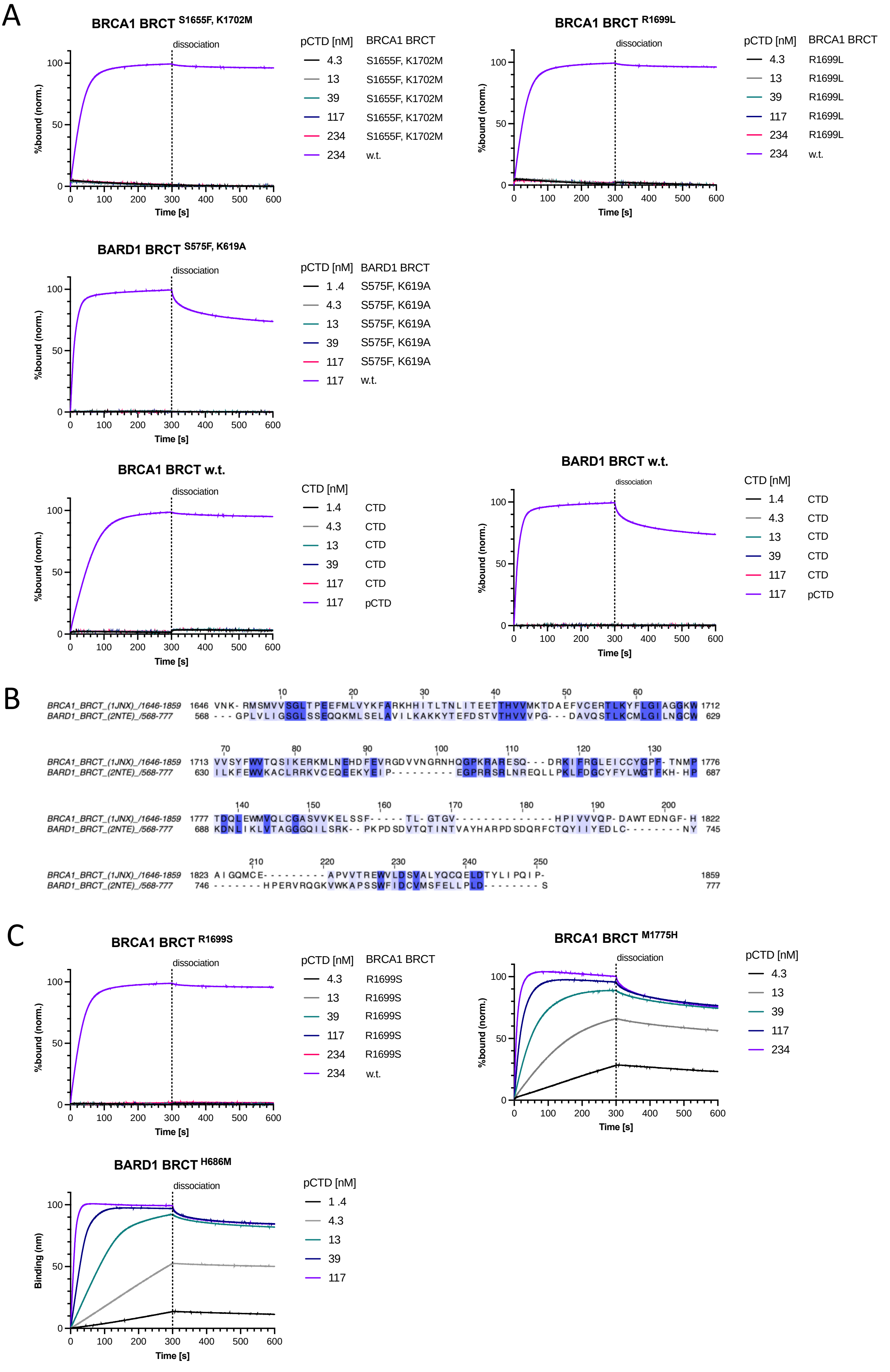

#### Supplementary figure 7

Klapstova *et al.*,

A

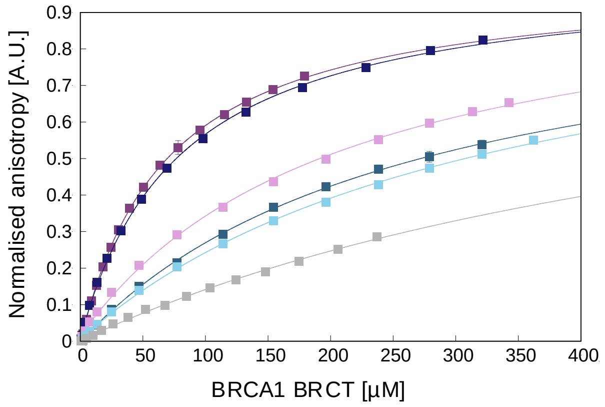


| CTD peptide | $K_D$ [µM] |
|---|---|
| pS2pS5pS7 | 68.2 ± 0.9 |
| pS5 | 78.4 ± 2.7 |
| pS2pS5 | 202.6 ± 14.3 |
| pS7 | 270.2 ± 12.2 |
| pS2 | 314.7 ± 16.6 |
| CTD | 618.7 ± 93.1 |

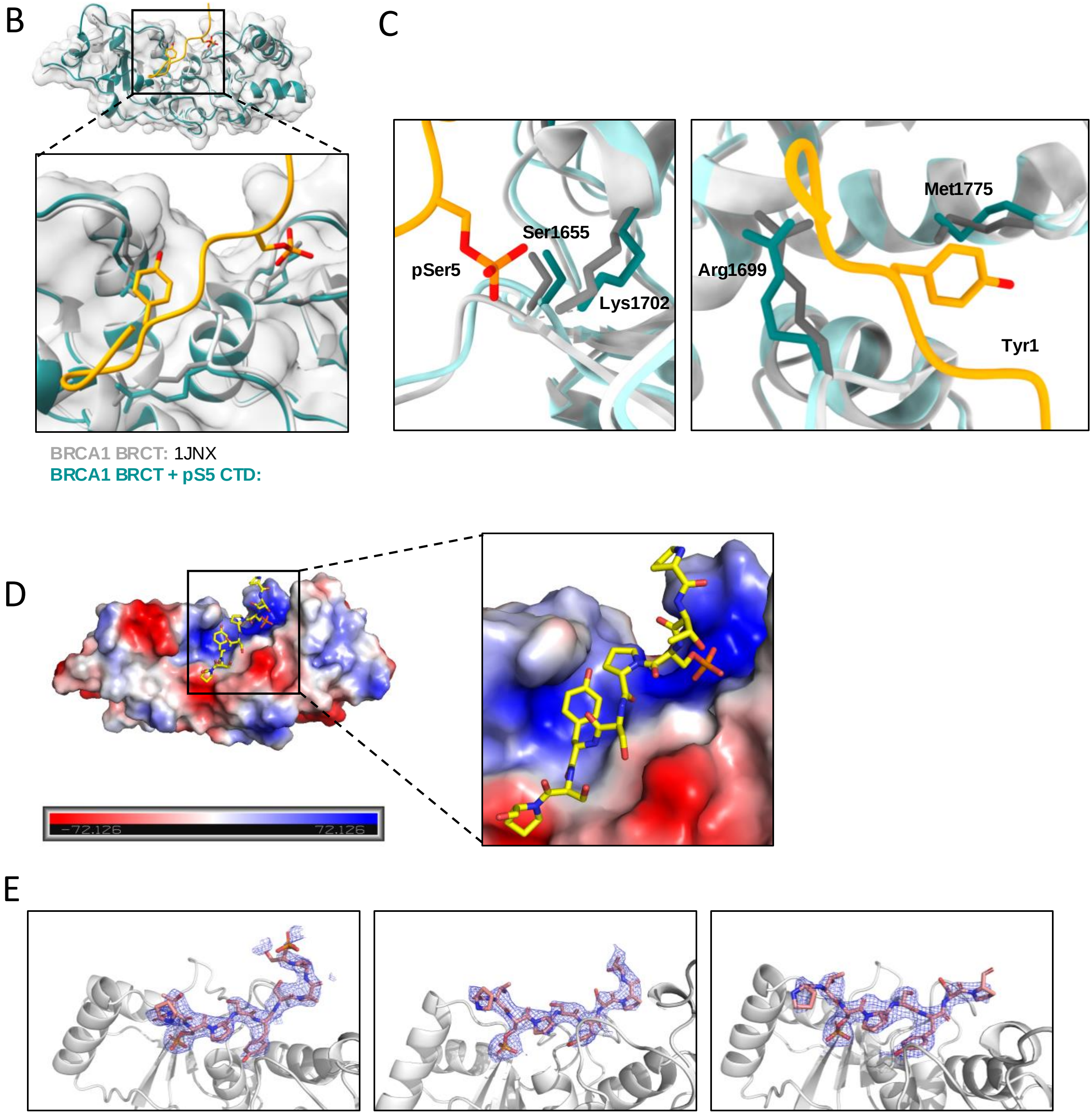

**Supplementary figure 8** Klapstova *et al.*,

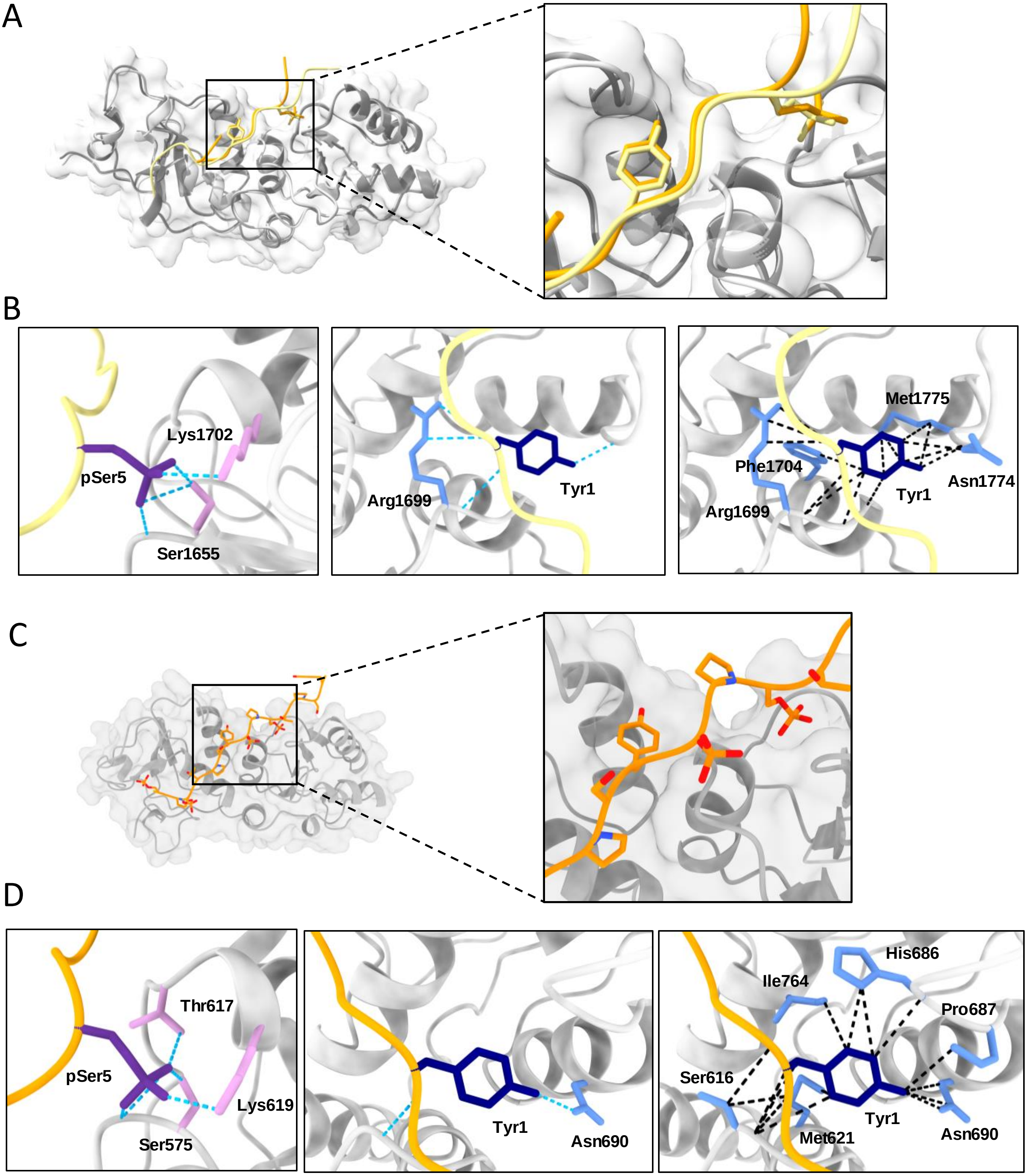

**Supplementary figure 9** **Klapstova *et al.*,**

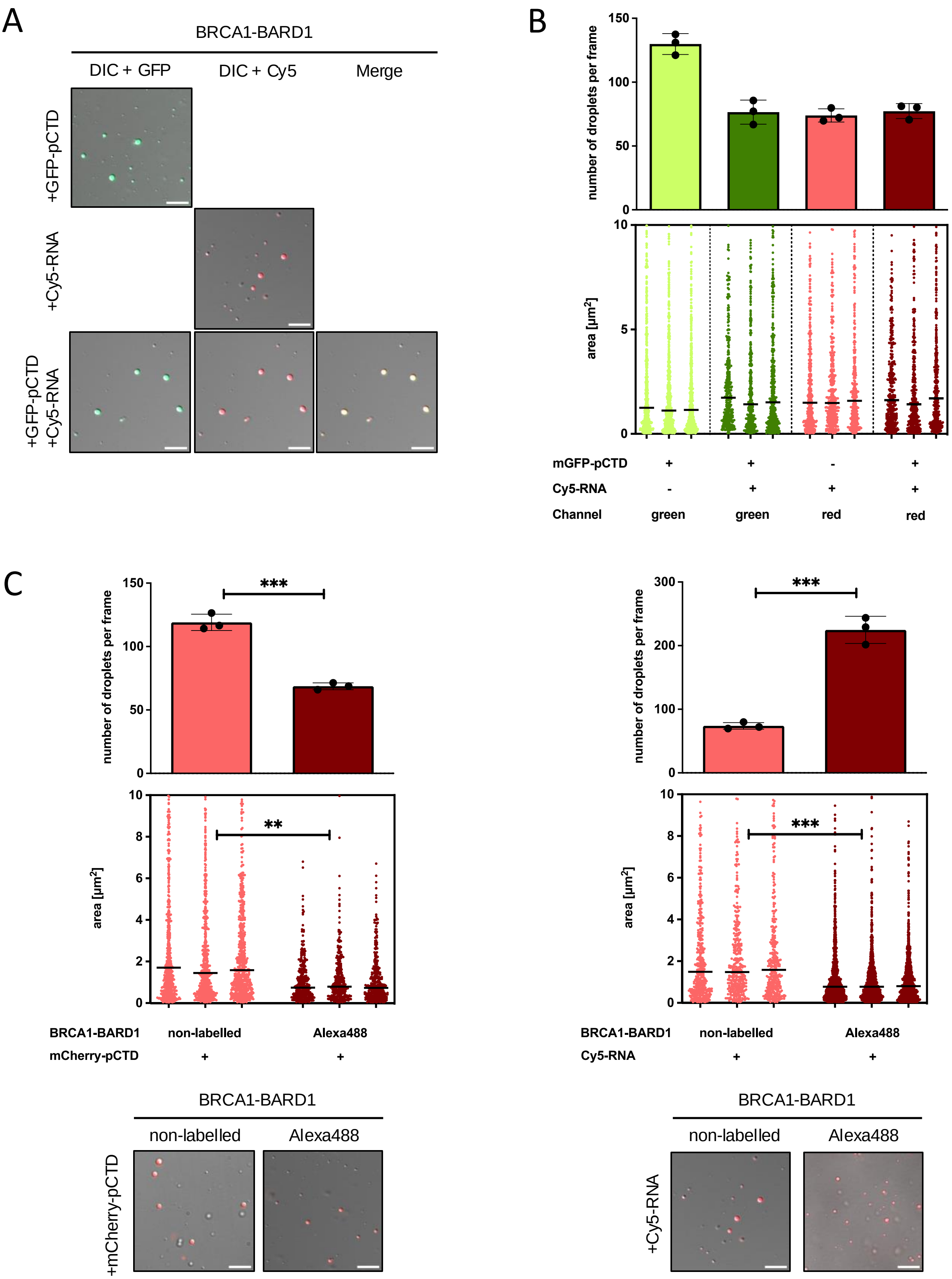

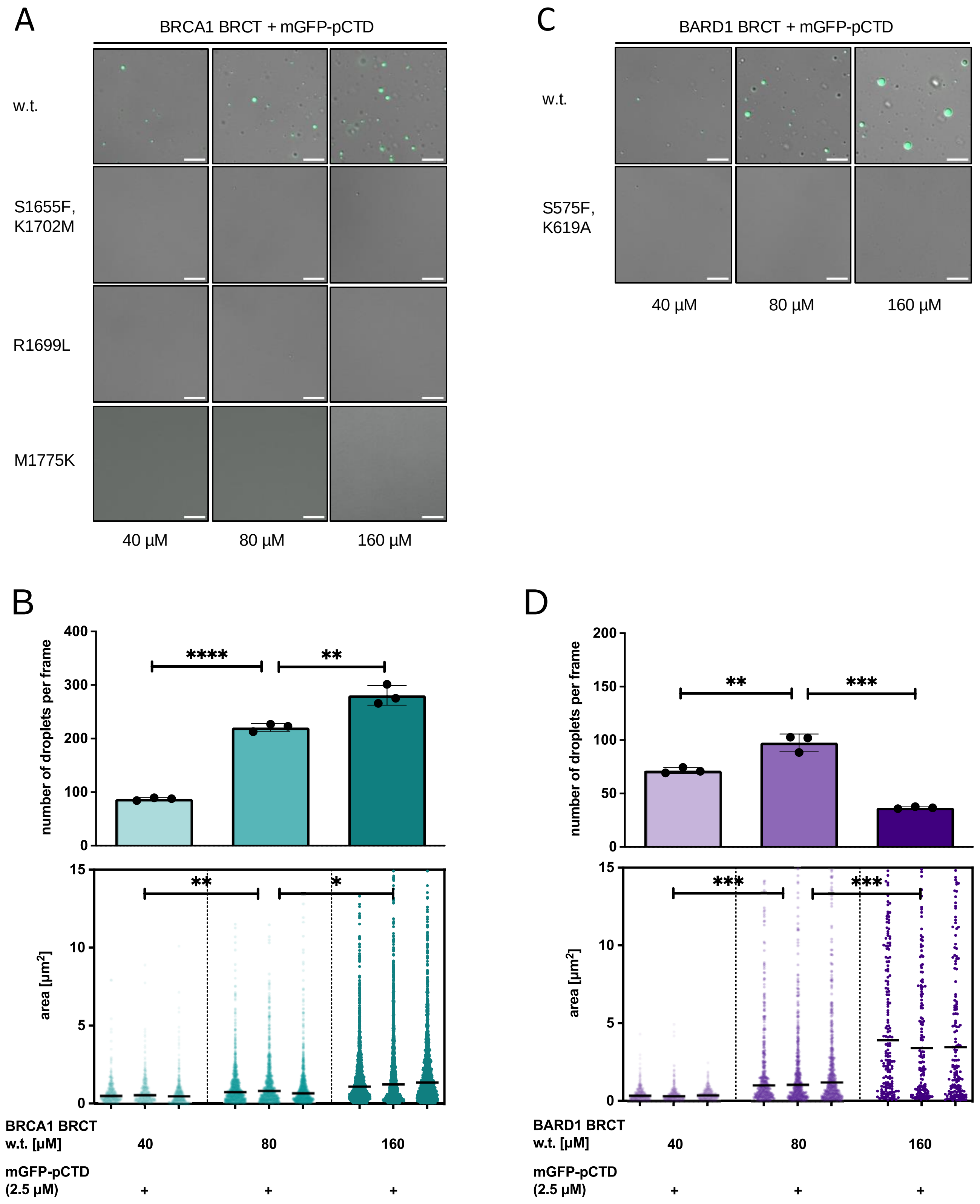
A
BRCA1 BRCT + mGFP-pCTD
w.t.
S1655F,
K1702M
R1699L
M1775K
40 µM
80 µM
160 µM
C
BARD1 BRCT + mGFP-pCTD
w.t.
S575F,
K619A
40 µM
80 µM
160 µM
B
number of droplets per frame
400
300
200
100
0
****
**
15
10
5
0
area [µm2]
**
*
BRCA1 BRCT
w.t. [µM]
40
80
160
mGFP-pCTD
(2.5 µM)
+
+
+
D
number of droplets per frame
200
150
100
50
0
**
***
15
10
5
0
area [µm2]
***
***
BARD1 BRCT
w.t. [µM]
40
80
160
mGFP-pCTD
(2.5 µM)
+
+
+

**Supplementary figure 11** **Klapstova *et al.*,**

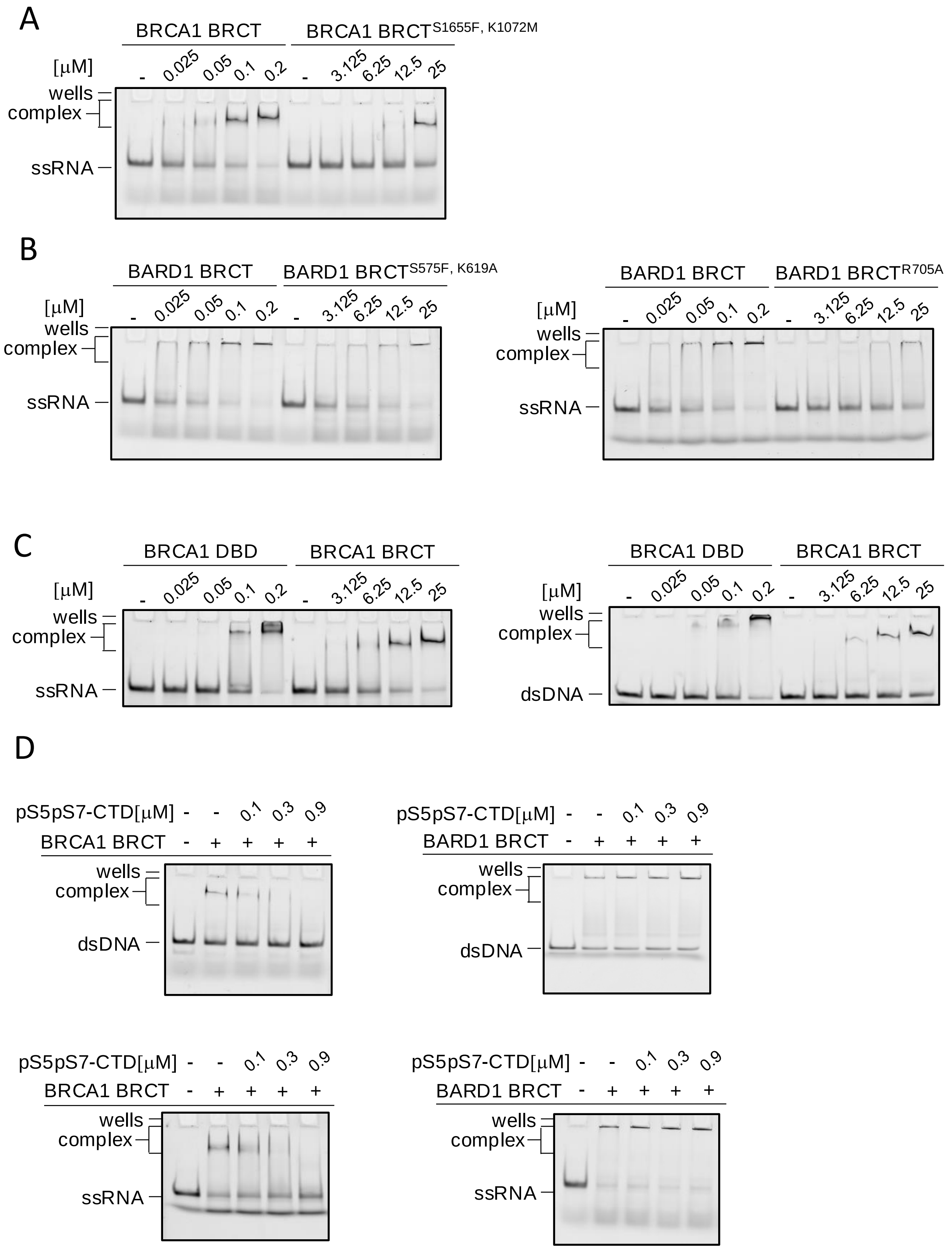

**Supplementary figure 12** **Klapstova *et al.*,**

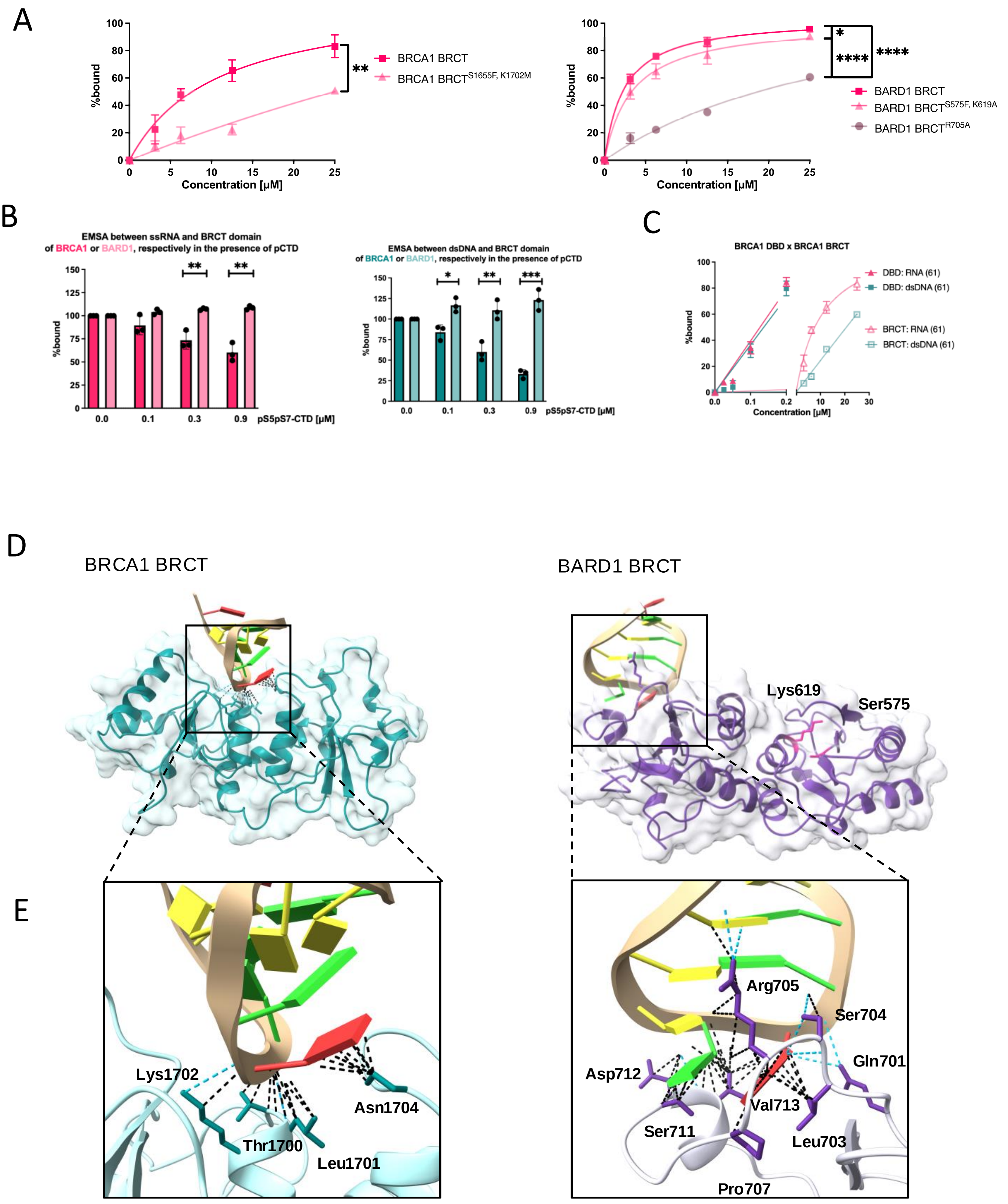

**Supplementary figure 13** **Klapstova *et al.*,**

A

BRCA1 BRCT BARD1 BRCT

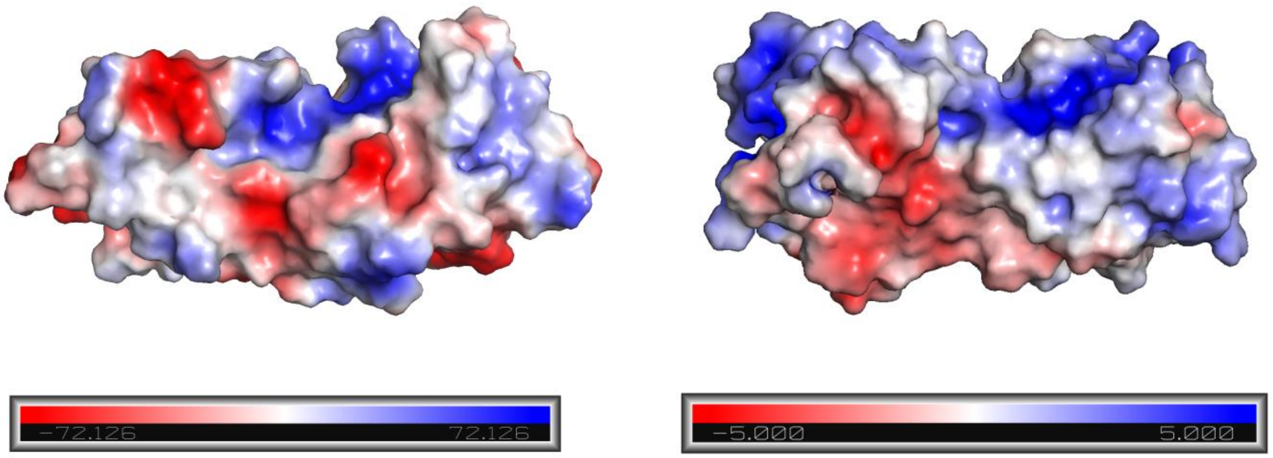


B

BARD1 BRCT

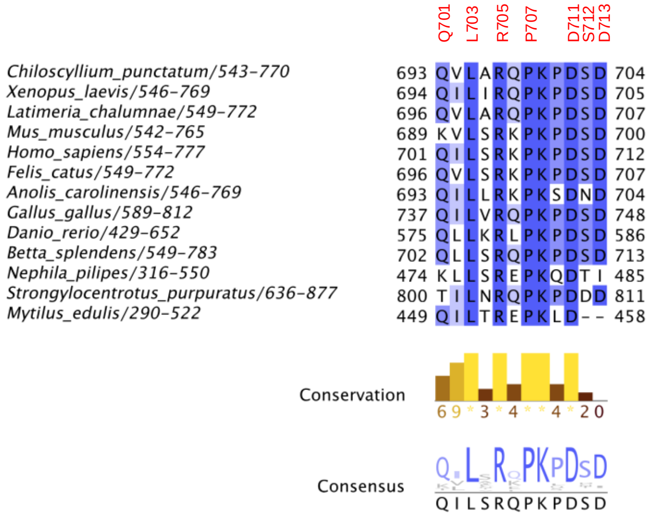

### Supplementary figure 14

Klapstova *et al.*,

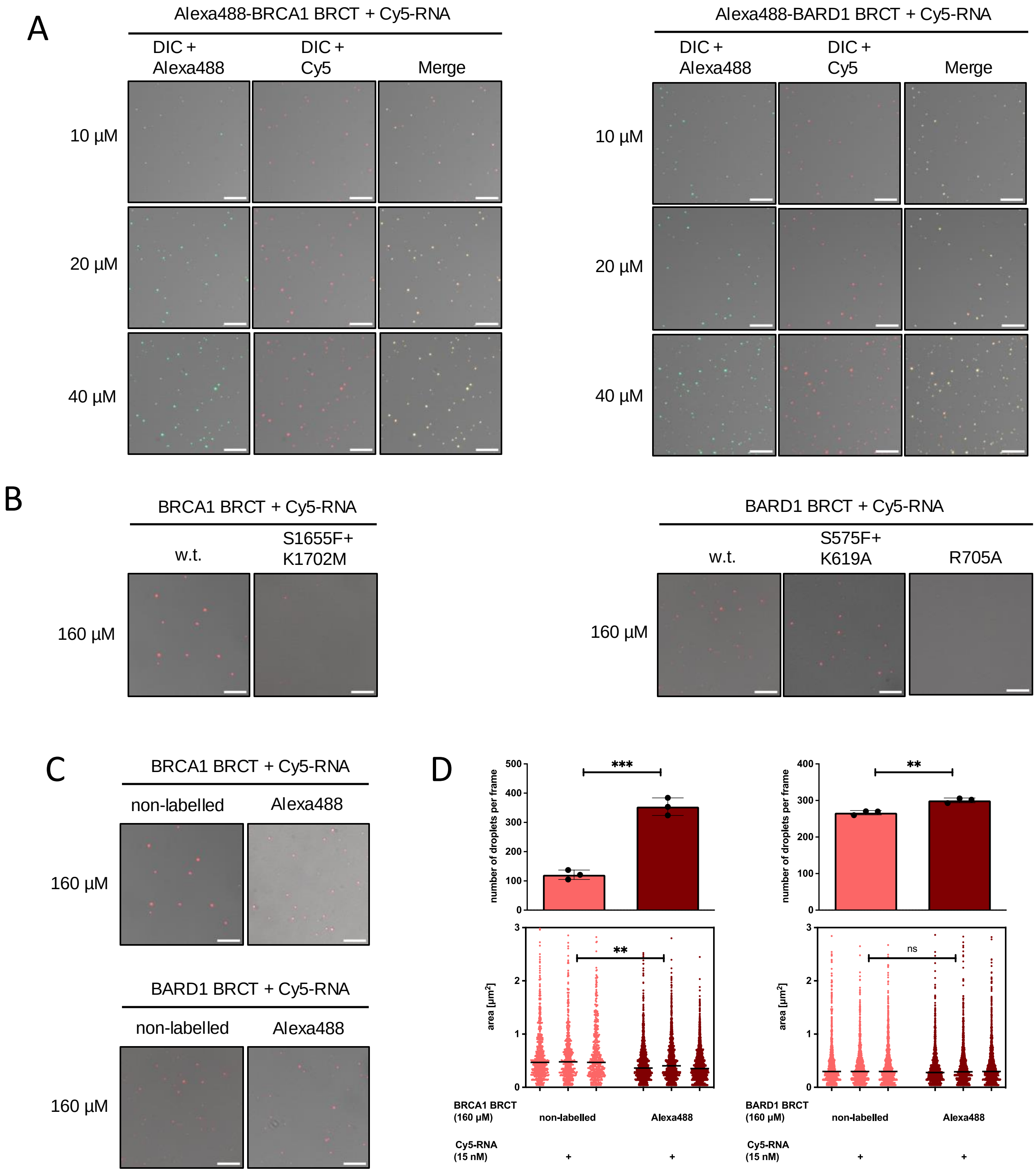

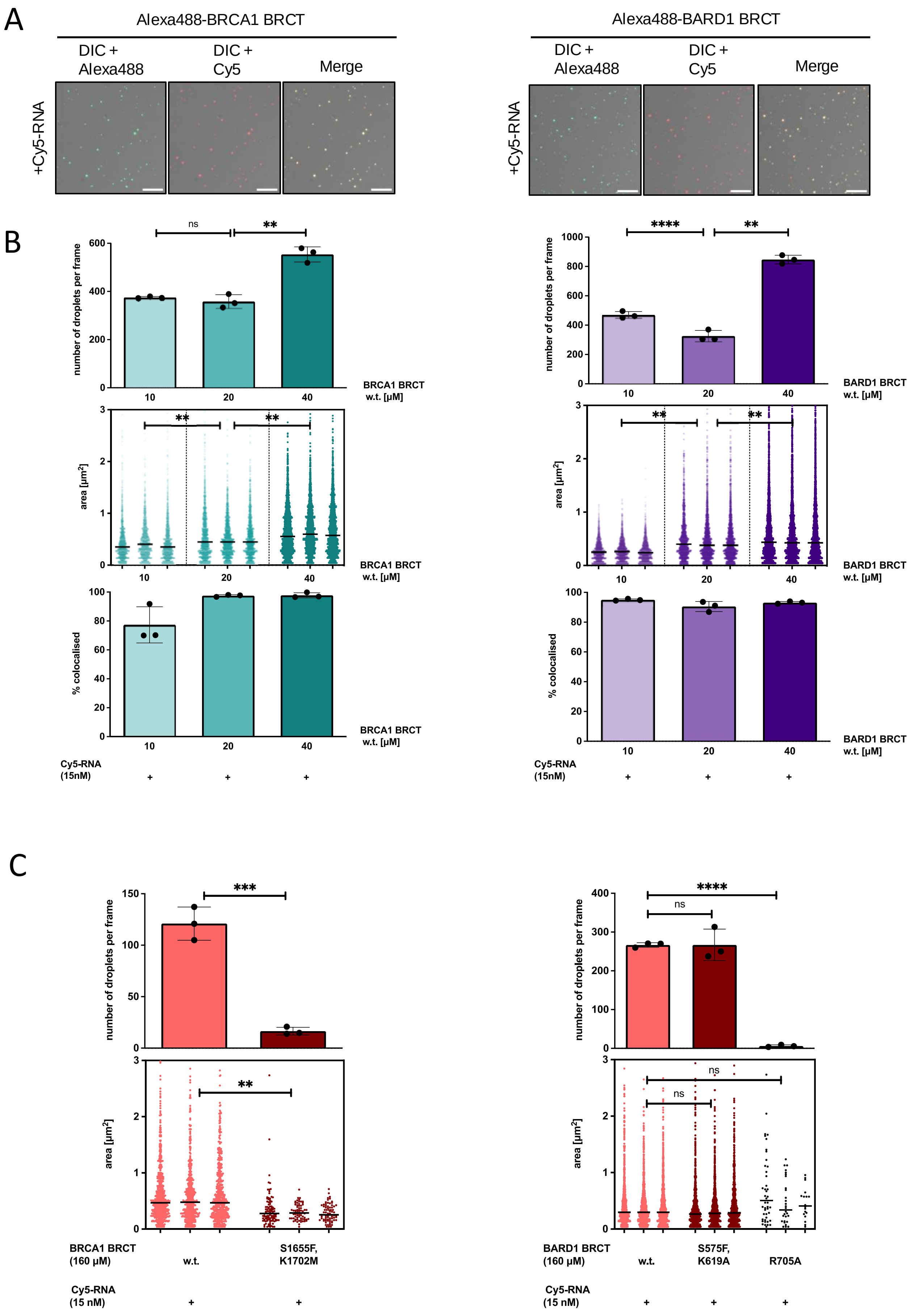
A
Alexa488-BRCA1 BRCT
DIC + Alexa488
DIC + Cy5
Merge
+Cy5-RNA
Alexa488-BARD1 BRCT
B
ns
**
number of droplets per frame
BRCA1 BRCT w.t. [µM]
10
20
40
****
BARD1 BRCT w.t. [µM]
area [µm2]
% colocalised
Cy5-RNA (15nM)
+
C
***
ns
BRCA1 BRCT (160 µM)
w.t.
S1655F, K1702M
Cy5-RNA (15 nM)
BARD1 BRCT (160 µM)
S575F, K619A
R705A

Supplementary figure 16 **Klapstova *et al.*,**

A

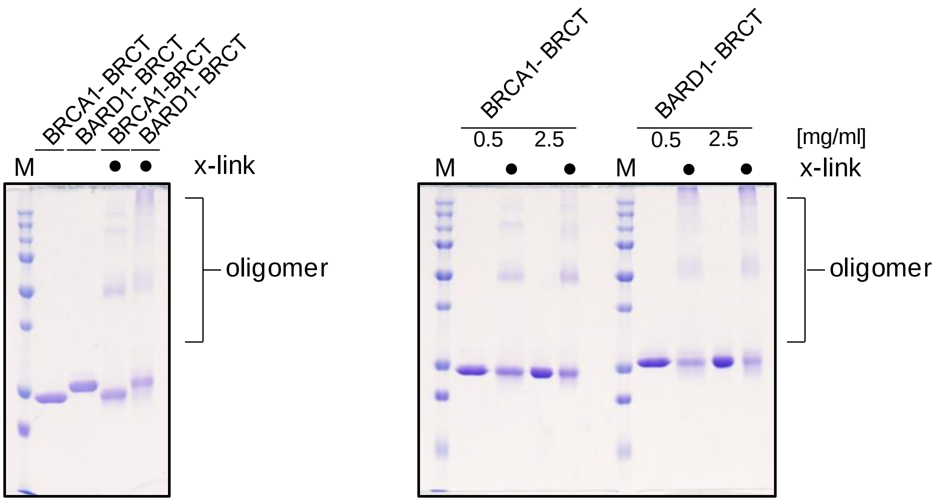


B

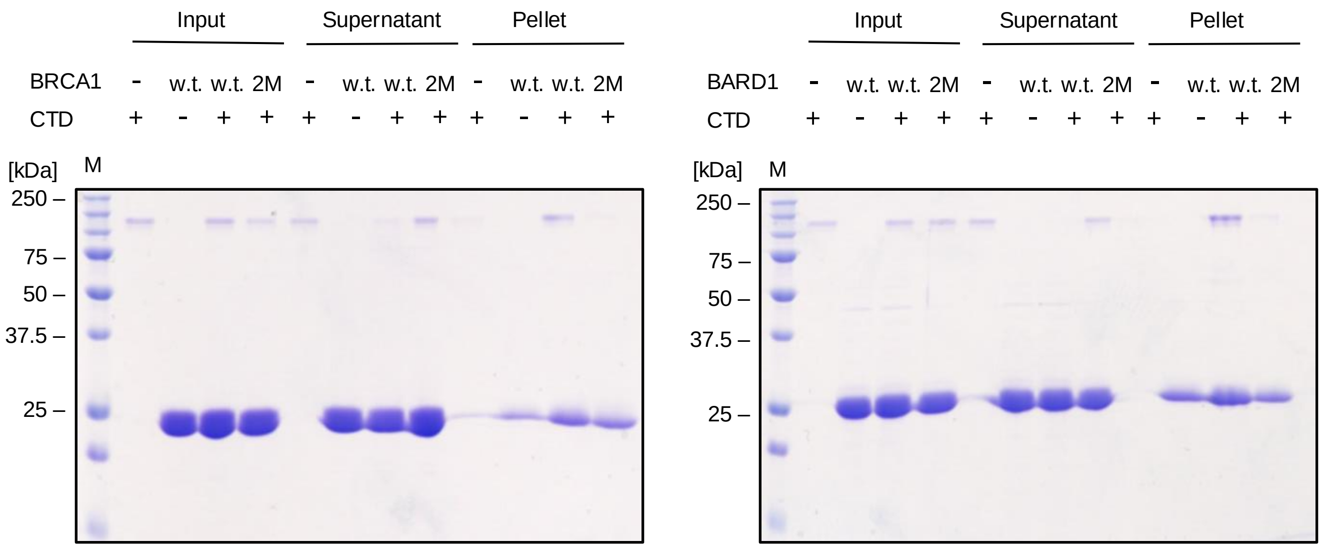


C

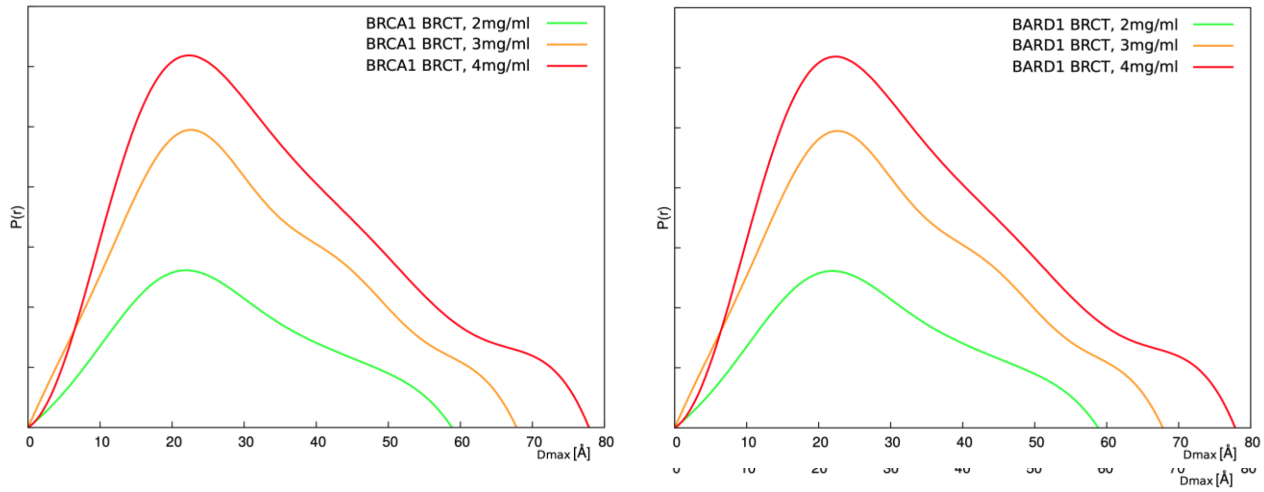

### Supplementary figure 17

Klapstova *et al.*,

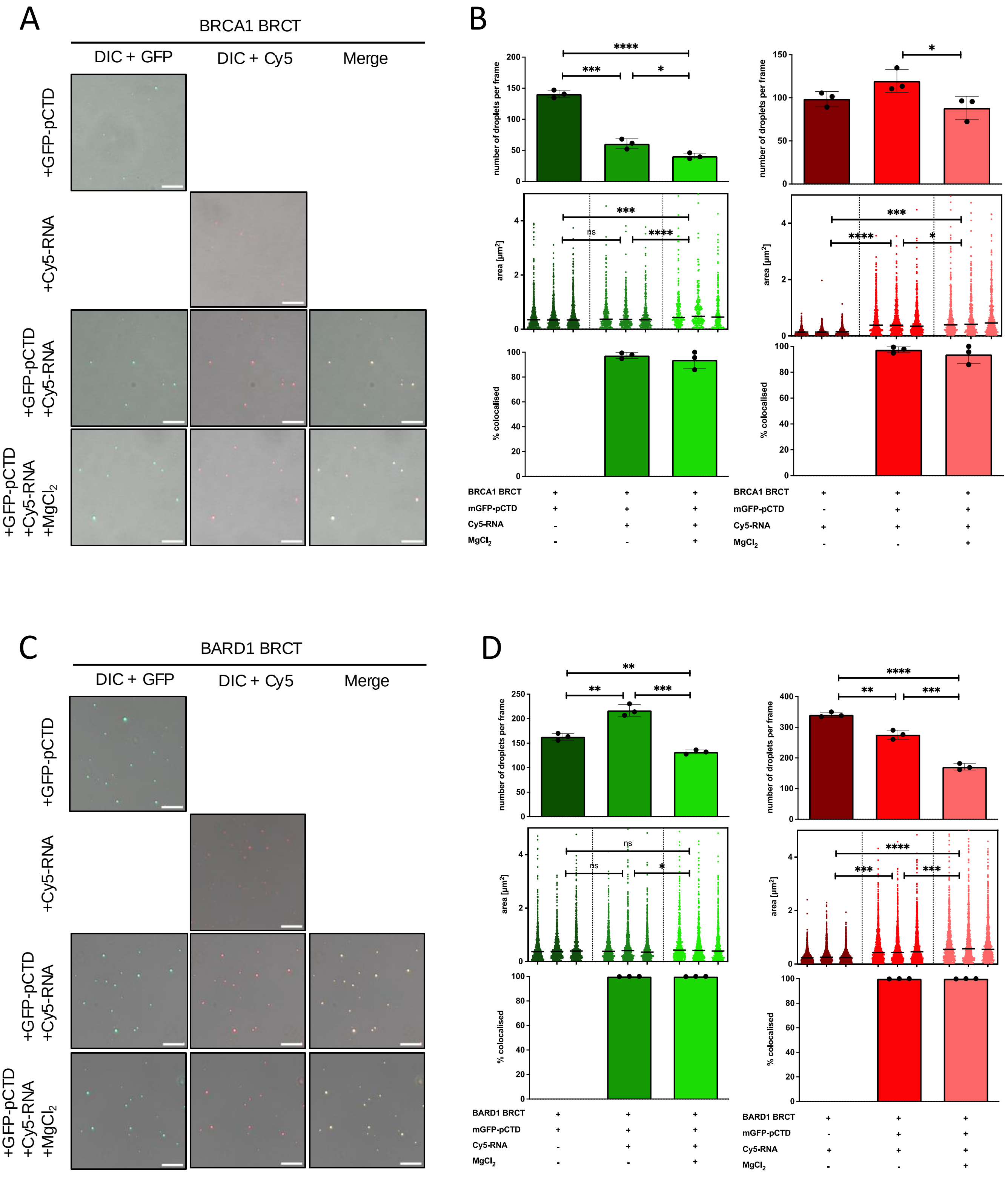

### Supplementary figure 18

Klapstova *et al.*,

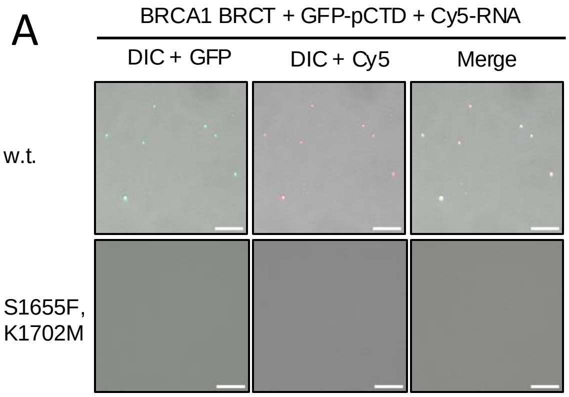


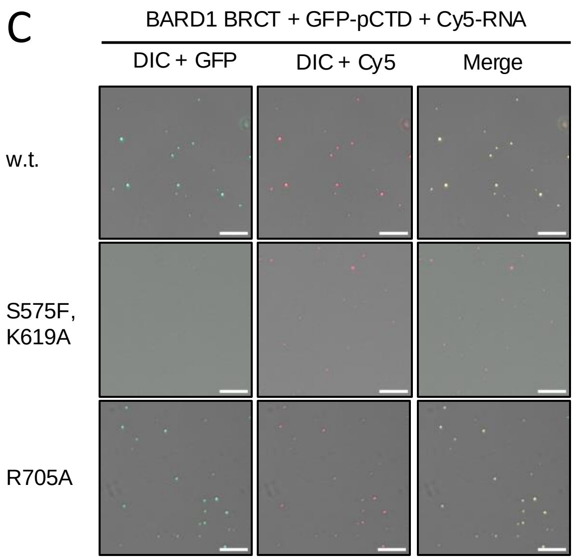


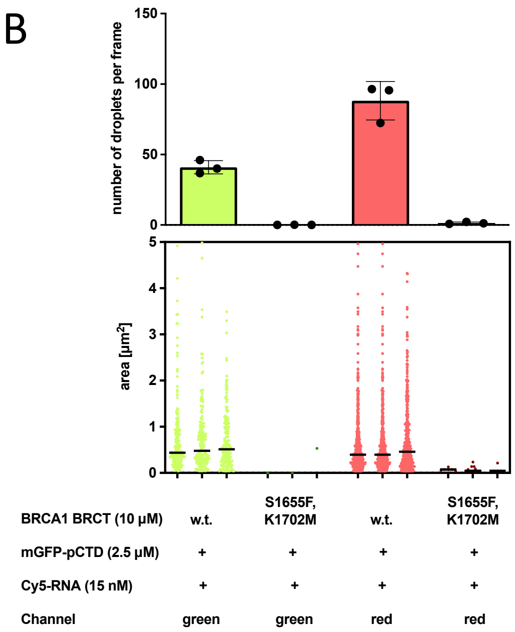


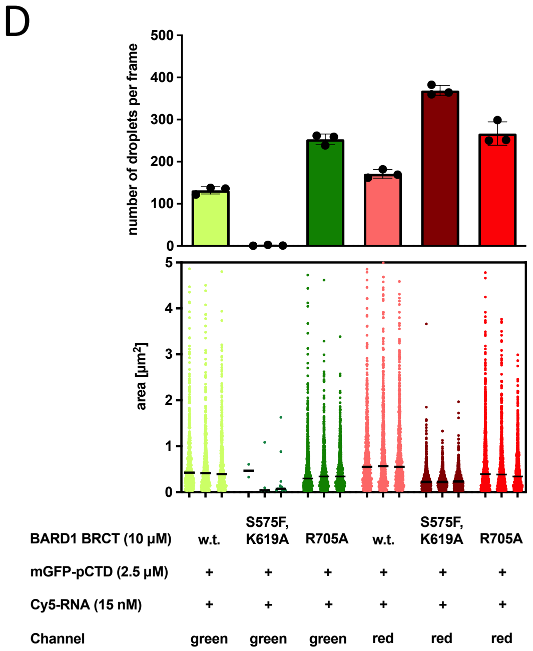


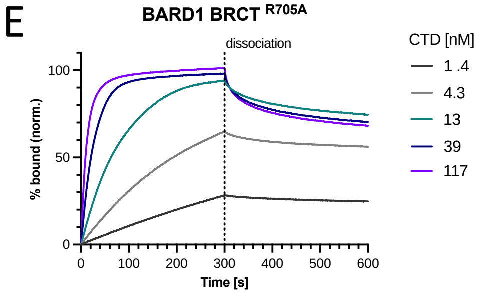


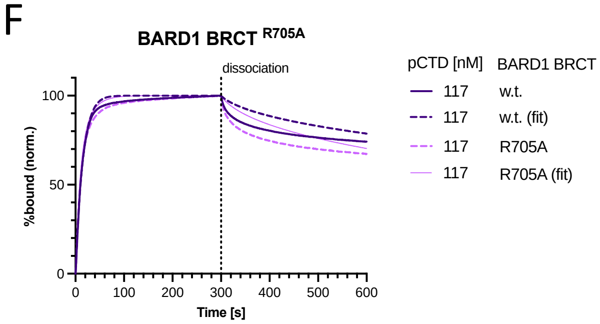


| BARD1 BRCT | $k_D$ (M) | $k_D$ error | Full $R^2$ | $K_{as}$ (1/Ms) | $K_{dis}$ (1/s) |
|---|---|---|---|---|---|
| w.t. | 1,93E$^{-07}$ | 2,88E$^{-08}$ | 0,9730 | 2,66E$^{+05}$ | 5,12E$^{-02}$ |
| R705A | 3,05E$^{-07}$ | 3,07E$^{-08}$ | 0,9893 | 2,17E$^{+05}$ | 6,63E$^{-02}$ |

#### A BRCA1 BRCT

| | | E1682 | | E1754 | S1755 |
|---|---|---|---|---|---|
| *Mytilus_edulis/2266-2571* | 2302 | K | 2364 | Q | S |
| *Bombus_vosnesenskii/1560-1794* | 1596 | K | 1665 | C | R |
| *Strongylocentrotus_purpuratus/2411-2641* | 2448 | A | 2520 | T | V |
| *Salmo_salar/1380-1588* | 1416 | S | 1488 | T | T |
| *Xenopus_laevis/1369-1579* | 1405 | D | 1477 | L | G |
| *Mus_musculus/1589-1812* | 1625 | E | 1697 | E | S |
| *Homo_sapiens/1646-1863* | 1682 | E | 1754 | E | S |
| *Felis_catus/1647-1871* | 1683 | E | 1755 | E | S |
| *Anolis_carolinensis/1487-1697* | 1523 | E | 1595 | E | S |
| *Gallus_gallus/1537-1749* | 1573 | D | 1645 | Q | S |
| *Latimeria_chalumnae/1652-1873* | 1688 | S | 1760 | Q | M |
| *Chiloscyllium_punctatum/1785-1999* | 1821 | S | 1893 | K | T |

#### B BARD1 BRCT

| | | E587 | | E665 | | S711 | | K754 |
|---|---|---|---|---|---|---|---|---|
| *Chiloscyllium_punctatum/543-770* | 577 | K | 657 | E | 703 | S | 746 | K |
| *Xenopus_laevis/546-769* | 580 | K | 658 | Q | 704 | S | 747 | K |
| *Latimeria_chalumnae/549-772* | 582 | K | 660 | D | 706 | S | 749 | K |
| *Mus_musculus/542-765* | 575 | K | 653 | E | 699 | S | 742 | K |
| *Homo_sapiens/554-777* | 587 | E | 665 | E | 711 | S | 754 | K |
| *Felis_catus/549-772* | 582 | E | 660 | E | 706 | S | 749 | K |
| *Anolis_carolinensis/546-769* | 579 | K | 657 | E | 703 | N | 746 | K |
| *Gallus_gallus/589-812* | 623 | K | 701 | E | 747 | S | 790 | K |
| *Danio_rerio/429-652* | 460 | R | 539 | G | 585 | S | 628 | K |
| *Betta_splendens/549-783* | 588 | K | 666 | C | 712 | S | 755 | K |
| *Nephila_pilipes/316-550* | 347 | K | 436 | E | 484 | T | 522 | H |
| *Strongylocentrotus_purpuratus/636-877* | 673 | T | 762 | Q | 810 | D | 853 | K |
| *Mytilus_edulis/290-522* | 322 | K | 411 | K | | - | 498 | K |

Supplementary figure 20 Klapstova *et al.*,

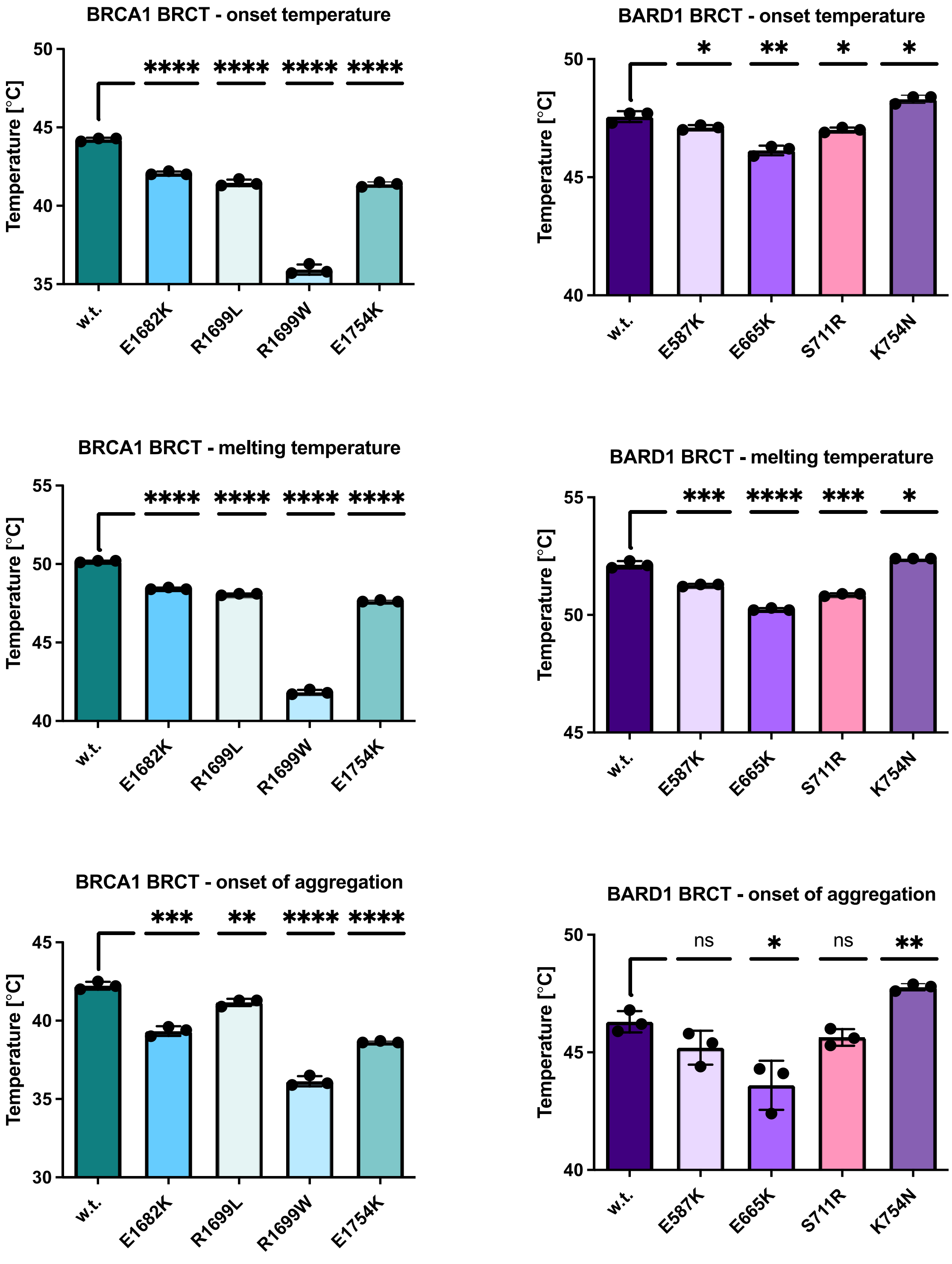

A

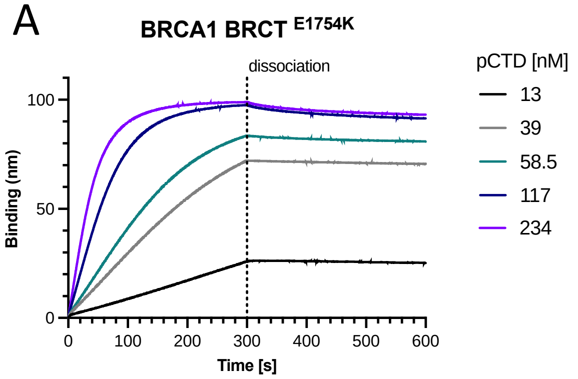


B

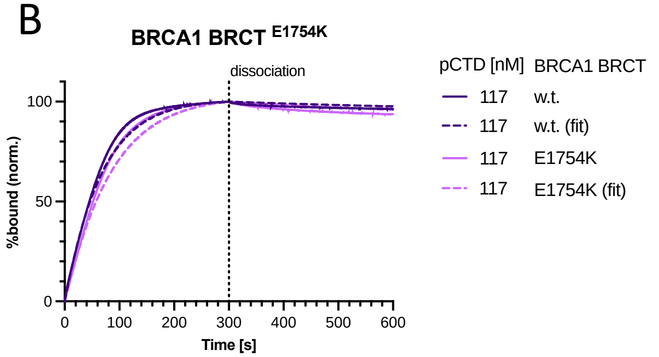


| BRCA1 BRCT | $k_D$ (M) | $k_D$ error | Full $R^2$ | $K_{as}$ (1/Ms) | $K_{dis}$ (1/s) |
|---|---|---|---|---|---|
| **w.t.** | 1,98E-08 | 9,60E-10 | 0,9982 | 3,22E+04 | 6,37E-04 |
| **E1754K** | 3,05E-08 | 1,52E-09 | 0,9941 | 2,60E+04 | 7,92E-04 |

C

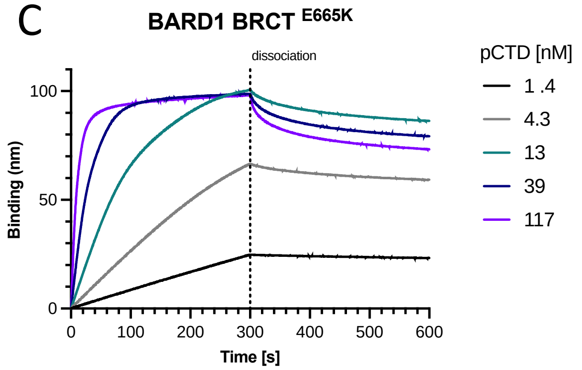


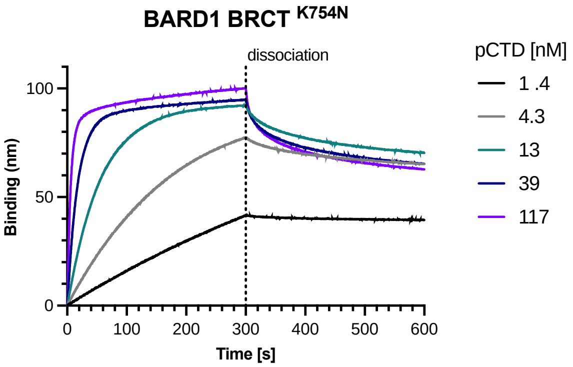


D

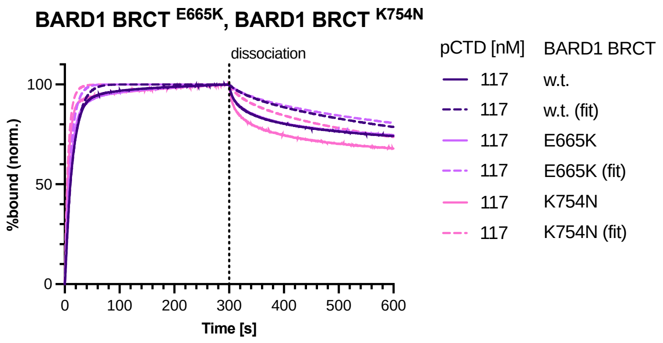


| BARD1 BRCT | $k_D$ (M) | $k_D$ error | Full $R^2$ | $K_{as}$ (1/Ms) | $K_{dis}$ (1/s) |
|---|---|---|---|---|---|
| **w.t.** | 1,93E-07 | 2,88E-08 | 0,9730 | 2,66E+05 | 5,12E-02 |
| **E665K** | 3,28E-07 | 8,26E-08 | 0,9819 | 2,07E+05 | 6,79E-02 |
| **K754N** | 1,11E-07 | 9,54E-09 | 0,9693 | 3,64E+05 | 4,03E-02 |

**Supplementary figure 22** Klapstova *et al.*,

A

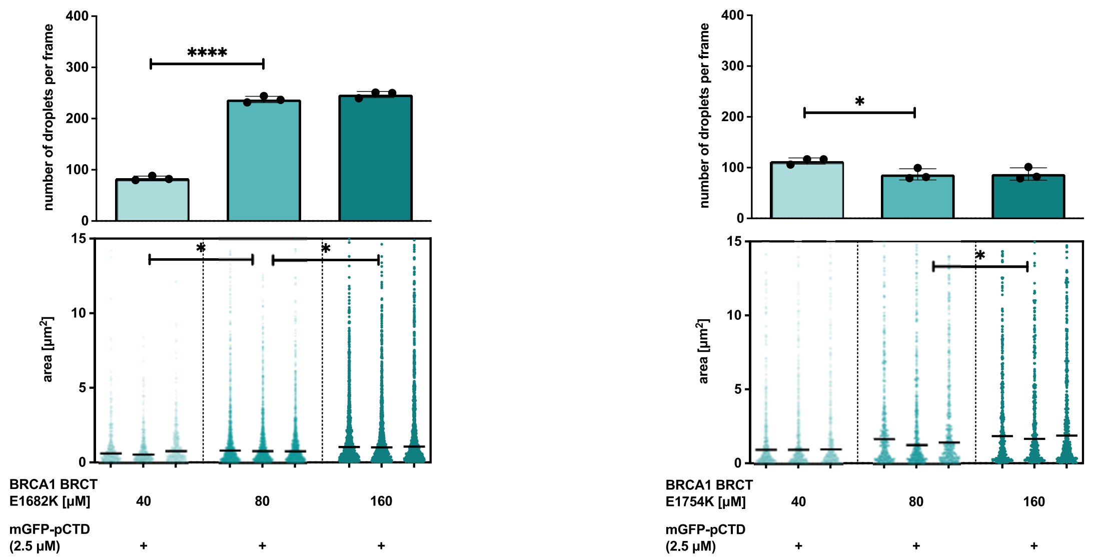


B

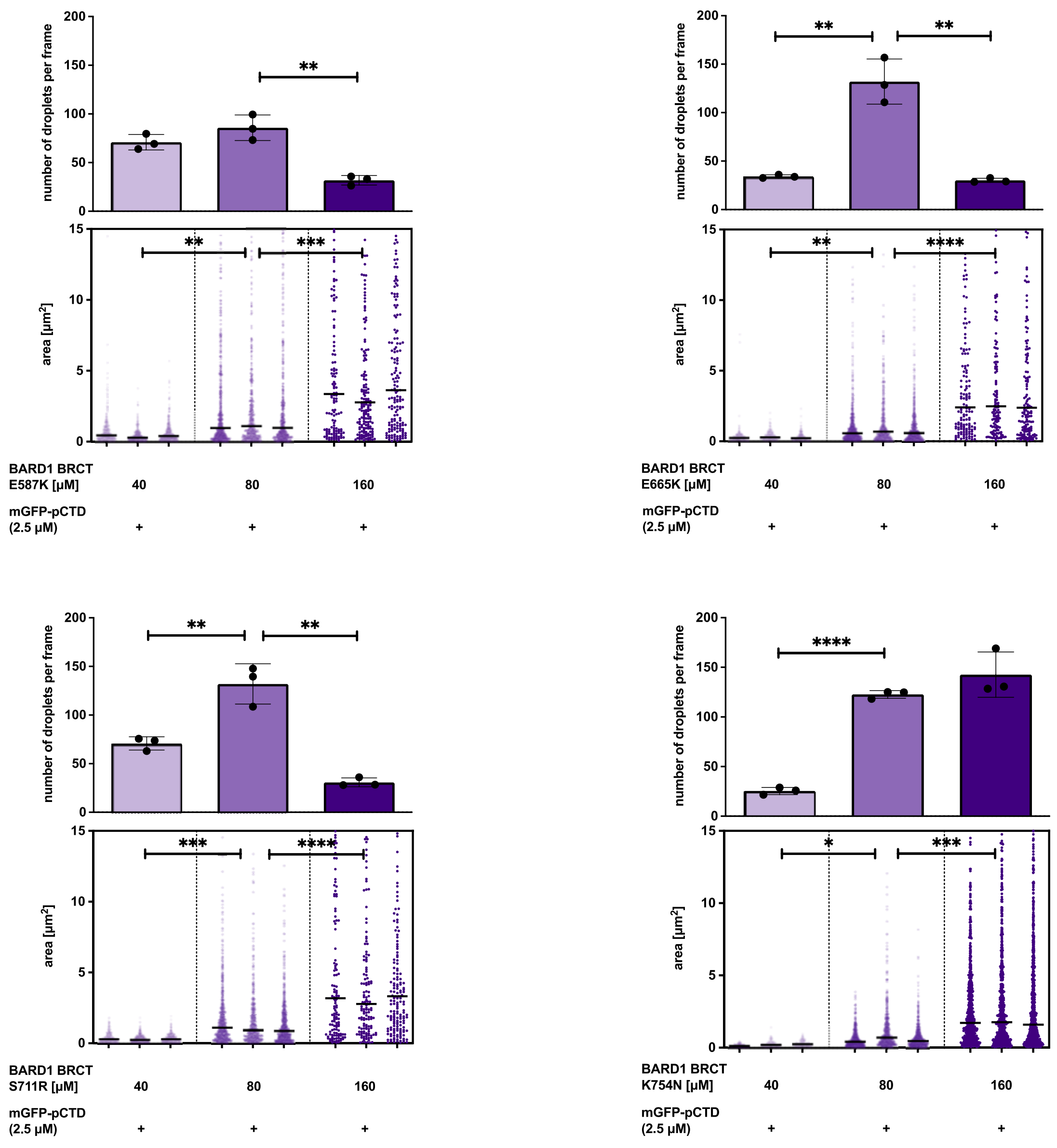

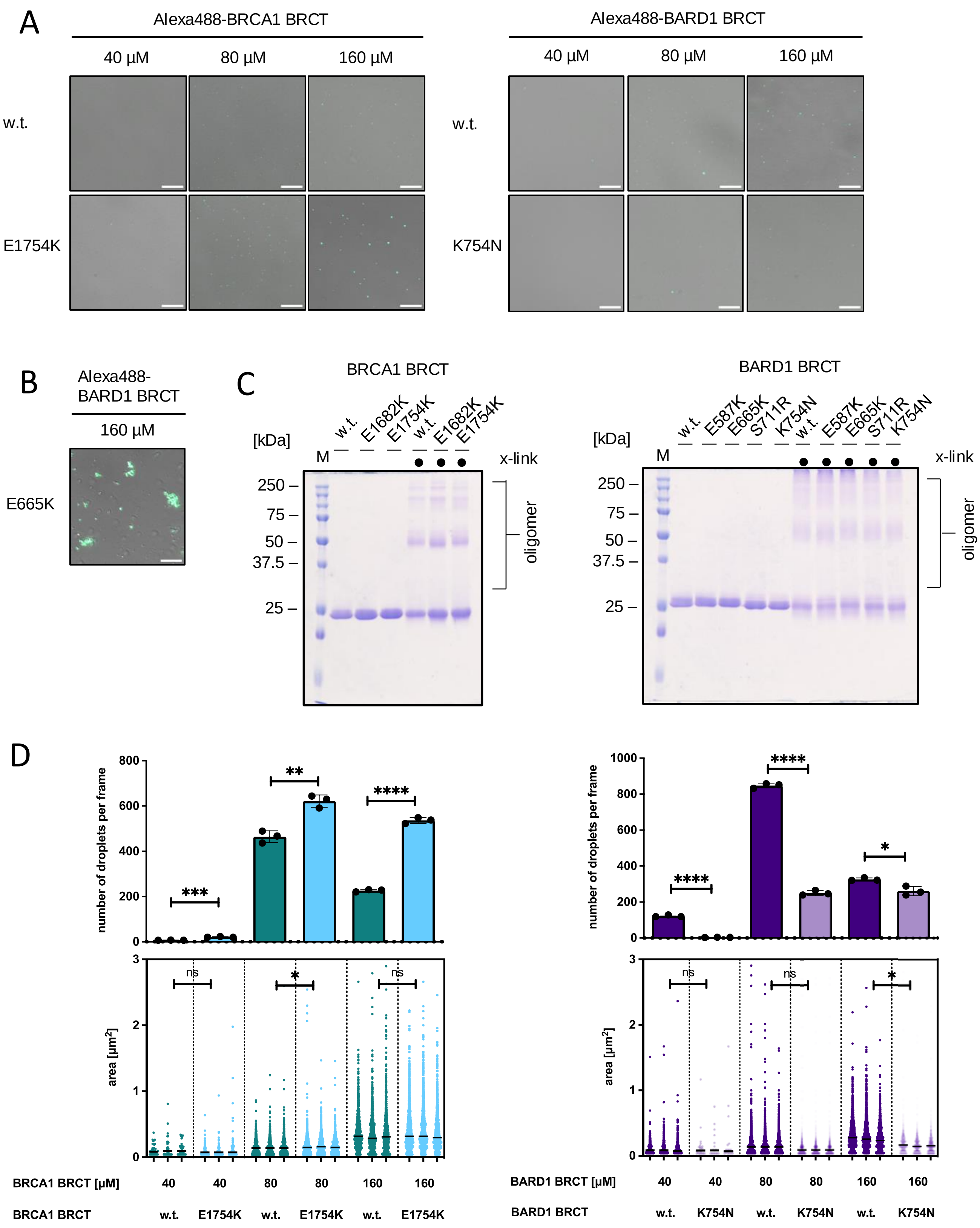
A
Alexa488-BRCA1 BRCT
40 µM
80 µM
160 µM
w.t.
E1754K
Alexa488-BARD1 BRCT
40 µM
80 µM
160 µM
w.t.
K754N
B
Alexa488-BARD1 BRCT
160 µM
E665K
C
BRCA1 BRCT
[kDa]
M
w.t.
E1682K
E1754K
w.t.
E1682K
E1754K
x-link
250
75
50
37.5
25
oligomer
BARD1 BRCT
[kDa]
M
w.t.
E587K
E665K
S711R
K754N
w.t.
E587K
E665K
S711R
K754N
x-link
250
75
50
37.5
25
oligomer
D
number of droplets per frame
800
600
400
200
0
***
**
****
area [µm²]
3
2
1
0
ns
*
ns
BRCA1 BRCT [µM]
40
40
80
80
160
160
BRCA1 BRCT
w.t.
E1754K
w.t.
E1754K
w.t.
E1754K
number of droplets per frame
1000
800
600
400
200
0
****
****
*
area [µm²]
3
2
1
0
ns
ns
*
BARD1 BRCT [µM]
40
40
80
80
160
160
BARD1 BRCT
w.t.
K754N
w.t.
K754N
w.t.
K754N
